# From Functional Architecture to Organizing Principles of Neuronal Ensembles in Mouse Area V1

**DOI:** 10.1101/2024.02.29.582364

**Authors:** Maria Papadopouli, Ioannis Smyrnakis, Emmanouil Koniotakis, Mario Alexios Savaglio, Christina Brozi, Eleftheria Psilou, Ganna Palagina, Stelios M. Smirnakis

**Author notes:** Corresponding author: Maria Papadopouli.

## Abstract

While single-neuron responses in mouse V1 are well characterized, less is known about how functional ensembles— groups of neurons that co-activate more frequently than expected by chance—emerge as computational units within laminar V1 circuits. Even with increasingly detailed knowledge of structural connectivity, the rules governing ensemble organization and interactions remain unclear. We imaged pyramidal neurons across granular (L4) and supragranular (L2/3) layers of mouse V1 and applied pairwise functional connectivity analysis to identify multi-neuronal ensembles as putative information-processing modules. In the absence of visual stimulation, 19–34% of pyramidal pairs within 300*µ*m were functionally connected, declining to 10% at 1 mm. Layer 2 to 4 laminar networks exhibited a small-world architecture, L4 displaying slightly denser connectivity and a near-uniform degree-of-connectivity distribution. We propose that neurons together with their first-order functionally connected (1FC) partners constitute putative elementary units of cortical computation. The firing probability of layer 2/3 neurons exhibits a ReLU-like nonlinearity, emerging when ≥ 13% of L4-1FC “putative inputs” co-fire, yielding sparse yet reliable responses. Moreover, L2/3 neuronal responses depend on the count (N), not the identity, of co-active L4-1FC partners, with response sensitivity scaling as a power law in N. These properties persist during visual stimulation and across different states of alertness. Interestingly, L2/3 neurons with L4-1FC modules of different sizes exhibit distinct coupling to brain-state and different computational signatures. This framework yields mechanistic insight into cortical circuit organization, complementary to structural connectivity, helping to link biological circuitry to deep-learning models of artificial intelligence.

## 1. Introduction

Athough much is known about the properties of individual neurons, we still lack a clear understanding of how cortical networks coordinate their activity to transmit and transform information across layers. Neuronal ensembles—groups of neurons that co-activate more frequently than expected by chance—are thought to operate as basic units of cortical computation, efficiently relaying shared information to downstream targets. This view is supported by pioneering studies describing structured patterns of spontaneous and stimulus-evoked population activity^1–14^, which manifest, among others, within the laminar microcircuits of primary visual cortex (V1). Spontaneous activity reflects the intrinsic dynamics of the brain at rest, in the absence of external stimulation or task demands. Under these conditions, pairwise functional connectivity, a measure of synchrony across pairs of neurons, is shaped directly or indirectly by underlying anatomical connectivity. Functional-connectivity analysis at resting state can therefore reveal the functional architecture of ensembles of neurons that share paths of anatomical connectivity with each other. It has been proposed that activity patterns expressed by such ensembles constitute a “vocabulary space,” capturing core elements of information processing in the cortex, which are thought to overlap with population activity patterns evoked by sensory stimulation^15–17^. Whether and how spontaneous ensemble functional architecture constrains stimulus-evoked population dynamics, however, remains an open question.

Here, we used two-photon calcium imaging to record activity from thousands of pyramidal neurons approximately simultaneously across layers 2–4 in mouse V1 and examined how neuronal ensembles emerge from pairwise functional connectivity (see methods) within and across cortical layers, revealing principles that govern interlaminar communication between granular and supragranular circuits. Our approach is “local” in nature and contrasts with prior studies that relied on regression or dimensionality-reduction methods to extract low-dimensional subspaces for modeling inter-areal communication, correlating neural activity with behavior, encoding stimulus dimensions^18^, capturing global trends, or characterizing response geometry^19–21^. While highly informative, these approaches typically depend on pre-specified task or behavioral variables and emphasize global response dynamics, potentially overlooking local computational structures that do not align with those dimensions.

Recent advances have mapped anatomical connectivity with unprecedented resolution^22^, providing a static description of cortical wiring. However, structural connectivity alone does not specify how neurons dynamically coordinate their activity, nor how functional ensembles interact across cortical layers. In contrast, our approach focuses on functional connectivity, capturing coordinated, time-varying interactions among specific neurons to reveal ensemble organization at single-cell resolution.

We leverage pairwise functional connectivity (see methods^23^) to define *first-order functionally-connected* (1FC) group of neurons that co-activate synchronously with a reference neuron more frequently than expected by chance and are separated by a single functional connection. We then examine the architecture and response properties of these ensembles across granular and supragranular layers of mouse V1 during both spontaneous activity and visual stimulation and assess how their organization is modulated across brain states, including quiet wakefulness, locomotion, and varying levels of alertness.

We argue that these functionally-defined modules reveal core systems-level principles by which V1 organizes activity, implementing algorithmic rules that govern signal propagation and ensemble-to-ensemble communication across cortical layers, thereby shaping information flow and supporting cortical computation. These principles generalize across spontaneous and stimulus-evoked activity, different brain states and level of alertness, pointing to a general computational motif of cortical organization. This work complements large-scale anatomical and dimensionality-based approaches, contributing a tractable, biologically grounded model of how cortical microcircuits process information.

Functional connectivity approaches based on aggregate signals—such as local field potentials or BOLD activity, which reflect the coordinated activity of large neuronal populations—have been instrumental in revealing principles of large-scale brain integration and its disruption in neurological and psychiatric disorders^24,25^. To our knowledge, this study presents the first meso-scale application of resting-state functional connectivity analysis at single-cell resolution to probe intrinsic network computations within the cortical column. This ensemble-based framework provides a principled abstraction for understanding cortical circuit function and dysfunction and is naturally suited for integration into deep learning and spiking neural network models, enabling systematic exploration of biologically grounded neuro-computational hypotheses.

## 2. Results

### Data Collection and Pre-Processing

Five postnatal 70–85 (10–12-week-old) F1 BL6-SLC17a7-Cre X Ai162 mice, expressing GCaMP6s in pyramidal neurons, underwent mesoscopic two-photon imaging while head-fixed on a treadmill during quiet wakefulness. Imaging covered dorsal primary visual cortex (V1) and adjacent extrastriate areas (Figs. 1B, 1C), spanning a ∼1.2 × 1.2 *mm*^2^ field of view (Fig. 1D). Data were acquired at ∼6Hz while simultaneously sampling four planes corresponding to V1 layer 2 (L2, 80-210*µm*), L3 (285-330*µm*), L4 (350-400*µm*), and L5 (500-600*µm*) (Figs. 1A-C). Imaging data were motion-corrected, automatically segmented, and deconvolved using the CNMF-CaImAn pipeline^26^. Deconvolved signals were thresholded (Methods) to generate binary calcium event trains (“eventograms”) used for subsequent analyses. Neurons with mean event rates below 0.01 Hz (0.27% of V1 neurons, on average) and neurons located within 15 µm of the field-of-view boundary (7.76% of V1 neurons) were excluded to minimize potential artifacts (see Suppl. Tables 1,2). Threshold values were selected to yield population event rates comparable *2 RESULTS* to those reported previously for mouse visual cortex^27^ (Fig. 1E). Importantly, sensitivity analysis revealed that functional connectivity measurements were robust to the choice of calcium deconvolution threshold (see below and Suppl. Fig. 1). We first explore functional connectivity properties in the absence of visual stimulation.

**Fig. 1.**
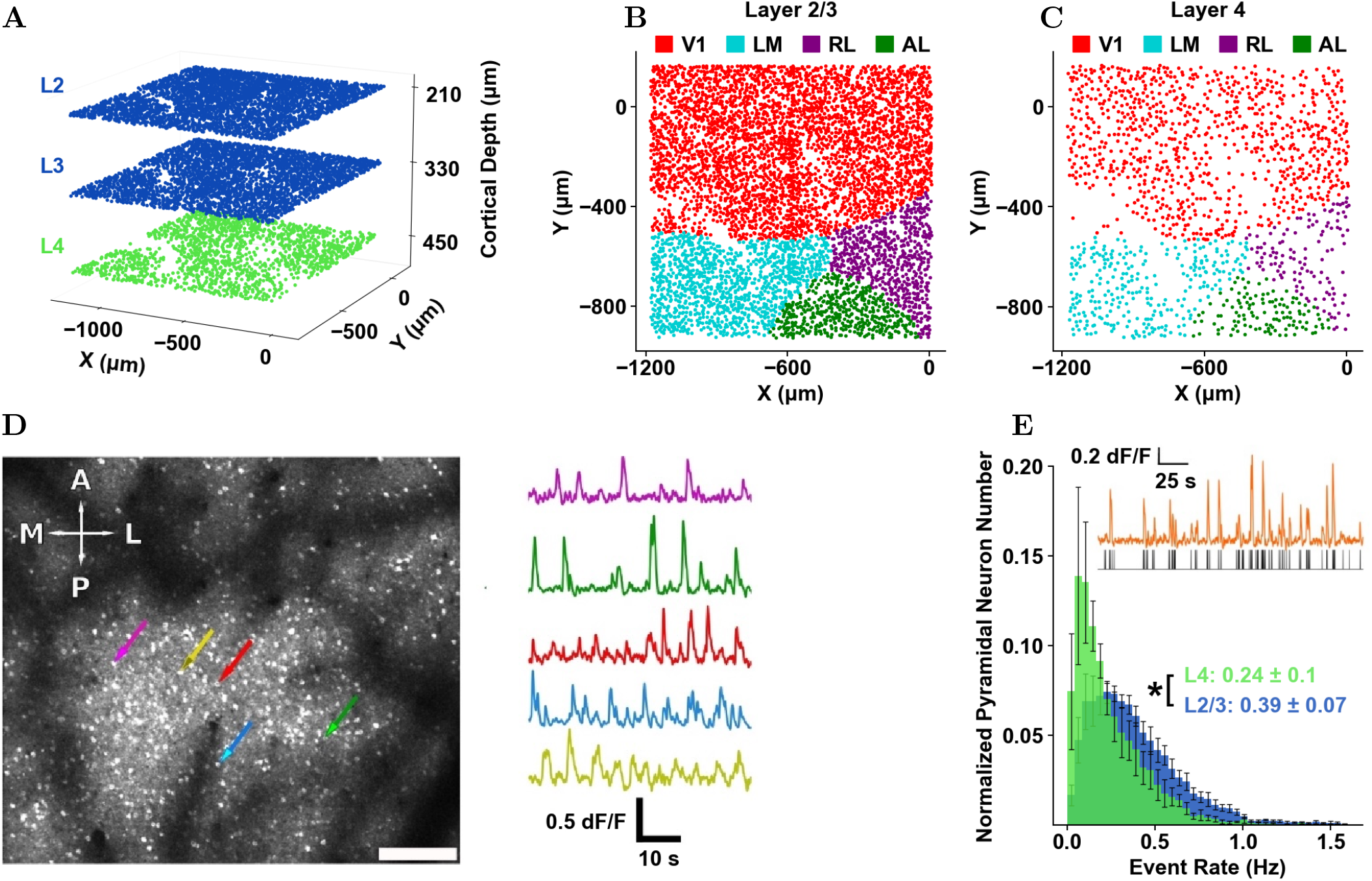
Imaging Paradigm. **A**) Illustration of L4, L2, L3 fields of view (FOVs) simultaneously acquired at ∼6Hz. L2/3: blue. L4: green. **B**) Retinotopic map of the FOV acquired in L2/3 for an example mouse. **C**) Retinotopic map of the FOV acquired in L4 for an example mouse. The boundary between the visual areas was determined on the basis of retinotopic mapping that was performed is described in Bae *et al*.^22^. **D**) Example FOV acquired in layer 2/3 at depth 210 µm. A: anterior, L: lateral, P: posterior, M: medial. Bar = 75 µm. Color arrows indicate 5 example cell bodies whose traces are shown in color on the right. **E**) Pyramidal neuron event rate histogram across animals. Error bars represent the standard error of the mean across mice (n=5). Inset: Mean event rate ± standard deviation of the sample means across mice (n=5), for each layer. P-value: “*”*<* 0.05. The highest p-value obtained from the permutation of means, the Welch’s t-test, and the ANOVA F-test is considered for the level-of-significance. Inset: deconvolution procedure. Orange: representative calcium responses. Gray: deconvolved firing probability traces obtained using the CaImAn algorithm^26^. dF/F: fractional fluorescence change. Gray traces in the bottom represent the thresholded, binarized, probability that specific imaging frames contain a calcium event (0: no event; 1: event).

### Functional connectivity analysis in the absence of stimulus

Pairwise functional connectivity was quantified using the spike time tiling coefficient (STTC)^23^, symmetrized and normalized to lie in [-1, 1]. STTC is defined relative to a synchrony window Δt; here we set Δt=0, restricting synchrony to events occurring within the same imaging frame (∼158.7 ms). Accordingly, our estimates are insensitive to sub-frame timing differences; nonetheless, we observed significant functional coupling and obtained robust results despite this relatively coarse temporal resolution. STTC was selected (Methods) because it is largely independent of firing rate and performs favorably relative to thirty-three other commonly used correlation measures^23^, rendering it well suited for assessing inter-neuronal correlations independent of neuronal activity level. Statistical significance was assessed by comparing observed STTC values to null distributions generated by circularly shifting calcium event time series 500 times, yielding z-scored functional connectivity measures. Observed pairwise STTC values extended well beyond their corresponding null distributions in both positive and negative directions, with a stronger positive tail (Figs. 2A, B: L4-L2/3 inter-layer pairs; Suppl. Figs. 5, 7: intra-layer pairs; Suppl. Fig. 9A: negative correlations). Accordingly, a substantial fraction of pyramidal neuron pairs exhibited significant positive correlations, consistent with prior work^28–31^. We focus on positively correlated pairs to characterize cooperative ensemble structure.

**Fig. 2.**
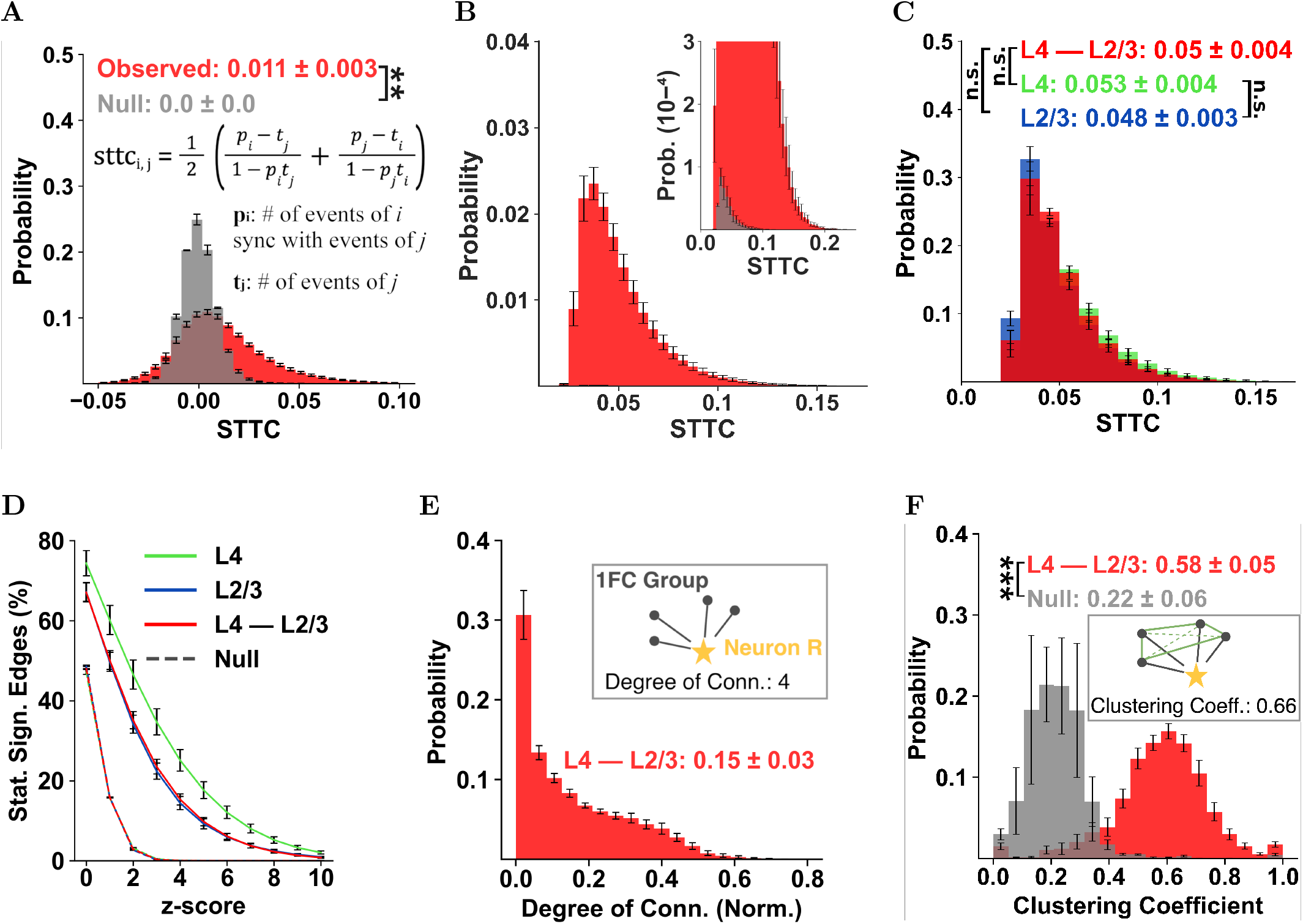
Inter-Layer First-Order Functionally-Connected (1FC) Groups from L4 to L2/3. **A**) The distribution of actual (red) and null (gray; circular shifted control) STTC weights from L4 to L2/3 (inset: the STTC computation formula; see methods for more details). **B**) Histogram of statistically significant (z-score *>*4) positive functional correlations. The magnified inset reveals the tail of the null distribution. **C**) Histogram of STTC weights of statistically significant (z-score *>*4) positive functional correlations L4 (green), L2/3 (blue) and L4−L2/3 (red). The distributions are remarkably similar both within and across layers. **D**) Percentage of statistically significant connections as a function of the significance threshold (z-score) for all layer cases. The latter two curves are nearly identical. **E**) The histogram of *normalized* degrees of connectivity from L4 →L2/3 (red) expressed as a fraction of the total number of neurons belonging to L4. Only edges with statistically significant (z-score *>* 4) positive STTC were included in the calculation. The degree of connectivity distributions (size of the groups) is asymmetric with a long tail to the right, with a mean of ≈ 138 neurons and a standard deviation of ≈ Given that cortical connections are weak, this is not an unreasonable number of neurons to be functionally connected with, in testament to the promiscuity of cortico-cortical connectivity we make in this work. The inset shows the first-order functional connectivity (1FC) group of an example neuron (star; R), i.e., the four direct neighbors of R (degree of connectivity of R: 4). **F**) Distribution of clustering coefficients of L2/3-neuron 1FC groups in L4, computed after excluding the frames containing calcium events of the L2/3 neurons themselves. The inset shows the clustering coefficient of an example (star) neuron, i.e., the extent of connectivity among its neighbors (here 4 of 6 possible connections in green). Figure insets represent the mean ± standard deviation of the sample means across mice (n=5); Error bars represent the standard error of the mean across mice (n=5). P-values: “**”: *<* 0.01; “***”: *<* 0.001, and “n.s.”: non statistically significant. The highest p-value obtained from the permutation of means, the Welch’s t-test, and the ANOVA F-test is considered for the level-of-significance.

At a conservative threshold of z-score*>*4, approximately 14% of neuronal pairs from L4 to L2/3 exhibit significant functional correlations (Fig. 2D) compared to ∼25% within L4 and ∼14% within L2/3 (Fig. 2D). A lower prevalence of positively correlated pairs within L2/3 (Fig. 2D; Suppl. Fig. 7E) is consistent with literature reports of more sparse firing in supragranular layers^32^. Unsurprisingly, STTC magnitude increased monotonically with significance (Suppl. Fig. 5). However, despite this, the large majority of functional connectivity weights remained small, consistent with Ecker *et al*.^32^: most STTC values were *<<* 0.1 (Figs. 2A, **B**; Suppl. Fig. 7), with mean STTC = 0.011 (interlayer L4-L2/3), 0.018 (within L4), and 0.010 (within L2/3). Notably, among significant pairs, STTC weight distributions were essentially indistinguishable within and across layers (Fig. 2C), consistent with a population-level homeostatic constraint that maintains the scale of pairwise functional coupling across the cortical microcircuit. Extensive sensitivity analysis showed that, although the fraction of significantly connected neuronal pairs naturally varied with the significance cutoff (Fig. 2D; Suppl. Fig. 7), our key findings remained unchanged across a range of significance thresholds and were insensitive to the choice of functional connectivity metric (STTC versus Pearson; see Suppl. Fig. 4).

### First-Order Functional-Connectivity (1FC) Groups

To extend the analysis beyond pairwise correlations, a natural graph to investigate consists of a pyramidal neuron with its significantly correlated (functionally-connected) pyramidal neighbors. This is the neuron’s *first-order functional-connectivity* (1FC) group. The 1FC group represents the simplest multi-neuronal ensemble beyond the single neuron and therefore provides a principled starting point for higher-order analysis. Its size—the degree of connectivity (DoC)—indexes how broadly the “index” neuron participates in functional interactions. DoC distributions within and across L4 and L2/3 are long-tailed (Fig. 2E; Suppl. Figs. 8A,C-D), consistent with hub-like organization and robust small-world network architecture^33,34^ (Suppl. Figs. 14,15). Moreover, 1FC groups are strongly *cooperative*: their clustering coefficients substantially exceed null expectations, even when computed using only frames in which the index neuron is silent (Fig. 2F; Suppl. Fig. 8B), indicating robust co-firing among group members that is not trivially driven by the index neuron itself. Figs. 2E–F summarizes the degree of connectivity and clustering coefficients of the L4-1FC groups associated with L2/3 neurons. These groups exhibited elevated within-group synchrony relative to null (Fig. 2F), consistent with coherent ensemble co-activity. Although our temporal resolution precludes assigning definite directionality or causality (but see Suppl. Fig. 16B), we asked whether L4-1FC co-firing nonetheless informs us about the responses of the corresponding index L2/3 neuron, motivating the analysis below.

### L2/3 Pyramidal Neuron Response as a Function of L4-1FC Group Cofiring

Fig. 3A illustrates a L2/3 neuron, whose probability of firing per imaging frame increases linearly with the number of cofiring events in its L4-1FC group after an initially flat region, approaching ∼1 at high cofiring values. In contrast, the null prediction remains essentially flat (gray line). The large majority of L2/3-neurons exhibit a monotonic increase in their probability of firing as a function of L4-1FC cofiring, well-fit by a ReLU function (Fig. 3A-B; Methods). We extrapolated the number of cofiring events that yield a response probability of 1 and histogram them in Fig. 3C. On average, this number is ∼70, commensurate to prior estimates made in a different context by Shadlen and Newsome^35,36^. Fig. 3D histograms the rising slope of ReLU fits with *R*^2^ ≥ 0.8, quantifying how much the response probability of L2/3-neurons rises per ten additional L4-1FC cofiring events.

**Fig. 3.**
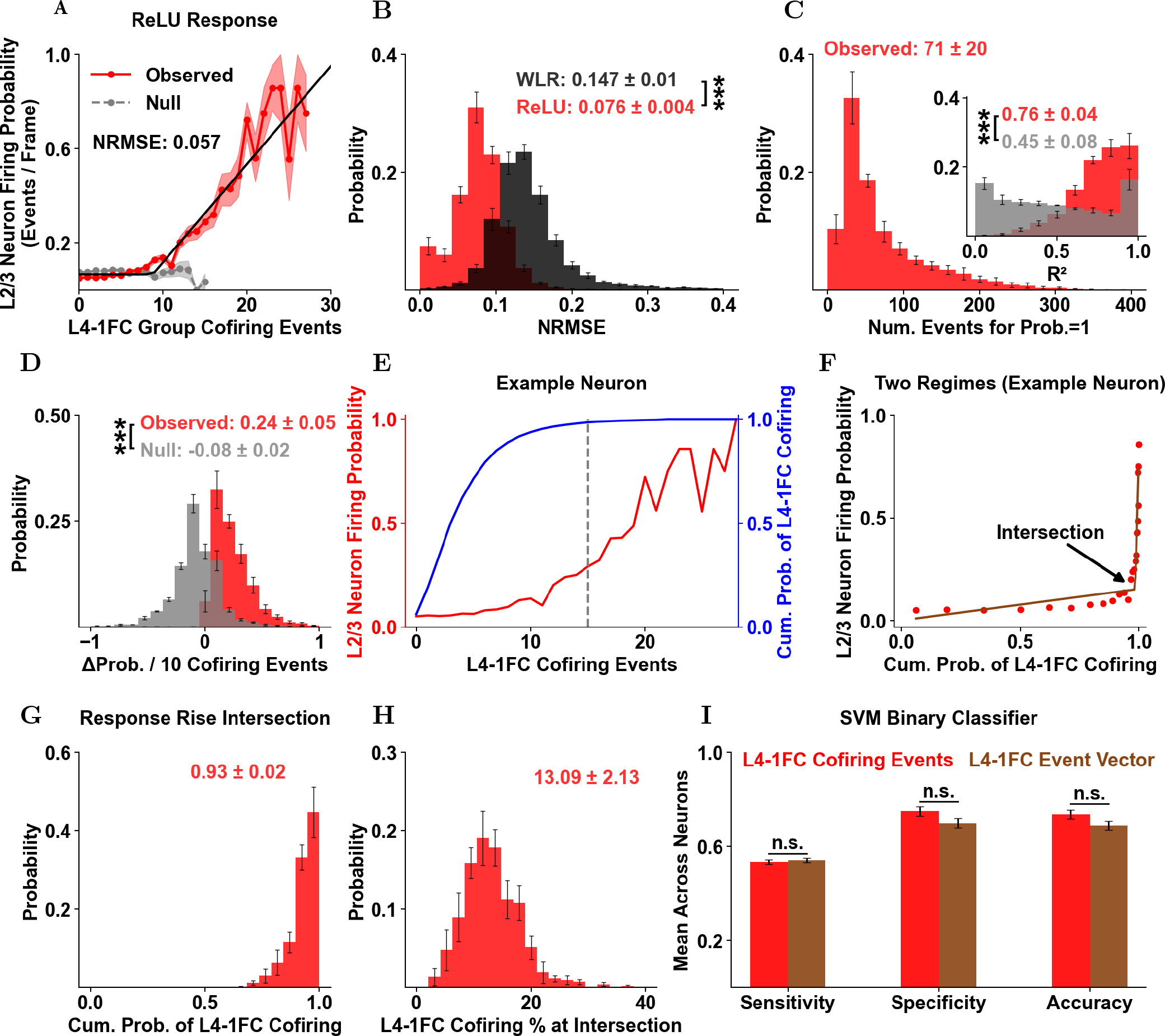
L2/3 Pyramidal Neuron Response as a Function of the Number of Cofiring of its L4-1FC Group. **A**) Calcium event firing probability of an example L2/3-pyramidal neuron as a function of the number of cofiring events in its L4-1FC (putative input) group (red line). Shaded region: SEM across cofiring events. Black Line: ReLU fit. Gray dashed line: Null (control; see Methods). **B**) Comparison of the normalized root mean square error (NRMSE) between the linear (black) and ReLU fits (red), indicating the superiority of the latter. **C**) Extrapolated number of cofiring events that yield probability of firing=1, derived from fits with *R*^2^≥0.8; Inset: Histogram of *R*^2^ values resulting from the ReLU fits, appropriately weighted by the standard error of each point (see methods). The majority of the fits are acceptable (*>*0.75) in contrast to null responses (gray), which exhibit significantly worse fits. **D**) Histogram of the slope of the ReLU fit, in units of the increase in probability of firing per 10 cofirings, derived from fits with *R*^2^≥0.8; **E**) The cofiring CDF of the L4-1FC group (blue line) plotted together with the response function (red line) for an example L2/3-pyramidal neuron. Dashed line: number of cofirings that correspond to the approximate intersection of the no-response and response regions. This L2/3-neuron fires for large, relatively rare, cofiring events. **F**) Firing probability of the neuron presented in **E** as a function of the cumulative probability of L4-1FC cofiring events. This neuron illustrates the typical response we observed across L2/3-cells, which exhibits a weak nearly flat response-region and an abrupt and steeply rising one, which emerges near the right tail of the cofiring event distribution, when cofiring events are large and have relatively low probability. The response can be fitted by two separate linear trends, shown here. The first line has a slope close to 0 and represents the *weak-response* regime, while the second demonstrates the *sharply rising* neuronal-response. For this neuron, the intersection of these regions occurs at ∼CDF=0.97. That is, only 3% of frames (rare events) have a sufficiently large number of L4-1FC cofirings to make the corresponding L2/3-neuron fire. **G**) Histogram of the cumulative probability of L4-1FC cofiring events corresponding to the start of the sharply rising response’s intersection of the two lines that fit the response. **H**) Histogram of L4-1FC group cofiring percentages that correspond to the response intersection. On average, the switch to the fast-rising response regime for L2/3-neurons occurs at CDF=0.93, which corresponds to ∼13% of its putative input group firing together. Only L2/3-neurons that reach firing probability ≥0.6, have a ReLU fitting *R*^2^≥0.8 (as in A), and a response-region (F) with a slope≥1 are included in E-G. The slope in F is estimated as the difference of the likelihood of response at two numbers of cofiring events over the difference of their corresponding cumulative probabilities. **I**) The knowledge of the identity of the L4-1FC neighbor neurons that fire, in addition to the L4-1FC aggregate cofiring, does not enhance the classification accuracy in determining the L2/3 neuronal firing. Specifically, here, we employ a Linear Support Vector Binary Classifier to predict L2/3-neuron firing during the spontaneous period using as input: the number of cofiring of its L4-1FC neighbors (red), and a binary vector of N elements, where N is the size of its L4-1FC group and each of its elements corresponds to a specific L4-neuron of said group, with a value that indicates whether or not that neuron fires at a particular frame (brown). Note that the input of the second case (brown) includes the identity of the neurons that fire. We compared their performance, in terms of sensitivity, specificity, and accuracy (see Methods). Histogram insets: mean ± standard deviation of the sample means across mice (n=5). Error bars: SEM across mice (n=5). Histogram insets represent the mean ± standard deviation of the sample means across mice (n=5). Error bars: SEM across mice (n=5). P-values: “***” *<*0.001. The highest p-value obtained from the permutation of means, the Welch’s t-test, and the ANOVA F-test is considered for the level-of-significance.

Consistent with their ReLU functional form, L2/3-neurons exhibit two response regimes: a flat region of minimal slope followed by a sharp transition to a region of high slope. This becomes especially evident when L2/3 neuron responses are plotted against the probability of occurrence of the cofiring events that elicit them. Fig. 3E illustrates a L2/3 neuron, whose response (red) is plotted together with the cumulative probability distribution (blue) of the cofiring events in its L4-1FC group. Evidently, L2/3 neuron firing probability starts to rise for events lying relatively far in the tail of the cofiring event size distribution (Figs. 3E-F). On average, L2/3-neurons tend to remain silent for the majority (∼93%) of cofiring events occurring within their L4-1FC groups, their probability of firing beginning to rise sharply after cofiring event number crosses into the ∼7% of largest and rarest L4-1FC group cofiring events (Figs. 3F-G). This corresponds to ≥13% of L4-1FC neurons firing in synchrony (Fig. 3H). In sum, L2/3-neurons respond sparsely, their probability of firing rising sharply only relatively rarely, when a high number of cofiring events occurs in their input group. In order to further understand the relationship between L4-1FC cofiring events and index L2/3 neuron firing, we employed a series of classifiers to predict frame by frame whether L2/3 neurons fire from the firing events recorded in their L4-1FC groups. Consistent results were obtained using SVM classifiers with linear kernel (Fig. 3I) as well as Random Forest, Naive Bayesian and Logistic regression (Suppl. Fig. 19). Remarkably, the accuracy of predicting L2/3 responses did not improve when, in addition to the aggregate number of cofiring events, we included information about the identity of L4-1FC group neurons that fire (Fig. 3I). Instead, we found that L2/3-neuron responses scale with the number of cofiring events within their L4-1FC groups and are comparatively insensitive to the specific identities of the participating neurons. We emphasize, as mentioned earlier, that our analysis does not establish causal directionality. Nonetheless, the structure of the observed statistical dependence is informative and suggests that L4-1FC group co-activity captures a relevant signal that could reflect, at least in part, convergent drive onto the corresponding L2/3 neuron. Building on these ensemble-level signatures we introduced above, we next asked whether the extent of an L2/3 neuron’s functional coupling to L4—as quantified by its L4-1FC degree of connectivity—systematically relates to its properties.

#### L2/3 neuron properties depend on L4-1FC group size

The slope of L2/3 neuron response functions depends on L4-1FC group size. To relate L2/3 spiking to L4 ensemble activity, we computed the mean firing probability of each L2/3 neuron as a function of the number of co-firing events within its L4-1FC group and stratified neurons by L4-1FC size (DoC) quartiles (small: 0–25%, middle: 37.5–62.5%, large: 75–100%). L2/3 neurons with smaller L4-1FC groups exhibited systematically steeper input–output functions than those with larger groups (Fig. 4A). This effect is unlikely to reflect incomplete sampling of L4 neurons (e.g., out-of-plane neurons that cannot be imaged): when we restricted large L4-1FC groups to subsets matched in size to “small” groups, the resulting control curves did not recapitulate the steep slopes observed for neurons with genuinely small L4-1FC groups, particularly during epochs of low population activity (Suppl. Fig. 23A). Therefore, the observed difference in slopes does not reflect imaging bias. We next tested whether the slope differences reflect a simple scaling with the size (*N*) of the associated L4-1FC. If L2/3 neurons operate over comparable output dynamic ranges while sampling different numbers of effective inputs, independent-input arguments predict that adaptation will force response slopes to come together when cofiring is measured in units Of 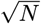, i.e., after normalizing by *N* ^0.5^ (Suppl. Section 5). Empirically, the input–output curves became identical when cofiring was rescaled by *N* ^*α*^, with *α* ≈ 0.6–0.7 across all five mice (Fig. 4B). The excess over 0.5 is consistent with correlated activity and elevated clustering within the L4-1FC groups. Under this normalization, L2/3 neurons exhibited an approximately common, near-linear response function largely independent of *N* (Fig. 4B), and the stability of *α* across animals and different L4-1FC group sizes suggests a conserved scaling property of the circuit.

**Fig. 4.**
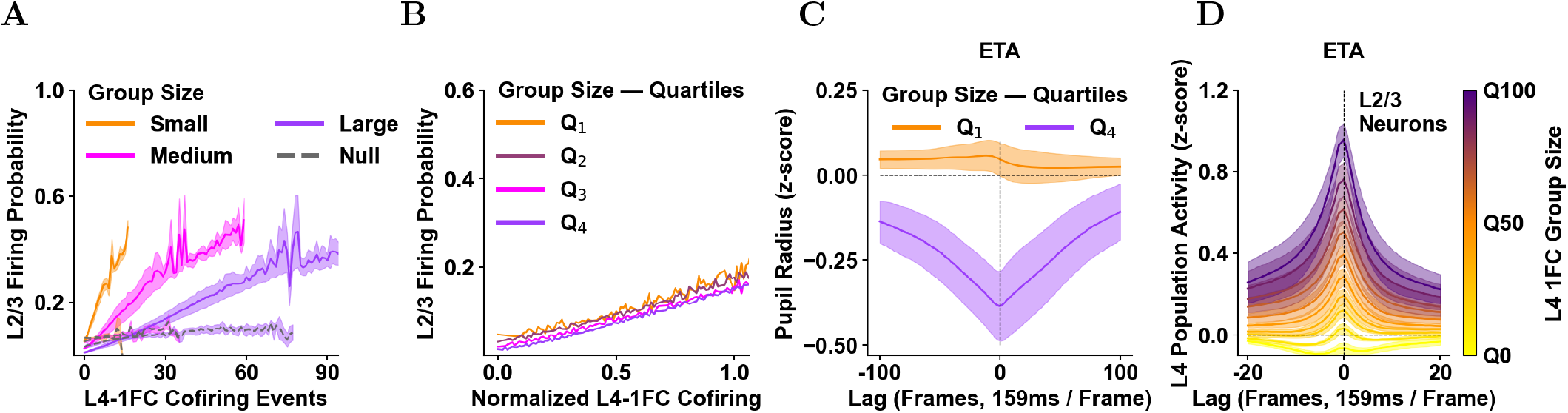
Likelihood of L2/3 Neuron Firing as a Function of L4-1FC Group Cofiring for Different L4-1FC Group Sizes. **A**) Average probability of firing of L2/3 neurons as a function of the number of co-firing events in their L4-1FC groups. Orange: L2/3 neurons with L4-1FC *group sizes* belonging to the smallest quartile (Q0-Q25). Purple: L4-1FC group sizes belonging to the largest quartile (Q75-Q100). Pink: L4-1FC group sizes belonging to the middle quartile (Q37.5-Q62.5). Note that smaller cofiring events translate into a larger probability of firing for small groups (orange) compared to medium or large ones. **B**) Interestingly, there is a power of N (size of the L4-1FC group), normalizing by which results in identical slope for all groups of different sizes (quartiles Q1 to Q4). The normalization is done by dividing by the L4-1FC group size raised to the power of a constant. Here the plot shows one example mouse with a normalization constant of ∼0.7. Across mice the constant varied from ∼0.6-0.7 (see Methods for a detailed discussion). Groups with sizes less than 5 have been discarded from Q1 to minimize low number bias. Shaded regions: SEM across mice (n=5). **C)** Event Triggered Average (ETA) on pupil radius of L2/3 recipient neurons that have small (quartile 1; yellow line) vs. large (quartile 4; purple) L4-1FC groups, respectively. Note that L2/3 neurons with small L4-1FC groups tend to fire on average with a weak, if any, dependence on pupil size, while L2/3 neurons with large L4-1FC groups tend to fire when pupil size is low. **D)** ETAs on the aggregate L4 neuronal activity for L2/3 neurons with different L4-1FC sizes. This confirms that the coupling to aggregate L4 population activity is weaker for neurons with small L4-1FC sizes. The shape of the ETA is also different. For L2/3 neurons with groups of large sizes, L4 population activity increases prior to the firing event, which tends to occur at the peak of the population wave of activity. For L2/3 neurons with small L4 groups, the modulation is much weaker and there appears to be an epoch of aggregate activity suppression before and after the firing event. Overall neuronal behavior exhibits a continuum rather than splitting dichotomously into two discrete categories. In all figures, time is measured in imaging frames (∼159 ms) from the time of a calcium event fired by the L2/3 neuron (time zero).

L4 functional connectivity coupling stratified how the activity of L2/3 neurons get modulated by internal-state variables under spontaneous conditions. L2/3 neurons with small L4-1FC groups showed, at best, weak modulation by pupil diameter (Fig. 4C) and by global population activity (Fig. 4D), whereas neurons with large L4-1FC groups were strongly modulated by these parameters (Figs. 4C-D, Suppl. Figs. 21 B,D, 25-26). This pattern parallels the “soloist–chorister” framework described previously^37^; importantly, the transition is graded, with modulation strength varying smoothly across the continuum of L4-1FC group sizes rather than defining discrete soloist versus chorister classes. Consistent with this stratification, L2/3 pyramidal neurons with large L4-1FC groups exhibited substantially coordinated firing, whereas L2/3 neurons with small L4-1FC groups fired nearly independently from each other (Suppl. Figs. 17A). Specifically, for 95% of L2/3-neuronal pairs drawn from the small L4-1FC pool, the observed cofiring probability did not differ significantly from the product of the marginal firing probabilities (Suppl. Fig. 17A), as expected under independence. Together, these results indicate that, in the absence of stimulus, L2/3-neurons with small L4-1FC groups constitute a comparatively decorrelated channel for processing circuit activity, whereas L2/3 neurons with large L4-1FC groups participate in a more coordinated, state-coupled subnetwork. This trend is still present, though much weaker, under sensory stimulation, as both large and small L4-1FC groups now reflect in large part stimulus correlations (Suppl. Fig. 44 A-B). As they are only weakly modulated by internal state, L2/3 neurons with small L4-1FC groups exhibit similar firing rates across different internal-state conditions (e.g., during high versus low population firing), whereas L2/3 neurons with large L4-1FC groups are more strongly modulated by internal state, showing state-dependent firing (Suppl. Figs. 21).

Having established in prior sections the relationship (ReLU) between the probability of firing of a L2/3 neuron and the cofiring of its associated L4-1FC group, we next asked how this might generalize to ensemble-to-ensemble interactions between L4 and L2/3 populations. This is motivated by prior work indicating that single neuron–to–neuron influences in cortex are typically weak^38^, implying that reliable activity propagation is more plausibly mediated by coordinated recruitment of neuronal ensembles (ensemble-to-ensemble communication)^39^. In what follows, we therefore aim to identify cross-laminar ensembles whose co-activation may constitute a putative channel of communication between L4 and L2/3.

#### Ensemble to Ensemble Functional Coupling between L4 and L2/3

Functionally connected L2/3 neuron pairs shared substantially more L4 partners than nonconnected pairs, as quantified by the overlap of their respective L4-1FC groups (Fig. 5B; Suppl. Fig. 30). This enrichment is consistent with the idea that L2/3 pairwise functional correlations reflect, in part, shared coupling to L4. Furthermore, L4-1FC overlap co-varied strongly with the overlap of the corresponding L2/3-1FC groups (Suppl. Fig. 30C). Overlap was dependent on the pair’s functional correlation strength/significance but was otherwise only weakly, if at all, dependent on inter-somatic distance or orientation-preference similarity (Suppl. Figs. 30D-E). This suggests that the cross-laminar network embedding of pairs of L2/3 neurons does not depend strongly on local proximity or even on tuning similarity.

**Fig. 5.**
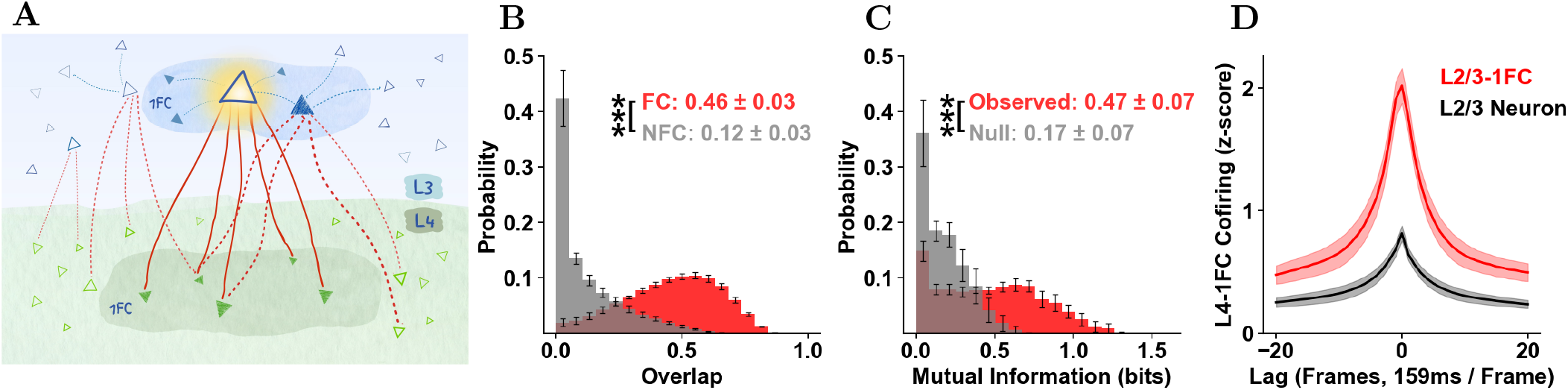
Pathways: Functional Coupling between L4 and L2/3-1FC Groups of an L2/3 neuron. **A**) Conceptual illustration of a pathway of an L3 neuron (highlighted target neuron), which includes the pyramidal neurons (filled triangles) of its L4-1FC and L2/3-1FC groups, encircled in green and blue shaded regions, respectively. Red arrows indicate the pairwise statistically significant positive functional connections between an L4-neuron and their putative recipient L3-neuron (solid: L4-1FC group of R; dashed: L4-1FC functional connections of *other* L3-neurons). Filled triangles indicate 1FC neighbors of the target neuron, whereas empty triangles denote neurons lacking functional connectivity to it (L4 neurons in green; L3 neurons in blue). L3-1FC neighbors of the target neuron share L4-1FC neighbors. L2 is omitted from the illustration for simplicity. Relationships illustrated for L3 can be directly extended to L2. **B)** Histogram of pairwise overlaps of L4-1FC groups of L2/3 neurons belonging to the same L2/3-1FC group (red) versus those in different L2/3-1FC groups (gray). The overlap of two groups is estimated as the size of the intersection of the two groups over the square root of the product of the sizes of the individual groups. The L4-1FC groups of two functionally-connected L2/3 neurons have larger overlap than pairs of L4-1FC groups of which their index L2/3 neurons are not functionally connected. **C**) Histogram of the mutual information between the L4-1FC group cofiring and the L2/3-1FC group cofiring of L2/3 neurons. The null is computed on a control L2/3-1FC group for each neuron that has the same size as the observed and is formed by random selection of L2/3 neurons that do *not* belong to the observed one. **D**) Event Triggered Average during the resting state of the group cofiring z-score of the L4-1FC groups of all L2/3 neurons, where lag 0 indicates frames where the L2/3 neuron exhibits a firing event (black) and frames where the L2/3-1FC group is active (red). The active frames of the groups are defined as the N frames with the highest L2/3 group cofiring, where N is the total number of events of the recipient L2/3 neuron during the entire resting-state period. The lines and shaded regions indicate the mean and SEM across mice, respectively. L2/3-neurons without L4-1FC groups have not been included. Inset values indicate the mean ± standard deviation of the sample means across mice (n=5). Error bars: SEM across mice (n=5). P-values: “**” *<* 0.01; “***” *<* 0.001. The highest p-value obtained from the permutation of means, the Welch’s t-test, and the ANOVA F-test is considered for the level-of-significance.

To test directly for cross-laminar ensemble coupling, it is natural to compare the cofiring in each L2/3 neuron’s L4-1FC group to the co-firing in the corresponding L2/3-1FC group. We found that L4-1FC cofiring carried significant mutual information about L2/3-1FC cofiring (Fig. 5C), supporting coordinated interactions at the ensemble-to-ensemble level.

We next asked whether ensemble-to-ensemble coupling is more informative than ensemble-to-single-neuron coupling. To enable a matched comparison, we discretized L2/3-1FC activity by thresholding its co-firing (z-scored) time series, selecting the threshold so that the number of suprathreshold L2/3-1FC events equaled the number of firing events of the corresponding index L2/3 neuron (Methods; Fig. 5D). Under this rate-matched framework, L4-1FC cofiring was more tightly coupled to the L2/3-1FC ensemble state than to the firing of the index L2/3 neuron (Fig. 5D), a result corroborated by decoding metrics (sensitivity, specificity, accuracy; Suppl. Fig. 31). Collectively, these analyses identify L4-1FC to L2/3-1FC group-to-group coupling as a robust candidate pathway for cross-laminar communication, exceeding the reliability of the corresponding group-to-single-unit relationship.

The observations described above were robust across different brain states. Specifically, the reported principles were preserved across levels of alertness using as proxy the pupil diameter (Suppl. Fig. 29) and held when locomotion epochs were analyzed separately from quiet wakefulness (Suppl. Figs. 34,37,38). The analysis presented thus far focused on resting-state activity, in the absence of visual stimulus. Because sensory stimulation introduces signal correlations that reshape functional connectivity, the next section asks whether the core phenomena identified in the spontaneous regime persist in the presence of stimulus-driven correlations. Below we show that the observations we made above generalized to periods of visual stimulation, even though stimulus-epoch functional connectivity ensembles (1FC-groups) differ substantially from 1FC-groups obtained at resting state, in the absence of stimulus.

#### Functional-Connectivity under Stimulus Presentation

We performed functional-connectivity analysis of data obtained with the animal resting under visual stimulation, consisting of a movie with pseudo-randomly interleaved segments containing motion direction and orientation information (“Monet”; see Materials and Methods). The functional connectivity was computed using the same pipeline as in the resting-state condition. Functional-connectivity architecture under stimulus presentation differs significantly from resting-state connectivity, exhibiting on average smaller STTC weights, smaller degrees of connectivity (DoC), smaller clustering coefficients (CC), and on average fewer statistically significant functional connections (Suppl. Figs. 41-42), that appear to be more concentrated near the soma (Suppl. Fig. 41J-K). As a result, resting-state functional-connectivity architecture is a weak predictor of signal correlation architecture, at least under the “Monet” visual orientation/direction stimulus conditions. Only ∼27% to 35% of statistically significant functional connections identified in resting-state remain significant under stimulus conditions (Suppl. Fig. 51A), while L4-1FC groups of L2/3 neurons display on average only ∼27% overlap under the two conditions (Suppl. Fig. 43A). This is consistent with the weak relationship seen between orientation/direction tuning similarity and pairwise functional connectivity strength at resting state, in the absence of stimulus (Suppl. Figs. 11B vs. 48), and is further corroborated by the observation that cofiring in L4-1FC groups defined at *resting-state* (i.e., **r**L4-1FC) is a poor predictor of L2/3-neuron activity under stimulus conditions (Fig. 6E).

**Fig. 6.**
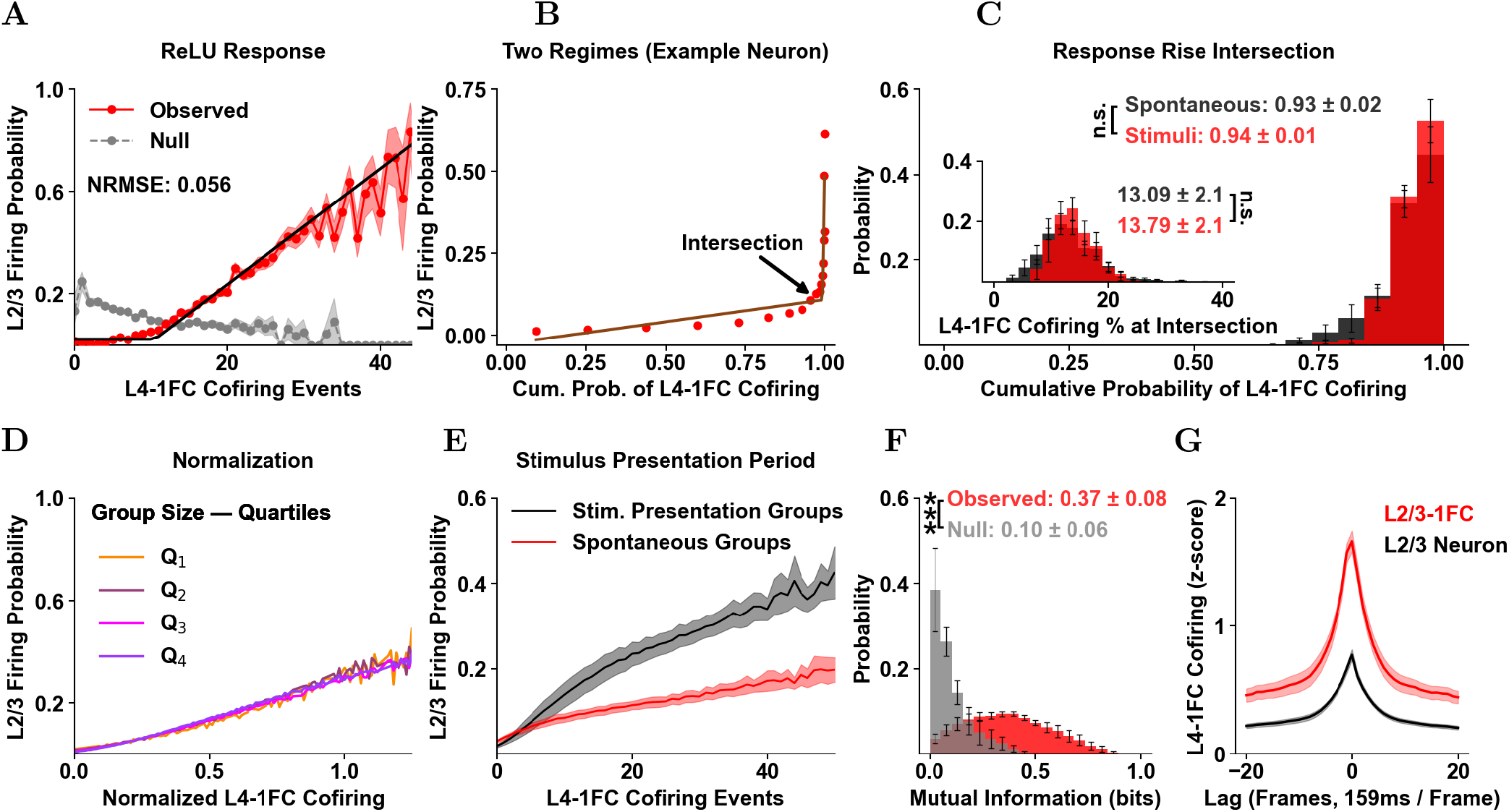
The key organization properties of the L2/3 pyramidal neuron response as a function of the number of cofiring of its L4-1FC group identified during resting state persist also under visual stimulation. **A)** Calcium-event firing probability of an example L2/3 neuron as a function of the number of cofirings in its L4-1FC group (red), computed during stimulus presentation. Shaded region, SEM across cofirings; black, ReLU fit; gray dashed, null control (see Methods). Unlike the observed response, the null remains approximately flat and shows fewer higher-order cofirings. **B**) Firing probability of the same neuron as a function of the cumulative probability of cofirings, as in 6A. This L2/3 neuron fires for *large, relatively rare* cofirings. **C**) Histogram of the cumulative probability value (CDF) of cofirings at the onset of the sharply rising response, defined by the intersection of the two fitted lines. Inset, histogram of the cumulative probability of L4-1FC cofiring percentages at the response intersection. On average, the transition to the fast-rising regime occurs at CDF = 0.94 for L2/3 neurons, corresponding to ∼14% of the putative input group firing together. Included were only L2/3 neurons with firing probability ≥ 0.6 and ReLU fit *R*^2^ ≥ 0.8; neurons with response-region slope *<* 1 were excluded (∼2% on average across mice). Slope was estimated as the change in response likelihood between two cofiring counts divided by the change in their corresponding ECDFs. **D)** Firing probability of L2/3 neurons as a function of normalized L4-1FC-group cofiring, obtained by dividing by group size raised to a constant power. Shown is one example mouse with normalization constant ∼0.7, as used for the resting-state period in Fig.4B. **E)** Response functions of L2/3 neurons during stimulus presentation for cofiring in L4-1FC groups defined during spontaneous conditions (Spont., red solid) or stimulus presentation (Stim., black solid), and for control groups of matched size containing randomly selected neurons outside the original group (dashed). **F**) Histogram of the mutual information between L4-1FC cofiring and L2/3-1FC cofiring in L2/3 neurons. The null was computed using a control L2/3-1FC of matched size, formed by randomly selected L2/3 neurons outside the observed group. **G**) Event-triggered average during stimulus presentation of the L4-1FC cofiring z-score for all L2/3 neurons: lag 0 marks frames in which the L2/3 neuron fires (black) and the L2/3-1FC is active (red). Active group frames were defined as the N frames with highest L2/3-group cofiring, where N is the total number of events of the recipient L2/3 neuron during the full stimulus-presentation period. The lines and shaded regions indicate the mean and SEM across mice, respectively. Histogram insets report the mean ± standard deviation of the sample means across mice (n=5). Error bars: SEM across mice (n=5). P-values: “n.s.”: non statistically significant. The highest p-value obtained from the permutation of means, the Welch’s t-test, and the ANOVA F-test is considered for the level-of-significance. Materials and Methods describes the experiment involving directional visual stimuli. Functional connectivity was computed using the same pipeline as in the resting-state condition, applied to the data from these recordings.

A substantial body of work has shown that spontaneous cortical activity can recapitulate, or at least anticipate, salient features of stimulus-evoked population patterns—supporting the idea that ongoing dynamics reflect an internalized model of sensory statistics^16,40–42^. The results above refine this perspective by demonstrating that fine-grained, functional-connectivity architecture is strongly context dependent: for example, the naturalistic “Monet” stimulus (see methods) changes network structure by reducing overall coupling, degree of connectivity and clustering coefficients, while increasing spatial localization of significant interactions (Suppl. Figs. 41–42).

Consistent with this reconfiguration, only a minority of resting-state significant edges and L4-1FC membership relationships are preserved under stimulation (Suppl. Figs. 51,43A), while co-activity in resting-state defined rL4-1FC groups carries limited information about L2/3 firing during the stimulus epoch (Fig. 6E). Thus, although resting-state activity captures important aspects of population-level activity structure, it provides a weak predictor of stimulus-epoch functional coupling rather than constituting a fixed graph invariant across different stimulation conditions.

However, despite these differences, the key observations we made regarding the relation of ensemble patterns of activity between layers L4 and L2/3 remain largely consistent under stimulus conditions: As in resting-state, L2/3-neuron response profiles conform to ReLU functions of L4-1FC group cofiring, provided L4-1FC groups are defined under the *stimulus* condition (i.e., **s**L4-1FC). Furthermore, L2/3 response functions remain largely consistent under stimulus conditions: comparable maximum firing probabilities (Suppl. Fig. 53I), closely matched extrapolated sL4-1FC cofiring event numbers required to reach a firing probability of 1 (71 vs 79; Fig. 3C, Suppl. Fig. 53H), analogous power-scaling dependence on sL4-1FC group size (Fig. 4, Suppl. Fig. 53M-N), similarly selective firing responses to relatively rare sL4-1FC cofiring events (Suppl. Fig. 53K-L), as well as improved reliability of sL4-1FC to sL2/3-1FC, group-to-group, activity transmission (Suppl. Fig. 55). The persistence of L4 to L2/3 ensemble-to-ensemble functional communication signatures across both spontaneous and stimulus-evoked conditions—despite substantial reconfiguration of pairwise connectivity—suggests they likely reflect a conserved organizational principle of cortical network dynamics.

To determine whether these functional connectivity observations generalize across neuronal subpopulations with different visual properties, we next compared orientation-tuned (“OT”) and non-orientation-tuned (“nOT”) V1 neuronal populations, the majority of the latter (67%-75%, depending on the layer) being visually unresponsive. Approximately 47% of V1 units were orientation tuned across mice. At restingstate, OT versus nOT units exhibited similar firing event rates, as well as similar normalized degree of connectivity (DoC) and clustering coefficient (CC) distributions within and across layers (Suppl. Fig. 40D-E, 49A-F). Interestingly, resting-state rL4-1FC groups of orientation-tuned L2/3-neurons contain ∼56% non-orientation selective units irrespective of group size, suggesting that functional-connectivity is promiscuous with respect to tuning properties under these conditions (Suppl. Fig. 52E). This conclusion is further supported by the weak relationship observed in the absence of stimulus between the strength of functional connectivity and orientation-tuning similarity (Suppl. Fig. 11). These observations change under visual stimulation (“Monet” stimulus) conditions, in that OT neurons exhibit on average higher firing event rates (Suppl. Fig. 49D) and L2/3 neurons with larger sL4-1FC groups are now more likely to be orientation tuned (Suppl. Fig. 49G), as expected, since functional connectivity is now driven in part by stimulus correlations. Consistent with this, the relationship between strength of functional connectivity and orientation tuning similarity is much stronger under stimulation conditions (Suppl. Figs. 48 vs 11). Despite this difference, OT versus nOT units exhibit again similar normalized DoC and CC distributions within and across layers (Suppl. Fig. 49E-F). Notably, the response functions of orientation-tuned L2/3-neurons as a function of their L4-1FC group cofiring events do not differ significantly from those of untuned L2/3-neurons, whether they are computed under resting or visual stimulation conditions (Suppl. Fig. 54).

It is natural to assume that the cofiring of sL4-1FC groups corresponding to orientation-tuned (OT) L2/3 neurons should convey information about the orientation of the Monet stimulus. To evaluate this hypothesis, we computed the probability of how much the orientation of the Monet stimulus differs from the orientation preference of the “index” OT L2/3-neuron as a function of the number of sL4-1FC cofiring events. The top row of Fig. 7A demonstrates that as sL4-1FC group cofiring increases, Monet stimulus orientation matches the orientation preference of the corresponding L2/3 “index” neuron more closely and can be predicted more accurately. A slightly weaker trend is observed for the sL2/3-1FC groups of orientation-tuned L2/3-neurons (Fig. 7A, second row). Predictably, functional-connectivity groups associated with non-orientation-tuned neurons convey little, if any, information about stimulus orientation (Fig. 7A, rows 3 and 4) even though they still convey information about the probability of firing of their “index” L2/3-neuron (Suppl. Figs. 54C-D). Functional-connectivity groups defined under resting-state conditions (rL4-1FC and rL2/3-1FC) also convey relatively low information about stimulus orientation (Suppl. Fig. 57B), confirming that resting-state functional-connectivity architecture fails to capture the bulk of the information encoding signal correlation structure observed under stimulation conditions.

**Fig. 7.**
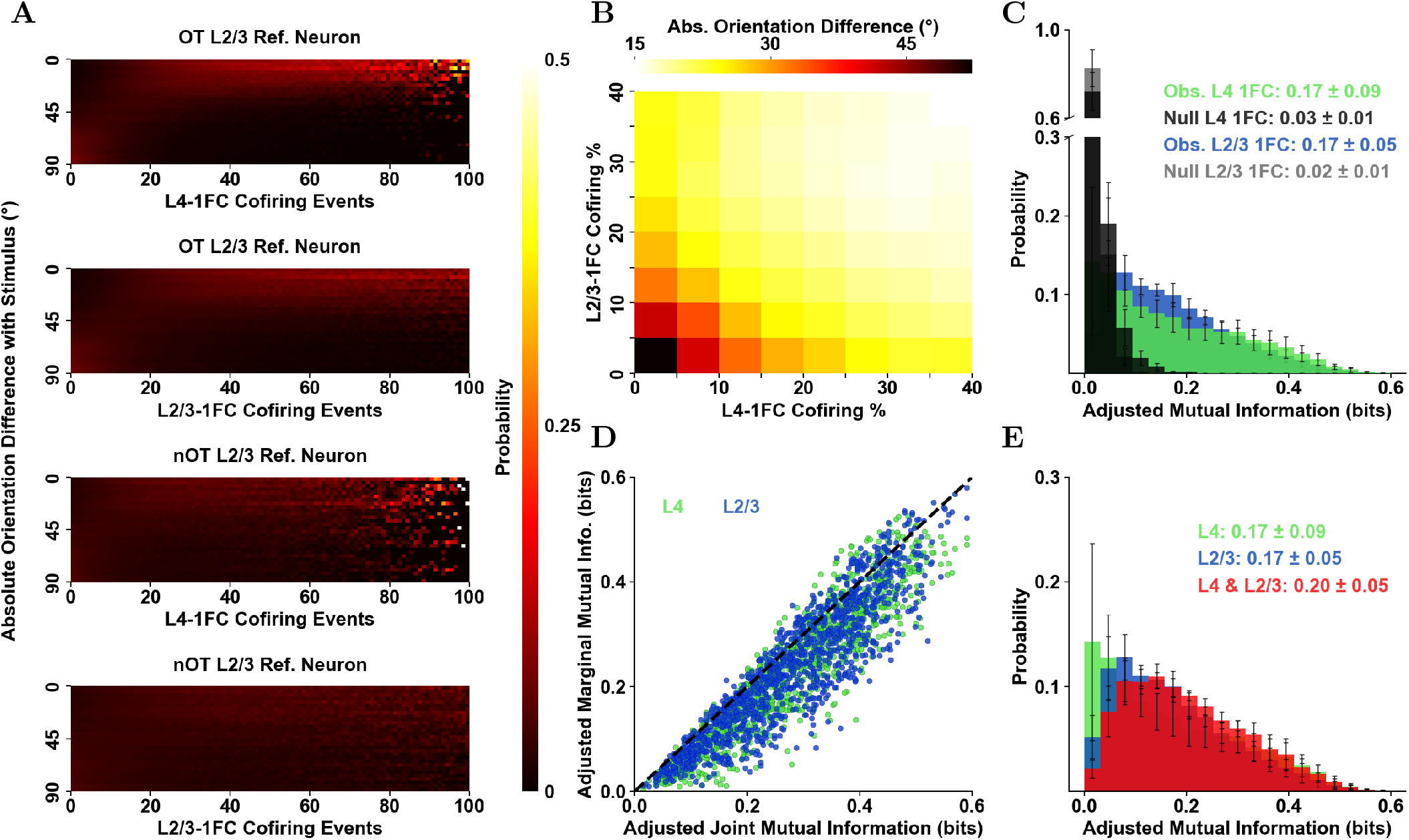
Prediction of the Angle of the Stimulus based on the Orientation Preference of the L2/3 Neuron and its 1FC Groups Cofiring. **A)** Prediction of the stimulus orientation as a function of the number of cofiring events of the L4-1FC group (first and third row) and the L2/3-1FC group (second and fourth row) for tuned and untuned L2/3-neurons. The y-axis histograms the absolute orientation difference between the angle of the stimulus and the direction preference of the corresponding L2/3-index neuron. The index neurons are orientation-tuned in the first and second row, and untuned in the third and fourth row. The color indicates the probability of each bin: the darker the color, the lower the likelihood of having this absolute orientation difference. **B)** For each L2/3-OT neuron, we calculate the median absolute orientation difference between its preferred angle and the orientation of the stimulus presented at frames where the co-firing percentage of its L4- and L2/3-1FC groups falls within the ranges specified on the x- and y-axes, respectively. Each square in the heatmap represents the mean across mice of the average of these medians across all L2/3 OT neurons. **C)** Mutual information of the L4-1FC group cofiring (green) and the L2/3-1FC group cofiring (blue) of L2/3 orientation-tuned neurons with the direction of the stimulus being presented (computed per frame). To account for potential estimation bias, we adjusted the values by randomly shuffling the stimulus angles 30 times, computing the mutual information for each shuffle, and subtracting the mean from the observed mutual information. The null distributions (dark grey: L4, light grey: L2/3) are computed by selecting control L4 and L2/3-1FC groups of identical size to the corresponding observed groups, ensuring no overlap with them (see Methods). **D)** Mutual information of the joint distribution of the L4- and L2/3-1FC group cofiring (x-axis), the L4-1FC group cofiring (y-axis, green) and the L2/3-1FC group cofiring (y-axis, blue) of L2/3 orientation-tuned neurons with the direction of the stimulus being presented for an example mouse (computed per frame). **E)** Histogram of the *joint mutual information* of each L2/3-OT neuron’s L4-1FC and L2/3-1FC group co-firing per frame with the stimulus (red). The information that the L4-1FC and L2/3-1FC *jointly* provide about the stimulus is greater than the information provided by randomly selected groups. Histogram insets: mean ± standard deviation of the sample means across mice (n=5). Error bars: SEM across mice (n=5). P-values: “***” *<*0.001, and “n.s.”: non statistically significant. The highest p-value obtained from the permutation of means, the Welch’s t-test, and the ANOVA F-test is considered for the level-of-significance.

Fig. 7B confirms that epochs of high activity in both sL4-1FC and sL2/3-1FC groups correspond to visual stimuli with orientation matching the orientation preference of the “index” L2/3-neuron defining those groups. Note that whether the L2/3 “index” neuron itself fires or not, does not alter substantially the stimulus-orientation prediction (Suppl. Fig. 58B vs. A), confirming information is contained in the group cofiring. We quantified the mutual information between stimulus orientation and various combinations of 1FC group-cofiring as well as single L2/3 neuron event firing. Fig. 7C histograms the mutual information conveyed by the cofiring of the sL4-1FC groups (green) and the sL2/3-1FC groups (blue) about the stimulus, clearly much higher than control (grey). Ensemble-to-ensemble outperforms ensemble-to-neuron information transfer during stimulus presentation, resulting in improved sensitivity and precision in predicting stimulus orientation, while the variance of the specificity and accuracy distributions decreases even though their means remain approximately unchanged (Suppl. Fig. 55). As expected, orientation-tuned L2/3-neurons by themselves convey much lower information (Suppl. Fig. 59A).

One question is whether the information about stimulus orientation contained in the cofiring of corresponding sL4-1FC and sL2/3-1FC groups is largely overlapping versus distinct and additive. Figs. 7D, 7E demonstrate that the mutual information conveyed by the joint probability of sL4-1FC and sL2/3-1FC group cofiring about the orientation of the stimulus is only slightly higher compared to the mutual information conveyed by either the sL4-1FC or the sL2/3-1FC group cofiring considered in isolation (Fig. 7D: example mouse; 7E: across mice), suggesting that they contain largely overlapping information. This is consistent with the possibility that these groups constitute successive stages of processing in an orientation-information transmission pathway.

## 3. Discussion

The ensemble-based cortical computation hypothesis posits that neuronal ensembles act as functional processing units. Yuste et al.^14^ argue that such ensembles arise intrinsically and are recruited spontaneously or by sensory input, echoing ideas from Lorente de Nó, Hebb, Hopfield, and others. Ensembles are thought to encode external sensory signals while also supporting internally generated representations, making the principles governing their organization and dynamics central to understanding cortical computation. Key open questions include how to robustly identify cooperative ensembles, what spatiotemporal rules govern their co-firing, how ensembles communicate across layers, how internal-state variables modulate ensemble dynamics, and whether these properties generalize across conditions (e.g., spontaneous vs. stimulus-driven activity; locomotion vs. quiet wakefulness). The functional correlation architecture of cortical microcircuits provides a valuable framework for exploring the algorithmic structure of cortical computations, complementing insights from anatomical connectivity. Here, we leverage this correlation structure to probe the organization of ensemble-level computations in cortical microcircuits. We mapped functional connectivity across granular and supragranular layers of mouse V1 to identify multi-neuronal ensembles operating within and across layers, and to compare their organization at rest versus during stimulus-driven conditions dominated by signal correlations. In what follows, we propose that first-order functional-connectivity groups of pyramidal neurons constitute tractable ensemble-level computational primitives—modules at an intermediate scale between single cells and whole-population activity—that support information processing within and across cortical layers.

### Pairwise Functional Connectivity Observations

We identified neuronal synchrony at the 2-photon imaging time scale of ∼158.7 ms/frame, using the STTC measure^43^. Even under a conservative threshold (z-score*>*4), 14-25% of pyramidal neuron pairs in layers 2/3 and 4 of mouse V1 showed significant positive correlations. A preponderance of significant pairs was seen in L4 (25.1%) compared to L2/3 (14.4%), consistent with the greater decorrelation characteristic of supragranular layers. Spatially, the distribution of significant positive pairwise correlations was concentrated within 300 *µ*m but extended up to 1mm, where 5-10% of pairs still exhibited significant functional connectivity (Suppl. Fig. 41J-K), indicating that local synchrony is embedded within a broader mesoscale network. Despite their statistical significance, pairwise correlation coefficients were uniformly weak in all layers (0.01-0.02 on average, ∼0.053 ±0.004 for significant positive correlations; Fig. 2B, Suppl. Fig. 7A-D), exhibiting only minimal dependence on distance (Suppl. Fig. 41K). Furthermore, they are similarly weak during both spontaneous and stimulus-driven activity (Suppl. Fig. 41), and remain weak across different aggregate activity epochs (Suppl. Fig. 22A-B). These findings align with prior reports of low noise correlations and weak unit-to-unit coupling in visual cortex^18,38,44,45^, supporting the view that weak positive pairwise correlations represent a universal feature of cortical circuitry. Notably, significant—though weaker—negative correlations were also detected, peaking at approximately 600–700 *µ*m from the soma (Suppl. Fig. 9); these were not examined further in the present analysis.

Interestingly, resting-state functional connectivity showed only a weak dependence on orientation preference (Suppl. Fig. 11B), in contrast to the much stronger dependence observed under stimulus-driven conditions (Suppl. Fig. 48). This dissociation suggests that spontaneous activity does not simply recapitulate stimulus-evoked correlation structure. Instead, the weak and across-orientation distributed structure of resting-state correlations is compatible with a network architecture that supports learning and generalization by preserving flexibility. Consistent with this view, V1 layers exhibit small-world organization under both spontaneous and stimulus-driven conditions (Suppl. Figs. 14,46, respectively) as well as robustness (Suppl. Figs. 15, 47).

Furthermore, the network topology changes across layers. Under resting state, L4 displays an approximately uniform hub-like topology that supports distributed processing (Suppl. Fig. 8C-D), compared to the more sparse organization of L2/3 (Fig. 2D, Suppl. Figs. 41C,I and 42). Interestingly, both within and across layers, the stimulus presentation condition is associated with reduced degree-of-connectivity and clustering-coefficients compared with spontaneous activity, indicating that functional organization is sparser in stimulus presentation (Suppl. Fig. 42). Together, these results argue that cortical computation is not mediated primarily by strong pairwise interactions but rather by the coordinated activity of neuronal ensembles embedded within weakly correlated networks^38,44^.

In this framework, information processing is best understood as the propagation of structured activity across ensembles—a principle we examine explicitly in the following sections.

### First-Order Functional Connectivity Response Properties

Resting-state functional connectivity analysis revealed firstorder functionally connected neuronal ensembles that serve as candidate computational primitives mediating communication across cortical layers. These ensembles exhibit markedly elevated clustering coefficients relative to controls (Fig. 2F, Suppl. Figs. 8B, 42D-F) indicating cooperative organization consistent with multi-neuronal computational modules. Crucially, statistically significant co-firing in large ensembles does not require strong pairwise correlations^46^; even weak pairwise dependencies are sufficient (Suppl. Fig. 10). This suggests that the weak pairwise “functional interactions” observed here can still give rise to population-level events that encode and transmit information.

Guided by this observation, we examined how aggregate co-firing within Layer 4 first-order functional-connectivity groups (L4-1FCs) relates to the activity of downstream L2/3 neurons. Across the population, the firing probability of L2/3 pyramidal neurons was well captured by rectified linear unit (ReLU)-like input–output relationships as a function of their corresponding L4-1FC group co-firing, with some neurons reaching firing probabilities approaching unity (Fig. 3A-3C). This functional form is notable, as ReLU nonlinearities confer key computational advantages—including sparsity, efficient learning, and representational efficiency—in both biological and artificial neural systems^47,48^. For neurons whose responses were well-fit by a ReLU (*R*^2^ ≥ 0.8), on average 70 aggregate co-firing events were sufficient to drive firing with probability approximately 1 (Fig. 3C), consistent with prior theoretical estimates^49^.

Strikingly, the slope of L2/3 response functions scaled inversely with the size of the corresponding L4-1FC groups: neurons associated with smaller groups exhibited steeper input–output functions, whereas those linked to larger groups showed shallower slopes (Fig. 4A). Control analyses ruled out potential imaging artifacts, such as partial sampling of larger groups extending beyond the imaging plane (Suppl. Figs. 23,26). The inverse relationship between response slope and L4-1FC group size is best explained by adaptive matching of the L2/3 neurons’ dynamic range to the statistics of their aggregate L4-1FC group input. We found that when cofiring was rescaled by *N* ^−*α*^, where *N* is L4-1FC group size, and *α* is consistently 0.6–0.7 across mice, L2/3 response functions collapsed onto a common curve (Fig. 4B). Under the assumption of independence, aggregate L4-1FC input fluctuations would scale as *N* ^−0.5^; the slightly larger observed exponent is consistent with the existence of positive pairwise correlations within L4. Importantly, retaining the identity of individual neurons within L4-1FC groups added little power beyond aggregate cofiring in predicting the activity of the associated L2/3 neuron. Specifically, classifiers incorporating neuron identity did not outperform models based solely on total co-firing across multiple algorithms (Linear-Kernel SVM: Fig. 3I; Random Forest, Naive Bayesian, and Logistic Regression classifiers: Suppl. Fig. 19), supporting the view that ensemble-level aggregate cofiring activity—rather than specific neuron identities—drives downstream responses.

Furthermore, group size appears to index functional pathways with distinct properties. Neurons with small L4-1FC groups fired more independently from each other (Suppl. Fig. 17A) and were weakly modulated by pupil diameter (Fig. 4C) and global population activity (Fig. 4D), resembling the “soloists” described by Okun et al.^50^. In contrast, L2/3 neurons associated with larger L4-1FC groups showed stronger modulation by these state variables (Fig. 4C-4D), akin to “choristers.” These phenotypes form a continuum rather than a binary classification. Across this continuum, L2/3 neurons fired sparsely: response probability increased sharply only for the largest L4-1FC events, approximately beyond the 93rd percentile of aggregate co-firing (Fig. 3G), dependent only weakly on group size (Suppl. Fig. 20). Consequently, only a small fraction of ensembles were strongly active in any given frame (Suppl. Fig. 58D), consistent with a coding regime that balances computational efficiency with reliable signal transmission. Notably, response slopes remained stable across fluctuations in behavioral state—including changes in pupil size and population activity—regardless of L4-1FC group size (Suppl. Figs. 24,28), indicating that L4-L2/3 coupling follows stable input–output rules despite internal-state variability. Moreover, small and large L4-1FC groups did not differ systematically in spatial extent or orientation tuning composition (Suppl. Figs. 18, 49H), indicating that group size does not simply reflect differences in visual field coverage or feature selectivity.

### Ensemble-to-Ensemble Functional Coupling

While the preceding analyses focused on ensemble-to-neuron coupling, information transmission in cortex is likely mediated by ensemble-to-ensemble interactions^1,7,36,39^. In this context, intra-L2/3 first-order functional-connectivity (L2/3-1FC) groups constitute natural downstream recipients of L4-1FC group input, since: **i)** L2/3-1FC ensembles exhibit high clustering coefficients (Suppl. Fig. 8B) suggesting strong cooperativity, **ii)** event-triggered average (ETA) analysis suggests functionally connected L2/3 neurons share significant common input from L4 (Suppl. Fig. 32), and **iii)** inter-layer L4-1FC domains of L2/3-1FC ensemble members exhibit much higher overlap than expected (Fig. 5B). Consistently, ensemble-to-ensemble coupling between L4-1FC and L2/3-1FC groups outperformed ensemble-to-neuron models across multiple metrics, including mutual information, sensitivity, precision, accuracy, and specificity (5C; Suppl. Fig. 31), identifying these coupled ensembles as putative functional information pathways. Notably, corresponding L4-1FC and L2/3-1FC ensembles convey highly overlapping stimulus information (see below and Fig. 7E), consistent with their hypothesized role as interlaminar channels for information transmission.

### Resting-State versus Stimulus-Driven Functional Connectivity Architecture

During sensory stimulation, correlations are expected to reflect shared stimulus drive (“signal correlations”) in addition to internally generated dynamics, leading to a reorganization of functional connectivity. Consequently, in our experiments, only 27-35% of functional connections identified at resting state remained significant under stimulation conditions (Suppl. Figs. 51A,43A), irrespective of whether neuronal pairs were tuned to orientation or not (Suppl. Fig. 51D, 43B). Moreover, under stimulus-driven conditions, functionally connected neurons exhibited a markedly stronger bias toward sharing similar orientation tuning (Suppl. Fig. 48), reflecting the dominance of feedforward, feature-specific inputs during sensory encoding. This is a sharp contrast to the observation that neurons functionally connected at resting state exhibit only a weak orientation tuning similarity (Suppl. Fig. 11).

It has been proposed that spontaneous cortical activity partially reflects the replay of patterns experienced during sensory stimulation^40^. Our data are consistent with the presence of such a tendency but indicate that it is relatively weak: neuronal pairs that exhibit strong coactivation during stimulation—based on shared orientation tuning—show at best a modest bias in functional connectivity (Suppl. Fig. 11B) under resting-state conditions. Rather, in the absence of sensory drive, functional connectivity appears to be more promiscuous, a property that likely reflects the latent capacity of cortical neurons to form associations rather than the expression of specific stimulus representations. That is, under spontaneous conditions, neurons are not constrained to participate in a single feature-defined ensemble, but instead remain broadly coupled to multiple partners with diverse functional properties. Such “promiscuity” may serve two complementary roles: first, enabling individual neurons to participate in multiple, partially overlapping ensembles that are selectively recruited under different stimulus or task contexts; and second, preserving the flexibility required for experience-dependent plasticity by keeping synaptic and ensemble memberships flexible rather than rigidly committed. In this view, spontaneous functional connectivity reveals not just the realized coding structure of the circuit, but its potential for forming meaningful associations as demanded by future sensory experience, generalization, or learning.

### Ensemble Level Principles of Communication are Conserved under Stimulation

Despite this substantial reconfiguration at the level of individual pairwise correlations, the ensemble-level organization and communication rules identified at rest were preserved when functional connectivity was redefined within the stimulus epoch. Specifically, when *stimulus-defined* first-order functional connectivity groups (sL4-1FC and sL2/3-1FC) were used, L2/3 neuron responses as a function of aggregate sL4-1FC group co-firing followed the same rectified linear (ReLU-like) input–output relationship observed at rest, including the same dependence on sL4-1FC group size (Figs. 6A-6C; Suppl. Figs. 53-56). Consistent with these results, stimulus-defined ensemble-to-ensemble communication from sL4-1FC to sL2/3-1FC groups likewise exhibited robust information transfer properties (Figs. 7B-7E; Suppl. Figs. 55). As in the resting state, aggregate ensemble cofiring consistently performed at least as well as models incorporating individual neuron identity, indicating that precisely identifying which neurons fire adds little, if any, explanatory power beyond ensemble-level activity for predicting L2/3 responses (Suppl. Fig. 56), or the identity of the stimulus (Suppl. Fig. 59B-E). This finding suggests that aggregate, population-level, coordination at the ensemble level, rather than the identity of individual participating neurons, dominates cortical information transmission between layers during both spontaneous activity and stimulus-driven conditions.

In sum, all key response properties of L2/3 pyramidal neurons as a function of the cofiring of sL4-1FC ensembles—namely the ReLU-like nonlinearity, the existence of weak and strong response regimes, and the adaptation of response gain to the dynamic range of sL4-1FC activity—held across stimulus presentation conditions (Suppl. Figs. 53-56), as well as across shorter epochs and distinct brain states, e.g., locomotion vs. quiet wakefulness (Suppl. Figs. 36-38) or high vs. low alertness (Suppl. Fig. 29). This suggests that ensemble-level input–output rules reflect intrinsic constraints of cortical circuit organization, rather than stimulus- or state-specific specializations.

Stimulus-driven conditions further allowed a direct test of the representational significance of first-order functional connectivity modules in encoding stimulus orientation. Cofiring within sL4-and sL2/3-1FC ensembles associated with orientation-selective L2/3 “index” neurons accurately predicted stimulus orientation (Fig. 7A), conveying substantially more mutual information about stimulus orientation than control groups matched for size (Figs. 7C,7E).

At the circuit level, ensemble-to-ensemble transmission from L4 to L2/3 consistently outperforms ensemble-to-neuron models in sensitivity, precision, accuracy, and specificity, supporting a feedforward, ensemble-based framework for interlaminar communication. We propose that corresponding L4 and L2/3 ensembles form information pathways, a view reinforced by their shared representational content: the mutual information between stimulus orientation and L4 ensemble cofiring is comparable to that between stimulus orientation and L2/3 ensemble cofiring and is not substantially increased when considering their joint activity. This equivalence indicates that L4 and L2/3 ensembles encode largely overlapping stimulus information, consistent with a pathway architecture in which information is relayed across layers.

These results align with prior evidence that population-level coordination, rather than single-neuron activity, underlies robust sensory encoding in cortex^9,18,19^ and go further in identifying candidate ensembles that are important for information transmission and warrant further investigation. While these findings do not establish causality, they motivate future experiments combining functional connectivity analysis with targeted perturbations to directly test the computational role of these ensembles.

### Relating Biological and Artificial Neural Network Principles

The organizational principles uncovered here appear to extend beyond biological circuits. Applying the same analytical framework to the activity of hidden units in deep neural networks—including spiking-network extensions of ResNet18, ResNet32 and VGG16 pretrained on CIFAR-100—during exposure to a set of inputs, revealed analogous ensemble-level structure and nonlinear input–output relationships^51,52^. Although these comparisons are necessarily conceptual rather than quantitative, they suggest that stimulus-based functional connectivity analysis captures encoding-relevant organizational features that may reflect intrinsic constraints on information flow in networks optimized for information processing.

The ensemble architecture observed in both systems is characterized by overlapping, distributed representations, conceptually related to the principle of superposition recently proposed as an efficient coding strategy in artificial networks^53^. Such overlap allows individual units to participate in multiple computational pathways, paralleling mixed selectivity in cortical neurons, which has been shown to support flexible, contextdependent computations^54^. In prior work, we formalized the information capacity of such ensemble-based pathways under simplifying assumptions^39^, providing a theoretical basis for how overlapping representations can balance capacity, robustness, and flexibility. Consistent with this view, incorporating analogous modular pathway structures into sparse feedforward artificial networks preserved comparable task performance to dense architectures while substantially reducing computational cost^55^.

Together, these observations support the idea that ensemble-based communication and overlapping representations may constitute a general computational motif shared across biological and artificial systems, even if implemented through distinct mechanisms. While preliminary, this convergence suggests that functional connectivity–based analysis is a useful level of abstraction for testing neurocomputational hypotheses linking circuit organization, efficiency, and flexibility, while providing powerful tools for interpretability in neuroAI models.

### Temporal Window Dependency on Functional Connectivity

Systematic variation of recording length further revealed that shorter imaging sessions yield fewer, but stronger and more reproducible, functional connections (Suppl. Figs. 3A,13). This observation suggests the existence of a relatively stable “core” functional-connectivity scaffold—comprising connections that persist even in brief time windows—onto which weaker, more transient interactions can be flexibly added or removed (Suppl. Figs. 6, 13, 39, 45). Such a two-tier architecture is well suited to support adaptive learning: a stable backbone that preserves computational integrity, coupled with a dynamic periphery that enables rapid reconfiguration and, later, consolidation. This view is consistent with recent reports of persistent neuronal ensembles that maintain long-term representations of perceptual states or memories^56^.

### Limitations and Sensitivity Analysis

The temporal resolution of calcium imaging (∼158.7 ms per frame) precludes inferring causality or resolving fast synaptic interactions at millisecond timescales. Nevertheless, within these constraints, our analyses proved robust and sensitive enough to recover expected circuit asymmetries, including a bias favoring feedforward L4-to-L2/3 interactions (Suppl. Fig. 16B). Importantly, while the absolute number of statistically significant functional connections varied with parameter choices (Fig. 2D), the core organizational principles reported here were preserved across a broad range of parameters. These included variations in the STTC integration window (Δt = 0 or 2 frames, Suppl. Fig. 2), significance thresholds, correlation metrics (Pearson versus STTC; Suppl. Figs. 4B), removal of global state components linked to population activity and pupil size (Suppl. Fig. 33), and overall imaging duration (Suppl. Fig. 3).

Finally, we note that the relationship between functional connectivity and its underlying anatomical substrate remains an open question. However, this ambiguity may be less limiting than often assumed. Cortical computation is implemented through dynamic patterns of activity that can diverge substantially from static anatomical wiring, particularly as neuromodulatory state, experience, and task demands change. In this context, functional connectivity provides a complementary—and in some cases more direct—window onto the emergent organization and algorithmic flow of information in neural circuits, even when the precise anatomical pathways are complex or incompletely known.

## 4. Conclusions

Our findings support and extend the view that cortical computations are organized around multi-neuronal, ensemble-based primitives. We show that first-order functionally connected ensembles can serve as basic, multiplexed, computational modules that mediate information processing within and across cortical layers. These ensembles exhibit strong cooperativity, such that aggregate cofiring, rather than individual neuron identity, predicts downstream neuronal responses through a robust, ReLU-like input–output relationship. Importantly, the gain of this relationship scales systematically with ensemble size, consistent with adaptive tuning to the dynamic range of incoming inputs. We further demonstrate that internal brain state modulates ensemble dynamics in a graded, ensemblesize–dependent manner, spanning a continuum from weakly coupled “soloist” neurons to strongly population-coupled “choristers”, highlighting the functional diversity of these modules.

Together, these findings establish first-order functionally connected ensemble cofiring as a functionally meaningful substrate for sensory encoding and interlaminar information transfer. While multiple approaches have been proposed to identify computationally meaningful neuronal ensembles^14,57,58^, our functional-connectivity-centric framework offers a simple, scalable and experimentally tractable means for interrogating the algorithmic principles of cortical computations. In contrast to prior studies that either lacked Layer 4 recordings or were more limited in scale^16,19,56,57,59^, our approach enables a more extensive interrogation of interlaminar communication at the ensemble level. The resulting picture is one of flexible, multiplexed, ensemble modules whose interactions can be dynamically reconfigured to support task-specific processing, adaptation, and learning.

Looking forward, combining this framework with causal perturbations will be essential for establishing the mechanistic role of specific ensembles in cortical computation, perception, and learning, as well as for understanding how these principles break down in disease. Beyond neuroscience, the organizational motifs identified here—overlapping, ensemble-based communication with adaptive gain control—may also inform the design of biologically-inspired artificial intelligence systems and neuro-realistic digital twins, providing a foundation for testing neurocomputational hypotheses across biological and artificial domains in the future.

## 5. Material and Methods

### Experiments and Data Collection. Mouse Lines and Surgery

All procedures are approved by the Institutional Animal Care and Use Committee (IACUC) of Baylor College of Medicine. 5 mice, 0-12 weeks of age, expressed GCaMP6s in excitatory neurons via SLC17a7-Cre and Ai162 transgenic lines (JAX stock 023527 and 031562, respectively) cross. Animals were anaesthetized and a 5 mm craniotomy was placed over visual cortex. Each mouse recovered for ∼2 weeks before the first experimental imaging session.

### 2-photon Imaging

Mice were head-mounted on a treadmill and calcium imaging was performed using Chameleon Ti-Sapphire laser (Coherent) tuned to 920 nm and a large field of view mesoscope equipped with a custom objective (0.6 NA, 21mm focal length). Laser power after the objective increased exponentially as a function of depth from the surface according to: *P* = *P*0 × *e*^(*z/Lz*)^, where *P* is the laser power used at target depth z, *P*0 is the power used at the surface (not exceeding 15mW), and *Lz* is the depth constant (not less than 220*µ*m.) Maximum laser output of 90 mW was used for scans approximately 450 *µ*m from the surface and below.

### Monitor Positioning and Retinotopy

Visual stimuli were presented to the left eye with a 31.1 ×55.3*cm*^2^ (*h*× *w*) monitor (resolution of 1440 × 2560 pixels) positioned 15 cm away from the mouse eye. Pixelwise responses across 2400 × 2400*µm*^2^ to 3000 × 3000*µm*^2^ region of interest (0.2 px/*µ*m) at 200-220*µ*m depth from the cortical surface to drifting bar stimuli were used to generate a sign map for delineating visual areas^60^. We chose an imaging site spanning all primary visual cortex visible within the craniotomy and a fraction of the adjacent medial and lateral higher visual areas. Imaging was performed at ∼6 Hz for all scans, collecting eight scanfields at 0.6 px*/µ*m along a cortical column (2 adjacent fields per layer). The two adjacent fields of view were 1200*µ*m × 600*µ*m, with an overlapping of 15*µ*m, for each of the 5 mice examined. Across 2-5 sessions, a scan was collected at each target depth by manually matching reference images to target depth within several microns using structural features including horizontal blood vessels (which have a distinctive z-profile) and patterns of somata (identifiable by GCaMP6s exclusion as dark spots). Imaging data were motion-corrected, automatically segmented and deconvolved using the CNMF algorithm^26^; cells were further selected by a classifier trained to detect somata based on the segmented cell masks. This resulted in ∼7000-8000 soma masks per animal per column. The retinotopic mapping that was performed is described in the method section of Bae *et al*.^22^.

### Directional Visual Stimulus

A stimulus using smoothened Gaussian noise with coherent orientation and motion was used to probe neuronal orientation and direction tuning. An independently identically distributed (i.i.d.) Gaussian noise movie was passed through a temporal low-pass Hamming filter (4Hz) and a 2-d Gaussian filter (*σ* = 4.4^?^ at the nearest point on the monitor to the mouse). Each scan contained 72 blocks, with each 15-second block comprising of 16 equally distributed and randomly ordered unique directions of motion between 0-360 degrees with a velocity of 42 degrees/s at the nearest point on the monitor. An orientation bias perpendicular to the direction of movement was imposed by applying a bandpass Hanning filter G(*ω*; c) where *ω* is the difference between the image 2d Fourier transform polar coordinates *ϕ* and trial direction *θ*, and

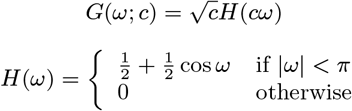

Here, c = 2.5 is an orientation selectivity coefficient. The resulting kernel is 72^?^ full width at half maximum.

### Direction/Orientation selectivity

The directional trial response was measured by taking the difference in cumulative deconvolved activity at the linearly interpolated trial onset and offset time points. Trial responses per direction were modeled as a two-peak scaled von Mises function in the form:

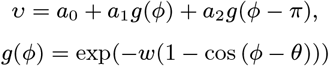

where *θ* is the preferred direction, *ϕ* is the trial direction, *w* is the peak concentration, *a*0 is the baseline, and *a*1, *a*2 are the independent amplitudes of two peaks. The two peaks share a preferred orientation, baseline, and width, but their amplitudes are fit independently. This function was fitted to minimize the mean squared error of all trial responses across 16 directions using the L-BFGS-B optimization algorithm^61^. Significance and goodness of fit were calculated by permutation. Trial direction labels were randomly shuffled among all trials for 1000 refits. The goodness of fit was calculated as the difference in fraction variable explained (FVE) between the original fit FVE and the median FVE across all 1000 shuffled fits. The p-value was calculated as the fraction of shuffled fits with a higher FVE than the original fit. Neurons were included for further network analysis if p-value ≤0.001 and the difference in FVE was *>* 0.025.

The neurons’ orientation and direction tuning were estimated as in Fahey *et al*.^62^. Briefly, responses to a dynamic stimulus of pink noise with coherent orientation and motion were fit with a two-peak von Mises function. Cells were included for further analysis by a dual threshold for the fraction of variance explained (*>*2.5%) and significance calculated by permutation *p* ≤ 0.001. In our sample, ∼46% of L4-neurons (2081 out of 4551) and ∼50% (7349 out of 14774) of L2/3-neurons were orientation selective using these criteria, consistent with^62^. All orientation selective units were then sorted “cyclically” into 128 bins according to the preferred direction of the larger amplitude von Mises peak. The cells that are oriented have p-value ≤ 0.001 and FVE *>* 0.025.

### Absolute Orientation Difference

For each pair of neurons (n1, n2), we estimate the absolute direction difference of their strongest amplitude angle w1 and w2, respectively, *ϕ* as follows:

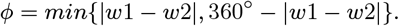

The corresponding absolute orientation difference *ω* of their strongest amplitude is equal to *ϕ*, for *ϕ* equal or less than 90^?^ otherwise, it is equal to 180^?^ − *ϕ*.

### Receptive Field Structure and Similarity

Receptive field maps were calculated using spike-triggered pixel-wise average (STA) of the monet movie frames preceding the spike event in the neuron by 250 ms (spike-triggered activity integration window [-250 0] ms). The resulting STA maps were filtered with 2d Gaussian filter using 10 * 10 pixels kernel. The maximum response location was set within the 95% of the Gaussian fit amplitude (receptive field region of interest (RF ROI)). Signal-to noise ratio (SNR) was determined as a ratio of variance within the receptive field ROI and variance of the background (all pixels of the map outside Gaussian fit for the receptive field). Only cells with SNR≥ 2 were considered to have defined visual receptive fields and accepted for further analysis. For accepted cells (SNR ≥2), the STA maps were converted to z-score maps (using map mean and standard deviation to assign z-score to every pixel). Next, the values [-2 2] were set to 0. For the resulting maps, we calculated the pairwise structural similarity index (SSIM^63^), indicative of the 2d-correlation between the on- and off-field structure of the two maps. Next, we explored the relationship between SSIM and STTC values between neurons.

### Software

Experiments and analyses were performed using custom software developed using the following tools: ScanImage, CaImAn, DataJoint^64,65^, PyTorch NumPy^66^, SciPy^67^, Docker, matplotlib^68^, cmocean, Jupyter, pandas, scikit-learn^69^, multiprocessing.

### Deconvolution

The fluorescence signal recorded through two-photon microscopy from neurons is deconvolved to obtain an estimated spike train as described in Pnevmatikakis *et al*.^26^. The neurons recorded express the GCaMP6s indicator. The Pnevmatikakis *et al*.^26^ method simultaneously identifies individual neurons, demixes spatially overlapping neurons, and finally deconvolves the fluorescence signal to produce probability amplitudes for the existence of spikes. The method involves constrained non-negative matrix factorization of fluorescence into time and space components, and finally an autoregressive deconvolution process for determining the (denoised) calcium transient and the spike probability amplitudes. In our work, the autoregressive model used is a two-step autoregressive model AR(2).

### Thresholding

We use raw fluorescence minus the background to find the “noise intervals”, which are the frames where the aforementioned fluorescence has negative values. The deconvolved signal restricted on the noise intervals is called *Snoise*. In the case of *Snoise* with more than 3 nonzero values, we evaluate the 68th percentile of the nonzero noise values distribution. This is meant to give the scale of the noise, and multiples of this are used as thresholds for determining spikes (indicated as *dc* thresholds). Note that the 68th percentile corresponds to the standard deviation in the normal distribution. The 68th percentile is used, instead of evaluating the standard deviation, to avoid an increase in the standard deviation due to outlier values. We found that spike determination is not very sensitive to threshold variations if we choose the threshold to be 1.5*(68th percentile of noise). In the case of *Snoise* with 3 or less non-zero values (*>* 0.001), all *Snoise* ‘s nonzero values become 0s and the rest 1s. This is justified by the fact that although there are noise intervals long enough, there are too few nonzero *Snoise* frames, suggesting that for these neurons there is very little noise. Note that less than or equal to 3 non-zero values for *Snoise* are not sufficient to permit characterization of noise (evaluation of 68th percentile of noise). This can be further tested by comparing the distributions of the firing rates of two neuronal populations, namely the one with *Snoise* of 3 or less non-zero values, and the remaining ones. For the second population, the noise standard deviation has been estimated through the 68th percentile of *Snoise*. For these neurons, the cutoff has been taken to be 1.5*(68th percentile of noise).

### Functional-connectivity Analysis Methods

**STTC** To quantify functional-connectivity, it is important to use a robust temporal correlation measure. Here, we use the spike time tiling coefficient (STTC), introduced by Cutts and Eglen^23^, which performs favorably against 33 other commonly-used measures and is less sensitive to firing rate.

The STTC of neuron A relative to B is defined as:

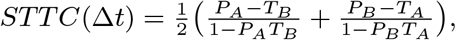

where *PA* is the proportion of firing events of neuron A found within an interval (± Δt) around each firing event of neuron B, *TB* is the proportion of the recording duration that falls within an interval (±Δt) around each firing event of neuron B, and likewise for *TA* and *PB*. STTC has been symmetrized and normalized to lie within [-1, 1]. The bulk of the results presented below use STTC values measured at Δt=0, i.e., the probability that neurons fire in synchrony within one calcium imaging frame, for which the two measures are identical. Since the calcium frames last ∼158.7ms in our experiments, the order of events with a temporal difference less than that cannot be distinguished, so the temporal synchrony correlation STTC measure we compute is insensitive to temporal differences smaller than ∼158.7ms. We also computed STTCs by taking Δt= ∼300ms (i.e. broadening the window of synchrony to 2 additional imaging frames; Suppl. Fig. 2 A-B), in which case we used an extension of the STTC measure we devised to take into consideration the temporal order of occurrence of spikes in A relative to B, i.e. reflect the probability that spikes of one neuron may systematically precede (or follow) spikes of the other.

### Estimation of the null in STTC

To evaluate the extent to which the observed STTC values could arise by sequences with the same number of firing events and inter-event intervals but without any temporal structure, we circularly shifted the firing events of each neuron by a uniformly sampled integer number of imaging frames within the interval 500 times independently. From these iterations, we obtained a null distribution of STTC values for each pair of neurons. The *z-scored* STTC weight of the edge between neurons (*i, j*) (say *wi,i*) is defined as (*wi,j* −*m*0)*/σ*0 where the *m*0 and *σ*0 are the mean and standard deviation of the corresponding *null* distributions of the pair (*i, j*). The z-score quantifies the distance of the observed STTC value from the mean of the null STTC distribution in units of standard deviations of the null STTC distribution.

### Statistical Significance of the STTC in each Layer

We identify the positive inter-neuronal functional correlations (“edges”) with z-score *>* 4 in each layer, considering neurons located at the same layer (L2, L3, and L4). In the histograms, we merge the L2 and L3 distributions (union of edges), and report it as L2/3. Thus, L2/3 includes only *intra-layer* edges, between pyramidal neurons that belong at the same layer. Similarly, we compute the negative inter-neuronal functional connections at different z-score levels.

### Functional-connectivity and Bias in Tuning Function

We compute the bias in tuning function, by considering all neuronal pairs in a specific layer case (e.g., intra-layer L4 or L2/3 or inter-layer from L4 to L2/3). For those neuronal pairs, we estimate the percentage of statistically significant edges (z-score *>* 4) that have absolute orientation difference within a certain interval (bin) (say, e). Similarly, for the aforementioned neuronal pairs, we form the null, which is the set of functional connections edges with z-score in [-2, 2] and identify the percentage of them that belong in that bin (i.e., n). We then report the difference computed in each bin normalized by the probability of the null in that bin ((*e* − *n*)*/n*).

### Estimation of the probability of firing of a recipient neuron

The probability of firing of a recipient L2/3-neuron as a function of the number of its incoming functionally-connected L4-neurons that fire simultaneously during a frame is shown in Fig. 3A, with bins (at the x-axis) for each number of cofiring events. To profile the probability of firing of the putative recipient L2/3-neurons, we then apply weighted linear regression on these data points, as well as a weighted ReLU fit. To improve the reliability of regression, we first merged the original bins to ensure that each bin has a sufficient number of frames (here at least 10 frames) as follows: For each recipient neuron, we start from the rightmost (largest) bin (i.e., [N, N+1)), where *N* is the maximum number of cofiring events of its incoming functionally-connected L4-neurons, and iteratively “scan” the bins from right to left merging bins to ensure that each merged bin has at least 10 frames. We only merge consecutive bins as follows: **1)** examine the number of frames at the current bin and if it has less than 10 frames, **2)** merge it to the next bin at the left (creating a “super-bin”). If the newly-merged bins (super-bin) has still less than 10 frames, we continue the bin-merging: merge the super-bin with the next bin at the left until the set of consecutive bins (super-bins) contain *at least 10 frames*. We then continue with the scanning as described above with the next bin. The center of the merged bins corresponds to the (rounded) weighted average of the original bins, with *weights based on their number of frames*. For each recipient neuron, we perform *weighted linear fitting and weighted ReLU fitting* on its data points, of the form (cofiring events, firing probability), for co-firing events in the range of [0, N]. The weight of the i-th point (bin) is based on the standard error of the mean in that bin (normalized by the maximum standard error of the mean (SEM) observed in that recipient neuron, across all its points). In the case of the percent of cofiring events, the bins used are of the form [0, 0.5), [0.5, 1.5), …, [99.5, 100] percent of cofiring events.

### Estimation of the Control Input Neuronal Group of a Putative Recipient Neuron

We randomly select neurons of the same layer and of *equal size as the observed input neuronal group of interest* that does *not* overlap with it. If a control group cannot be formed due to the large size of the observed group, the corresponding index neuron is excluded from the control data (while its observed group is still plotted).

### Weighted ReLU fitting

For each recipient neuron, the weighted ReLU fitting is performed on the points (x, y) (namely, the number of cofiring events of the neurons of the 1FC group and the firing probability of the recipient neuron, respectively) after bin merging, for cofiring events in the range [0, N], where N is the maximum number of cofiring events of the 1FC group of the recipient neuron. Each point is weighted based on the standard error of the mean of the firing probability in the corresponding bin and normalized by the maximum standard error of the mean observed across bins of that recipient neuron The ReLU fit is composed of two parts, namely a constant/fixed part (parameterized by the plateau intercept and the plateau length) and a linearly increasing one (parameterized by the slope and the line intercept). The plateau intercept is the intersection point of the horizontal part with the y-axis. The plateau length is the point in the x-axis where the horizontal part stops and the linear part begins. The value for the line intercept is estimated from the values of the other parameters, to maintain continuity. The other three parameters are estimated using a subspace-bounded implementation of the least squares algorithm, so that the parameters are restricted in specific value ranges.

### Weighted Linear Regression

For each recipient neuron, the weighted linear regression is performed on the points (x, y), namely, the number of cofiring events of the neurons of the 1FC group and the firing probability of the recipient neuron, respectively, after bin merging, for cofiring events in the range [0, N], where N is the maximum number of cofiring events of the 1FC group of the recipient neuron. Each point is weighted based on the standard error of the mean of the firing probability in the corresponding bin and normalized by the maximum standard error of the mean observed across bins of that recipient neuron. The slope and the intercept of the linear fit are then estimated using the weighted least squares algorithm.

### R2

The coefficient of determination is the proportion of the variation in the dependent variable that is predictable from the independent variable(s). It provides a measure of how well observed outcomes are replicated by the model.

#### Unweighted

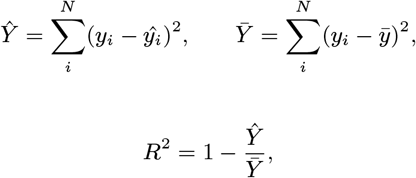

where *Ŷ* is the sum of squares of residuals, 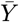 is the total sum of squares, *i* is the i-th element, *ŷi* is the model’s prediction for the i-th element, *yi* is the observed value for the i-th element, and 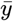 is the mean observed value.

#### Weighted

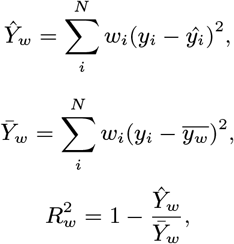

where *Ŷw* is the sum of weighted squares of residuals, 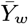 is the total sum of weighted squares, i is the i-th element, *ŷi* is the model’s prediction for the i-th element, *yi* is the observed value for the i-th element, *wi* is the weight assigned to the i-th element, and 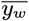 is the weighted mean of the observed values.

### RMSE

The root-mean-square error (RMSE) is a measure of the differences between values (sample or population values) predicted by a model or an estimator and the values observed and is therefore a measure of the model’s accuracy.

#### Unweighted

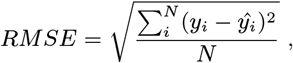

where i is the i-th element, *ŷi* is the model’s prediction for the i-th element, *yi* is the observed value for the i-th element, and *N* is the total number of elements.

#### Weighted

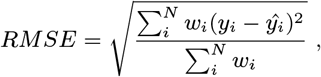

where i is the i-th element, *ŷi* is the model’s prediction for the i-th element, *yi* is the observed value for the i-th element, *wi* is the weight assigned to the i-th element, and *N* is the total number of elements.

### Normalized Root-Mean-Square Error (NRMSE)

The normalized root-mean-square error (NRMSE) is equal to the RMSE normalized by the value range of the values observed. It is a metric used for comparing models with different scales.

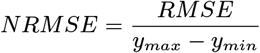

where *RMSE* is the root-mean-square error, *ymax* is the maximum observed value, and *ymin* is the minimum observed value. The *RMSE* used can be either weighted or unweighted.

#### Support Vector Machine Classification

For each L2/3 recipient neuron, we predict its firing under spontaneous conditions using as input the number of cofiring events of its L4-1FC group and the calcium eventogram of each individual neuron in its L4-1FC group. The first 70% of the hour-long recording was used for training. The three performance metrics, namely, sensitivity, specificity and accuracy, were computed on the remaining dataset (30%). Note that the L4-1FC groups were computed on the first 40 minutes of recording (approximate the dataset used for training). Only neurons with L4-1FC groups larger than 15 were considered. During training, we excluded a fraction of the frames where the L2/3 neuron does not fire in order to ensure that we have an equal number of frames in the two classes (i.e., firing events vs. silence). We also repeated this process under stimulus presentation.

### Statistical Tests and Statistical Significance

All histograms report statistics that indicate the sample mean and standard deviation of the sample means across mice (n=5), while the error bars indicate the SEM across mice for each specific bin. For all histograms, the statistical tests used are the permutation of means, the Welch’s t-test and the ANOVA (F-test) and the statistical significance reported results from the statistical test with the highest p-value (in order to be more conservative). For example, for the comparison of the observed vs. null for a specific metric, the means of the observed and the null are calculated per mouse, and then using the 5 means from each distribution, the above statistical tests are performed across mice. For each panel, the level at which the significance test was rejected is shown in the plots using the following notation: empty or “n.s.”, if the null hypothesis of equal means is not rejected (p-value *>* 0.05); “*”, if the null hypothesis is rejected for a significance level of 0.05 (p-value *<* 0.05); “**”, if the null hypothesis is rejected for a significance level of 0.01 (p-value *<* 0.01); and “***”, if the null hypothesis is rejected for a significance level of 0.001 (p-value *<* 0.001).

### Degree of Connectivity (DoC) and Clustering Coefficient (CC

For each neuron, we estimate the fraction of the neurons with statistically significant functional connections (z-score *>* 4), i.e., 1FC neighbors, per layer case, which corresponds to the *normalized* degree of connectivity in that layer. For example, a L4-neuron has an intra-layer degree of connectivity of 0.1 if that neuron is functionally-connected with the 10% of the L4-neuronal population. Moreover, we estimate the fraction of neuronal pairs of its 1FC neighbors that form statistically significant functional connections with each other, considering *only* the frames at which the index neuron does not fire, to account for possible bias. This corresponds to the clustering coefficient of the neuron. For example, an L2/3-neuron with a L4 →L2/3 clustering coefficient of 10% indicates that exactly 10% of all possible pairwise functional connections between its L4 neighbors are statistically significant.

### Small-worldness

A network is characterized as small-world-like when it has short paths between nodes and high clustering. In order to quantify the degree to which the observed networks exhibit small-world characteristics, we compare the average shortest path length and the clustering coefficient of the observed graphs to those of theoretical graphs^70,71^. The observed functional networks were formed considering all neurons within the layer of interest and their statistically significant functional connections (STTC z-score *>* 4). Each theoretical graph was constructed using the same number of neurons as their corresponding observed functional network.

For the **Erdős–Rényi** networks, *ER*(*N, p*), where *N* is the total number of neurons in the layer of interest and *p* the probability that two neurons are functionally-connected (equal to the mean normalized degree of the respective biological network), we placed an edge between each pair of neurons with probability *p*.

The **regular ring** graphs were created by assigning a fixed degree of connectivity to each of the N nodes. Specifically, the nodes are placed on the “ring”, in a circular topology and each node has a degree of connectivity equal to 2*k*, with *k* connections to its k nearest neighbors on its left and *k* connections with its nearest neighbors on its right. We then compare the three graphs concerning the shortest paths between pairs of neurons and the clustering coefficient.

### Small World Index

Another way to assess the small-worldness of a network is the Small World Index (SWI), a value between 0 and 1 which is higher for small-worldly graphs ands is defined as follows :

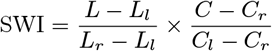

where *C* is the clustering coefficient, *L* is the path length, and *l,r* correspond to regular-ring and random graphs respectively.

#### Network Robustness

The networks of L4 and L2/3 are **robust** given the presence of a *giant component* that includes almost the entire neuronal population recorded at each corresponding layer. Note that a giant component is a sub-network that contains a large number of neurons (of the original network) and their functional connections. L2/3 follows a similar trend but up to a smaller z-score value.

The **Molloy-Reed criterion**^72,73^ 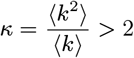 links the network’s integrity, as expressed by the presence or the absence of a giant component, to the mean degree of connectivity:

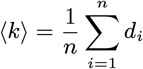

and its second moment:

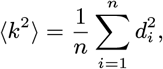

where *di* is the degree of connectivity of node *i* and *n* is the total number of nodes in the network. It is valid for any degree distribution *p*_*k*_^70^ (Chapter 8). In general, a network displays enhanced robustness, if its breakdown threshold (directly depending on *κ*) deviates from the random network prediction. The very high *κ* and high mean clustering coefficient, especially in L4 but also L2/3, significantly higher than the corresponding Erdős–Rényi graphs, suggest enhanced robustness (see Suppl. Figures 14, 46). In general, real networks are robust to random failures but fragile to targeted attacks.

### Normalization Process on the L4-1FC Group Cofiring for the Prediction of the L2/3-neuronal Response

In studying the response properties of L2/3-neurons with respect to their L4-1FC group, we searched for a normalization by a power of the group size, to have similar response curves irrespective of the group size. First of all, it is *not* at all obvious that such a normalization exists *over a wide range of group sizes*. Second, the input statistics depend on the group size, unless neurons fire independently and the input is normalized by the square root of the group size. Hence, if the input is normalized in such a way, the L2/3-neuronal response corresponds to the probability of the normalized input, hence the L2/3-neuron will know how rare cofirings are **irrespective of group size**. To conclude, one way to make the L2/3-neuron to respond according to the rareness of a cofiring, is *through the normalization of the input by the square root of the group size*. Below we present a simple theoretical model to show this normalization and provide an intuition:

Let’s suppose for simplicity that the L4 connectivity group consists of *N* independent neurons, each firing with probability *p* within one frame. In this case, the mean firing is *µ* = *Np* and the firing standard deviation is 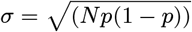. If we denote by *F* the L4 connectivity group firing random variable, then, by the central limit theorem, we have

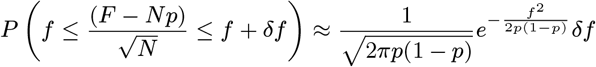

What is important about this distribution is that it is *independent of N*. Hence if L2/3-neurons have response probability proportional to the normalized group firing above the mean, 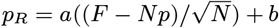, for the same constants *a, b*, they would respond under the same distribution *irrespective of the size of their L4 group*, permitting a more homogeneous response in L2/3. In this case, the slope of the L2/3-neuron response against their L4-1FC group cofiring normalized by *N* ^1*/*2^ would be the same irrespective of the L4-1FC group size. This is slightly less than the 0.6 −0.7 found experimentally. This difference might be attributed to either the existence of correlations between L4-neurons or differences in the probability of firing between different neurons in L4.

#### Noise Functional Connections under Stimulus Presentation

For each neuron (say *i*), we form its firing rate per segment time series (i.e., *ri*(t), for segment *t*) and estimate its mean firing rate per stimulus direction 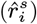, for stimulus direction *s*, estimated considering all segments during which the stimulus presented has direction *s*). The stimulus noise time series was formed per neuron by subtracting from the firing rate per segment time series (at segment t), the mean firing rate of that neuron for the corresponding stimulus direction (that is 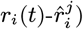), where *j* is the stimulus being presented at that segment *t*.

### Decoding Ability of 1FC Groups and Eventograms

We employed various machine-learning classifiers, such as Support Vector Machine (SVM), Logistic Regressor (LR), a Gaussian Naive Bayes Classifier (NB), and a Random Forest Classifier (RF), to predict L2/3-neuron firing using as input: i) the number of cofiring of its L4 1FC neighbors; ii) the number of cofirings considering its L4- and L2/3-1FC neighbors, and iii) the set of its L4-1FC neurons with their calcium event at each corresponding frame. Note that in the third case the input not only uses the number of cofiring events in the L4-1FC group but also the identity of the neurons that fire. Approximately the first 70% of the recording was used for training and for the identification of the 1FC groups. To ensure an equal number of frames in the two classes (i.e., calcium events vs. silence), a fraction of frames where the L2/3-neuron does not fire were excluded.

To assess the decoding ability of a given predictor (binary) variable in terms of a target (binary) variable, we define the following terms: TP (true positives), i.e., the number of frames where the predictor and target variables are both 1; false negatives (FN), i.e., number of frames where the predictor variable is 0 but the target is 1; FP (false positives), i.e., number of frames where the predictor variable is 1 but the target is 0; and TN (true negatives), i.e., number of frames where the predictor and target variables are both 0. Based on them, we employ the following classical machine-learning metrics, namely the *sensitivity (specificity)*, i.e., the proportion of positives (negatives) in the target variable that were correctly predicted, respectively, the *accuracy*, i.e., the overall correctness of the predictor variable over all frames, and the *precision*, i.e., the proportion of correct positive predictions.

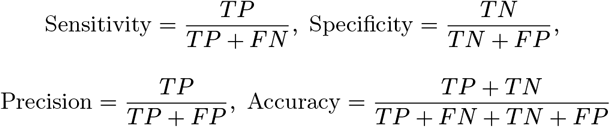

### Mutual Information

In information theory, the mutual information (MI) quantifies the amount of information that one random variable contains about another random variable. More specifically, it measures the reduction in uncertainty about one variable when the value of another one is known. Thus, high mutual information between two variables indicates a strong statistical dependence, which could imply a significant predictive relationship between them.

The mutual information of two jointly discrete random variables *X* and *Y*, which take values in the sets 𝒳 and 𝒴, respectively, is calculated as:

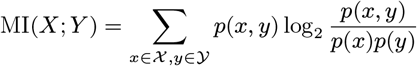

where *x* and *y* are index variables that represent each possible value of the random variable *X* and *Y*, respectively, *p*(*x, y*) is the joint probability distribution of *X* and *Y*, and *p*(*x*) and *p*(*y*) are the marginal probability distributions of *X* and *Y* respectively. ∑ denotes the sum over the possible values of the two random variables.

Mutual information can be measured in different units depending on the base of the logarithm. Base 2 gives units of bits (or “shannons”), base e gives units of nats (“natural units”), and base 10 gives units of “dits”, “bans”, or “hartleys”.

The normalized mutual information (nMI) between two jointly discrete random variables *X, Y* is calculated by dividing their mutual information by the average of their entropies. nMI values fall within the range of 0 to 1.

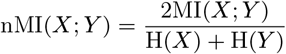

where H(*X*) is the entropy of X and H(*Y*) is the entropy of *Y*.

The joint mutual information measures the amount of information that two random variables jointly provide about another random variable, when considered together. Let *X*1, *X*2, and *Y* be three jointly discrete random variables, which take values in the sets 𝒳1, 𝒳2 and 𝒴, respectively. The joint mutual information between the pair of variables *X*1, *X*2 (joint variable constructed from *X*1 and *X*2) and the variable *Y* is expressed as:

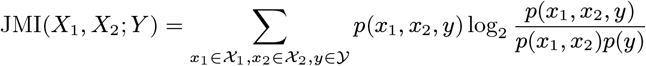

where *x*1, *x*2 and *y* are index variables that represent each possible value of the random variable *X*1, *X*2 and *Y*, respectively, *p*(*x*1, *x*2, *y*) is the joint probability distribution of *X*1, *X*2 and *Y, p*(*x*1, *x*2) is the joint probability distribution of *X*1 and *X*2, and *p*(*y*) is the marginal probability of *Y*. ∑ denotes the sum over the possible values of the three random variables.

The normalized joint mutual information (njMI) is calculated as follows:

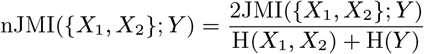

where H(*X*1, *X*2) is the joint entropy of the pair *X*1, *X*2, and H(*Y*) is the entropy of *Y*.

## Supporting information

Supplementary Material This section provides information necessary for a complete understanding and replication of the study.

## Acknowledgments

We would like to thank the members of Tolias Lab at the Department of Neuroscience at Baylor College of Medicine in Houston, TX for performing the two-photon (mesoscope) experiments, sharing the datasets with us, and providing valuable feedback about the data. We are also grateful to Vassilis Sideridis and Nikolaos Ntorvas who contributed significantly to the implementation of several aspects of the preliminary analysis. Several student interns as part of their training, their B.Sc. Thesis, or research assistanship funded by the Institute of Computer Science at FORTH, were involved in this project by implementing or validating different parts of the preliminary analysis, namely, Marianna Papadostefanaki, Andreas Sapoutzis, Ioannis Siriopoulos, Ioannis Fotis, Orestis Mousouros, Maria Plakia, Ioannis Charalambakis, Andreas Karambasis, Christos Mamoudis, Kostantinos Kokolakis, Charalampos Peteinarelis, Theodora Sambalou, Nikolaos Tzanakis, and Emmanouel Patronakis. Finally, we would like to thank Dr. Georgios Keliris (FORTH) for reviewing the paper and providing comments.

This work has received funding from the European Union’s Horizon 2020 research and innovation program under the Marie Sk-lodowska-Curie grant agreement No 101007926 and from the Hellenic Foundation Research Institute (HFRI), the neuron-AD project number 04058 and neuronXnet project number 2285 (PI: Maria Papadopouli). This research was also supported by R01 NS113890, and R21 NS127299 (PI: Stelios Smirnakis). AWS resources were provided by the National Infrastructures for Research and Technology GRNET and funded by the EU Recovery and Resiliency Facility.

## Code Availability

The code and additional information about the algorithms will be released during the peer review process. We will also release the code/algorithm as soon as the manuscript is published in a neuroscience journal (in github).

## Notes

### Competing Interest Statement

The authors have declared no competing interest.

### Summary of Updates

Most of the revisions here are editorial. We have added new plots on Figure 7 in the manuscript and new related plots in the Supplementary. Also, we have cited and very briefly discussed our recent work in applying this methodology in spiking deep learning models.

