## Supplementary Material This section provides information necessary for a complete understanding and replication of the study. for "From Functional Architecture to Organizing Principles of Neuronal Ensembles in Mouse Area V1"

This section provides information necessary for a complete understanding and replication of the study. Throughout the paper, the terms “spontaneous conditions,” “spontaneous neuronal activity,” and “resting-state” are used interchangeably.

**Assumptions and Default Values**: Below we state some of the assumptions and default values that apply in most of our analysis presented in the manuscript and supplementary material, *unless stated otherwise*: **1)** *Only pyramidal* neurons were recorded. Neurons with a *calcium event rate of less than 0.01 Hz* considering the entire period were excluded from this analysis. We also excluded the neurons that are located <= 15μm from the periphery of the field of view were also excluded in order to avoid potential edge effects arising from incomplete correction of motion artifacts (see Table 1). **2**) The deconvolved signal was thresholded (see methods) to yield calcium “eventograms" that were analyzed. The *threshold 1.5* was the default value for the analysis presented here. In the next section, we showed robustness of pairwise correlations under different deconvolution thresholds. **3)** We use the terms *functional “connection”* and *“edge”* (a term borrowed from graph theory) interchangeably. **4)** The default threshold for the *statistical significance* in the pairwise neuronal correlation (STTC weight) is at *z-score > 4*. **5)** The default period of the *estimation of the STTC weights is the one hour*, which is the longest duration recording available to us. 6) The pairwise correlations are formed using the synchronous STTC correlations, i.e., an *STTC Δt window (or lag) of 0 (*synchronization within one imaging frame). However, in the analysis, we have evaluated the impact of the time period, STTC Δt window, z-score thresholds, and correlation metric and our key results persist. **7)** The 1FC groups are formed based on the *pairwise correlations* estimated considering the *entire one hour interval of spontaneous conditions.* **8)** The **default datasets** used here are collected **in resting-state (i.e., spontaneous conditions),** which is the main focus of this work (and the first part of this Section)**.** We also present the functional connectivity under the *stimulation presentation condition (“Monet” stimulus, one hour duration, see methods), to* i) examine whether the principles identified under resting-state persist also under stimulus presentation; ii) assess information "transmission".from L4 to L2/3 under stimulus conditions. We explicitly indicate in the caption of Figures or legends of the plots if the datasets used are obtained via *the stimulus presentation experiments.* **9)** Neurons in L2/3 are analyzed at each layer, namely, L2 and L3, *separately*. For example, when we focus on the intra-layer L2/3 functional connections, we identify the (intra-layer) functional connections between pairs of L2 neurons, then separately between L3 neurons, and we then consider the union of these functional connections. Due to the **similarities of the two layers,** in the plots, we present the **L2/3 results of their union** (aggregate appropriately across mice). Insets in the histograms indicate the *sample mean ± standard deviation* across mice (n=5). The error bars correspond to the *SEM per specific bin across mice* (n=5). P-values: “*” < 0.05; “**” < 0.01; and “***”: < 0.001. The highest p-value obtained from the permutation of means, the Welch's t-test, and the ANOVA F-test is considered for the level-of-significance.

| | **M1** | **M2** | **M3** | **M4** | **M5** | **Across Mice** |
|---|---|---|---|---|---|---|
| **L2** | 1,402 | 1,733 | 3,245 | 1,334 | 1,357 | 1,814 ± 816 |
| **L3** | 1,503 | 1,706 | 1,113 | 1,280 | 1,344 | 1,389 ± 226 |
| **L4** | 1,271 | 888 | 659 | 1,179 | 923 | 984 ± 244 |
| **Total** | 4,176 | 4,327 | 5,017 | 3,793 | 3,624 | 4,187 ± 543 |

**Table 1: Number of V1 neurons per layer in our dataset.** The first three rows represent neurons in layers 2, 3, and 4, respectively. The last row provides the total count of V1 neurons. Each column corresponds to a different mouse, with the final column indicating the mean and standard deviation across mice.

| | M1 | M2 | M3 | M4 | M5 | Across Mice |
|---|---|---|---|---|---|---|
| **≤0.01Hz** | 1 (0.02%) | 27 (0.62%) | 20 (0.40%) | 12 (0.32%) | 0 (0.00%) | 12 ± 12 (0.3 ± 0.3%) |
| **At edge** | 313 (7.50%) | 349 (8.07%) | 300 (5.98%) | 334 (8.81%) | 258 (7.12%) | 311 ± 35 (7.5 ± 1.1%) |
| **Total** | 314 (7.52%) | 375 (8.67%) | 319 (6.36%) | 346 (9.12%) | 258 (7.12%) | 322 ± 43 (7.8 ± 1.1%) |

**Table 2: Number of V1 neurons (and corresponding percentage) meeting exclusion criteria.** The first row lists neurons with an event rate ≤ 0.01Hz during the entire resting-state period. The second row indicates neurons within 15μm of the edge of the field of view (FOV). The third row represents neurons meeting at least one of these criteria. These neurons were excluded from analysis. Each column corresponds to a different mouse, with the final column indicating the mean and standard deviation across mice.

#### 1. Robustness of pairwise correlations under different deconvolution thresholds

The deconvolved signal was thresholded (see methods) to yield calcium "eventograms" that were analyzed. The results reported in the manuscript were robust to the choice of threshold (see suppl. Fig. 1.1). This threshold is meant to give the scale of the noise, and multiples of this are used as thresholds for determining the calcium events (See methods). Through the paper, the plots we present employ the **threshold 1.5** (fixed across mice and layers).

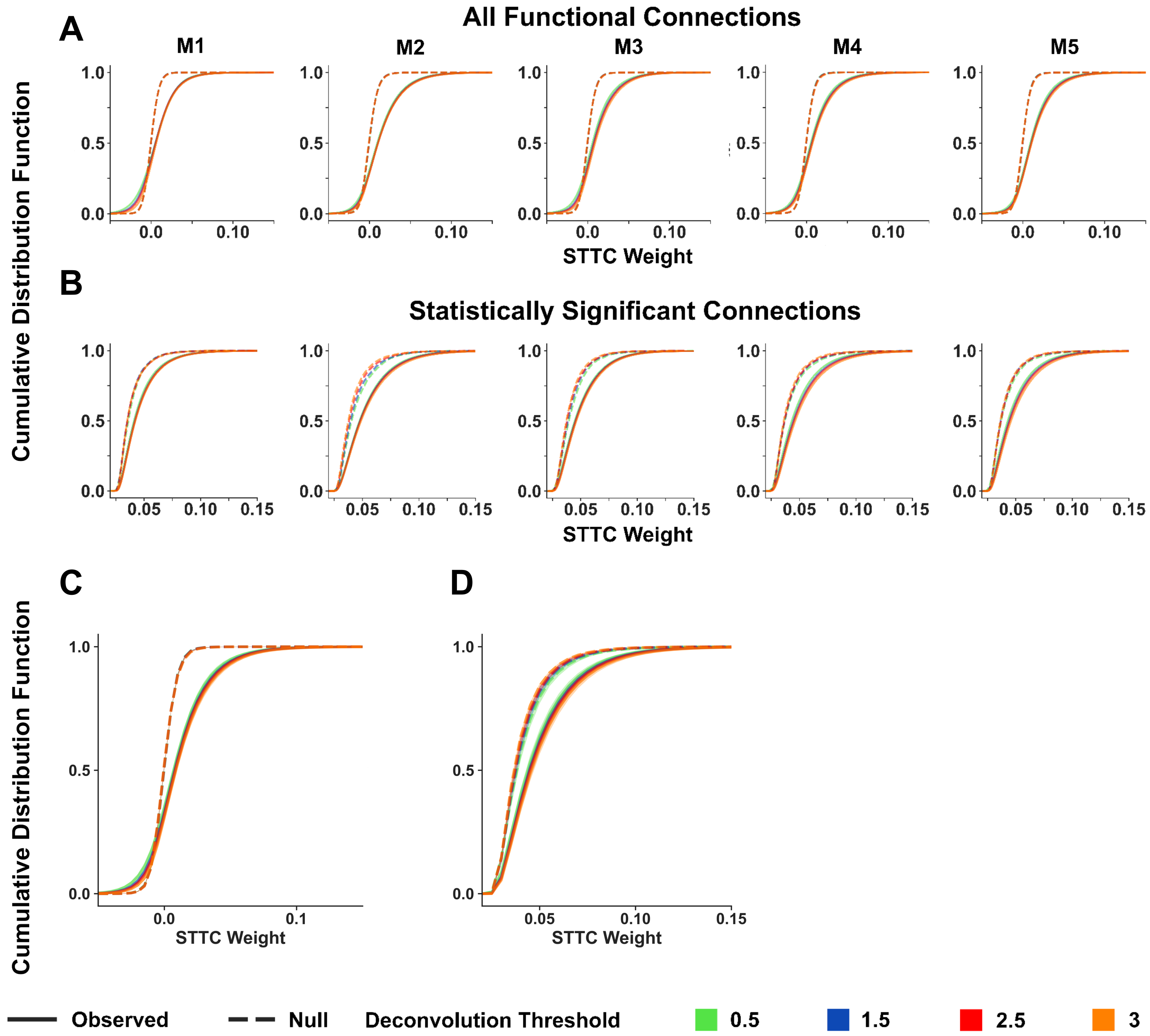


**Supplementary Figure 1: Pairwise functional correlations for different calcium event identification thresholds. A.** Cumulative Distribution Function (CDF) of STTC weights of all neuronal pairs recorded under resting-state conditions in V1 for each animal (M1 to M5). Null distributions (dashed lines) formed by random circular shifting of the neurons' calcium eventograms are also plotted on the same graph for comparison. Note that the shape of the observed STTC distributions is significantly different from the null and remains robust irrespective of the choice of deconvolution threshold for a broad range of thresholds (from 0.5 to 3). This threshold is meant to give the scale of the noise, and multiples of this are used as thresholds for determining the calcium events (See methods). **B.** Same as panel A but plotting only significant connections, where significance is defined at a z-score threshold of 4. **C.** Cumulative Distribution Function of STTC weights of all neuronal pairs recorded in V1 across all five mice. The lines and shaded regions indicate the mean and SEM across animals (n=5). **D.** Same as Fig. C but now only significant connections are plotted, where significance is defined at a z-score threshold of 4. Note that the fraction of pairwise functional correlations that are significant remains robust to the threshold of choice for calcium event identification. *All* pairwise connections *within and across layers* have been plotted here (A-D).

#### 2. Analysis of Pairwise Neuronal Correlations

This section examines the *temporal* and *spatial* dependencies on the pairwise neuronal correlations. We also computed STTCs by broadening the window of synchrony to 2 additional imaging frames (Suppl. Fig. 2.1), in which case we used an extension of the STTC measure we devised to take into consideration the temporal order of occurrence of spikes in A relative to B, i.e. reflect the probability that spikes of one neuron may systematically precede (or follow) spikes of the other (see Methods). Although as expected, the absolute number of significant functional connections (i.e., edges) changes depending on the choice of the window (Suppl. Fig. 2.1 B), the basic conclusions we report below remain the same whether synchronous or two imaging frames.

##### Temporal dependence of functional correlation measurements

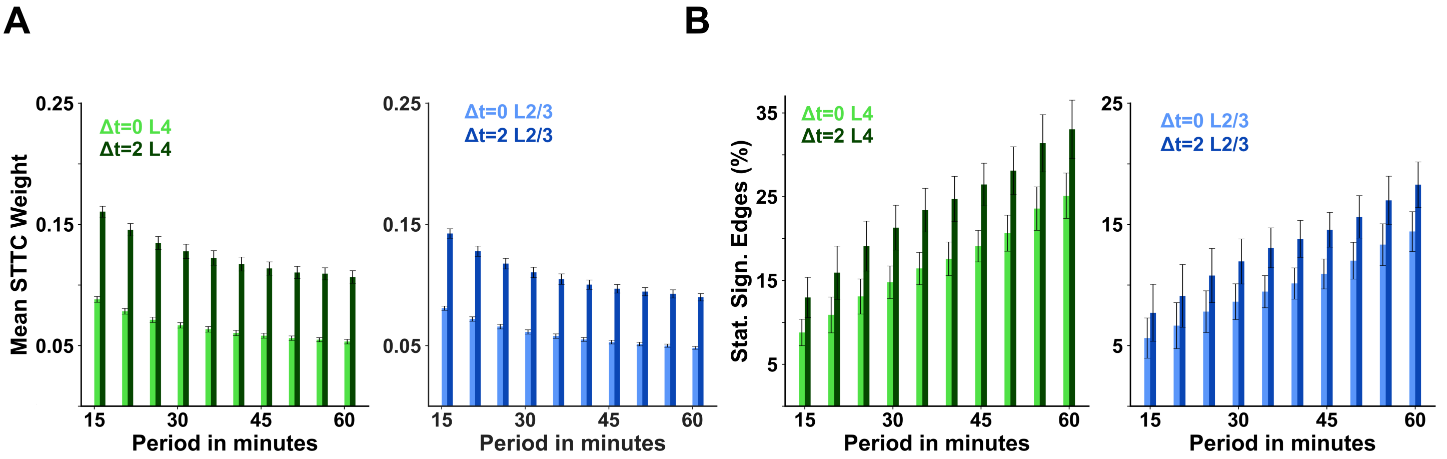


**Supplementary Figure 2. Effect of the time window (Δt) on the STTC measure of correlation. A.** The mean STTC weight of *all* statistically significant edges (z-score >4) for L4 and L2/3 (green and blue, respectively) as computed during different time intervals [0, T], T=15, …, 60 (with step size of 5), for *STTC Δt windows of 0* (synchronous frame, Δt=0, ~158.7ms) and *two additional calcium imaging frames (Δt=2, ~310ms), under resting-state conditions.* The darker color corresponds to STTC Δt=2, while the lighter to STTC Δt=0. The strength of the STTC weight of the edges increases for Δt ~310ms compared to Δt=0 (approximately doubles). **B**. Percentage of statistically significant edges (z-score >4) are plotted as a function of the duration of resting-state activity imaging for Δt=0 and Δt=2, denoted as PSSE(Δt, T), for *recording period T* and *STTC window Δt.* As expected, the percent of significant edges is higher when the correlation measure is computed over Δt ~310ms (green for L4, blue for L2/3). The mean *relative increase* in the percentage of statistically significant edges compared to the edges calculated by demanding synchrony, across time periods and considering all intra-layer L2/3 and L4 statistically significant edges ( $\sum_{\forall T} \frac{PSSE(2,T) - PSSE(0,T)}{PSSE(0,T)}$ ) is ~37%. Error bars represent the standard error of the mean across animals (n=5); Neurons with calcium event rate less than 0.01 Hz were excluded from this analysis.

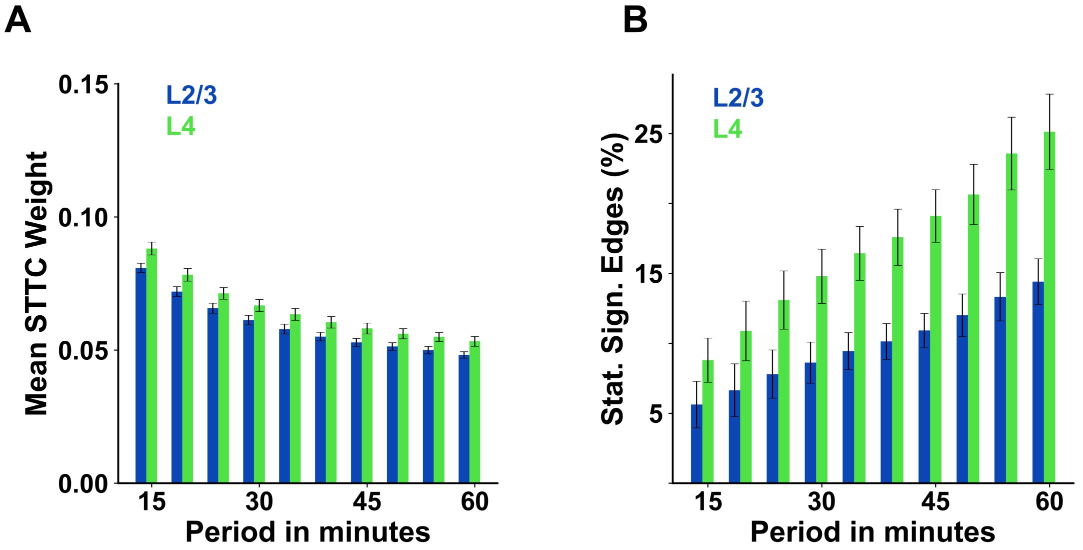


**Supplementary Figure 3. Impact of the duration of the recording on the STTC. A.** The mean STTC weight of **all statistically significant edges** (z-score >4) as computed during different **time intervals [0, T]**, T=15, ..., 60min (with step size of 5) of resting-state (spontaneous) neuronal activity imaging for Δt=0 (green bars for L4, blue bars for L2/3). As expected, the longer the recording, the smaller the mean STTC weight, as weaker correlations between neurons rise to significance. **B**. Percentage of statistically significant edges (z-score >4) are plotted as a function of the duration of spontaneous activity imaging for Δt=0. As expected the longer the duration, the larger the number of statistically significant edges. Error bars represent the standard error of the mean across animals (n=5); Neurons with calcium event rate less than 0.01 Hz were excluded from this analysis.

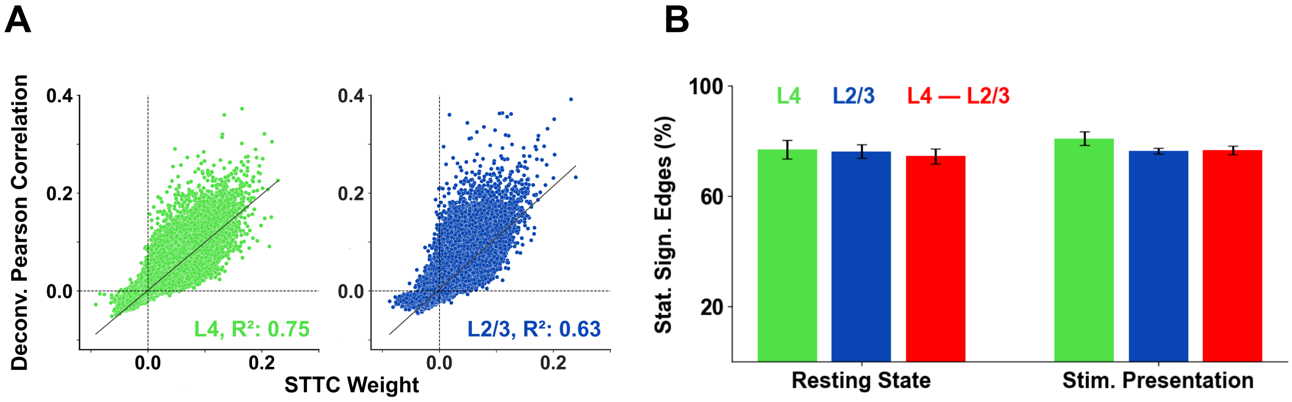


**Supplementary Figure 4 Functional connectivity between neurons formed using STTC on their *eventograms* vs. Pearson correlation on their *deconvolved* signals under spontaneous conditions and stimulus presentation. A.** Pairwise neural correlation using STTC on the calcium eventograms and Pearson correlation on the deconvolved signals, in all pairs, for an example mouse. Similar results were observed also in other mice. **B.** Percentage of the statistically significant edges based on STTC in L4 (green), L2/3 (blue), and from L4 to L2/3 (red), using z-score > 4, that *are also* statistically significant based on the deconvolved signal per layer case using z-score > 4, during resting state (left) and stimulus presentation (right). A significant number of neuronal pairs that are functionally connected using the STTC remain also functionally connected when the Pearson correlation is applied on their deconvolved signals. Error bars represent the standard error of the mean across animals (n=5).

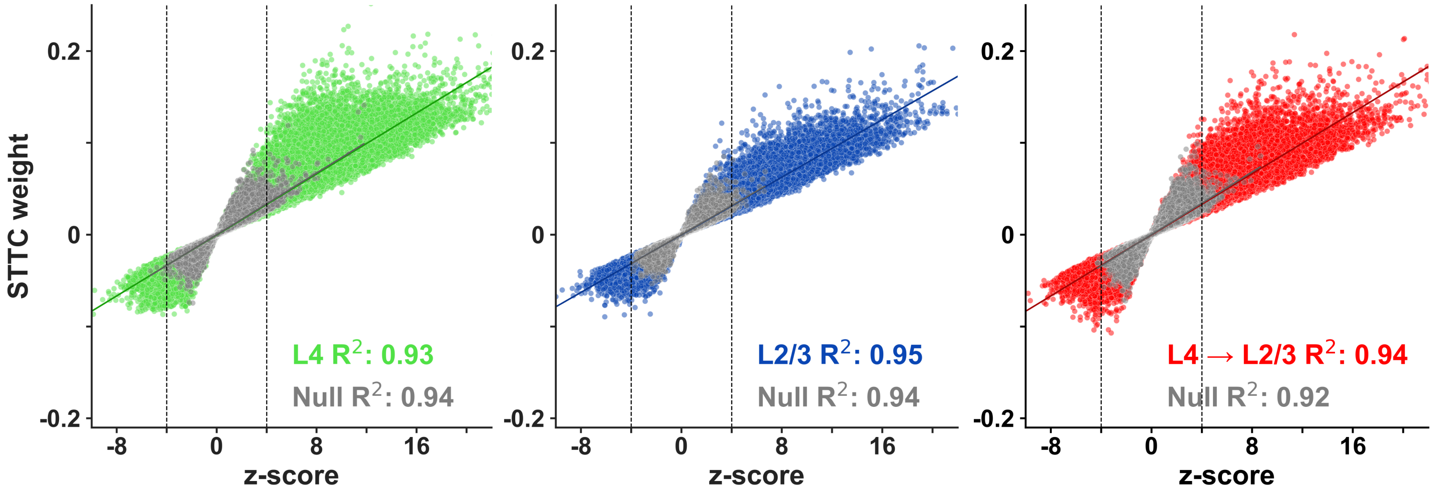


**Supplementary Figure 5. STTC weight of neuronal pairs across mice as a function of significance (z-score).** L4: green. L2/3: blue. L4 to L2/3:red. The higher the z-score, the larger the mean STTC weight and its variance. STTC connectivity weight values are approximately within the range of [0.02, 0.2] for z-score > 4. Gray: corresponding values derived from null distributions obtained by random circular shuffling. The STTC measurements presented here are obtained under spontaneous conditions (quiet wakefulness in the absence of visual stimulation) but essentially identical plots hold true under stimulation conditions.

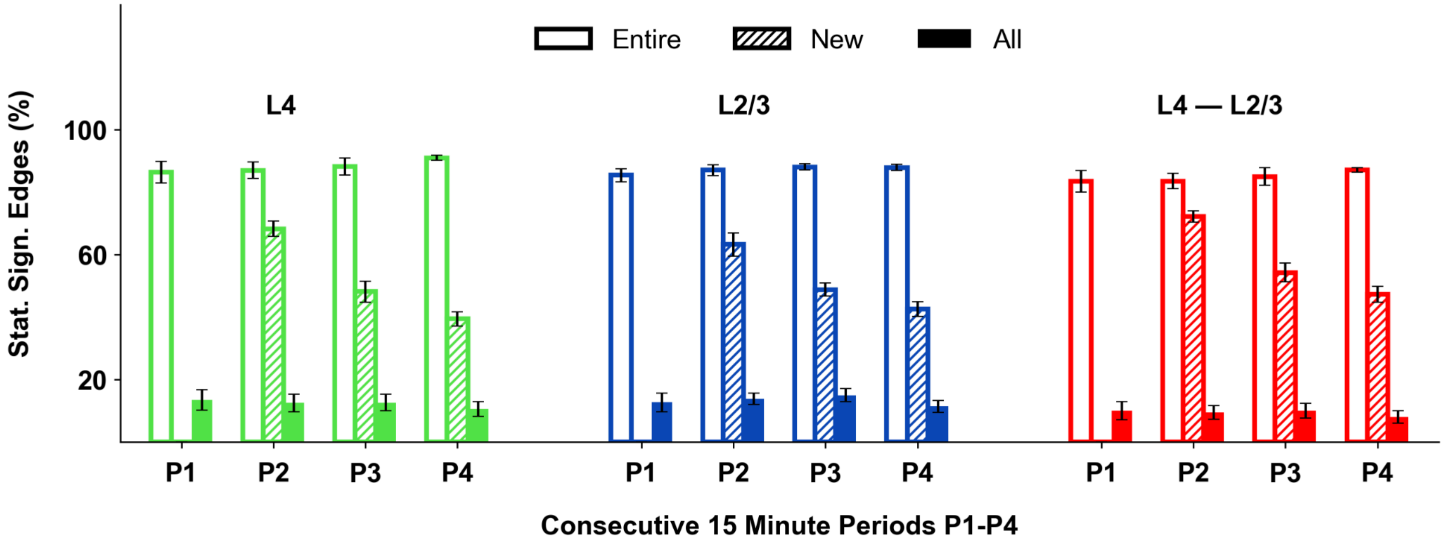


**Supplementary Figure 6. Robustness of functional connections and dynamic edges under spontaneous conditions.** Bar plots of the statistically significant edges (SSE), considering z-score >4, during *each 15-minute non-overlapping consecutive* period (P1-P4), which are *also* statistically significant in the *entire period* (Entire, white), the % of SSE in the specific period that has ***not*** been observed as statistically significant in *any other previous interval* (New, shaded), and the % of SSE in the specific epoch that are also statistically significant in all other *15-minute epochs* (All, solid). The first four leftmost bar plots correspond to L4. Of the statistically significant edges in the current period (indicated in the x-axis), approximately 12% of them remain statistically significant during *all other 15-minute epochs* (green, "All"), while over 80% of the edges in the current period are also statistically significant during *the entire one-hour recording period (green, "Entire")*. The other bar plots present the same statistics for L2/3 (blue) and for the inter-layer, from L4 to L2/3 (red). As expected, the percentage of new edges decreases in the following periods. Neurons with a calcium event rate of less than 0.01 Hz, considering the entire period, were excluded from this analysis; Error bars represent the standard error of the mean across animals (n=5).

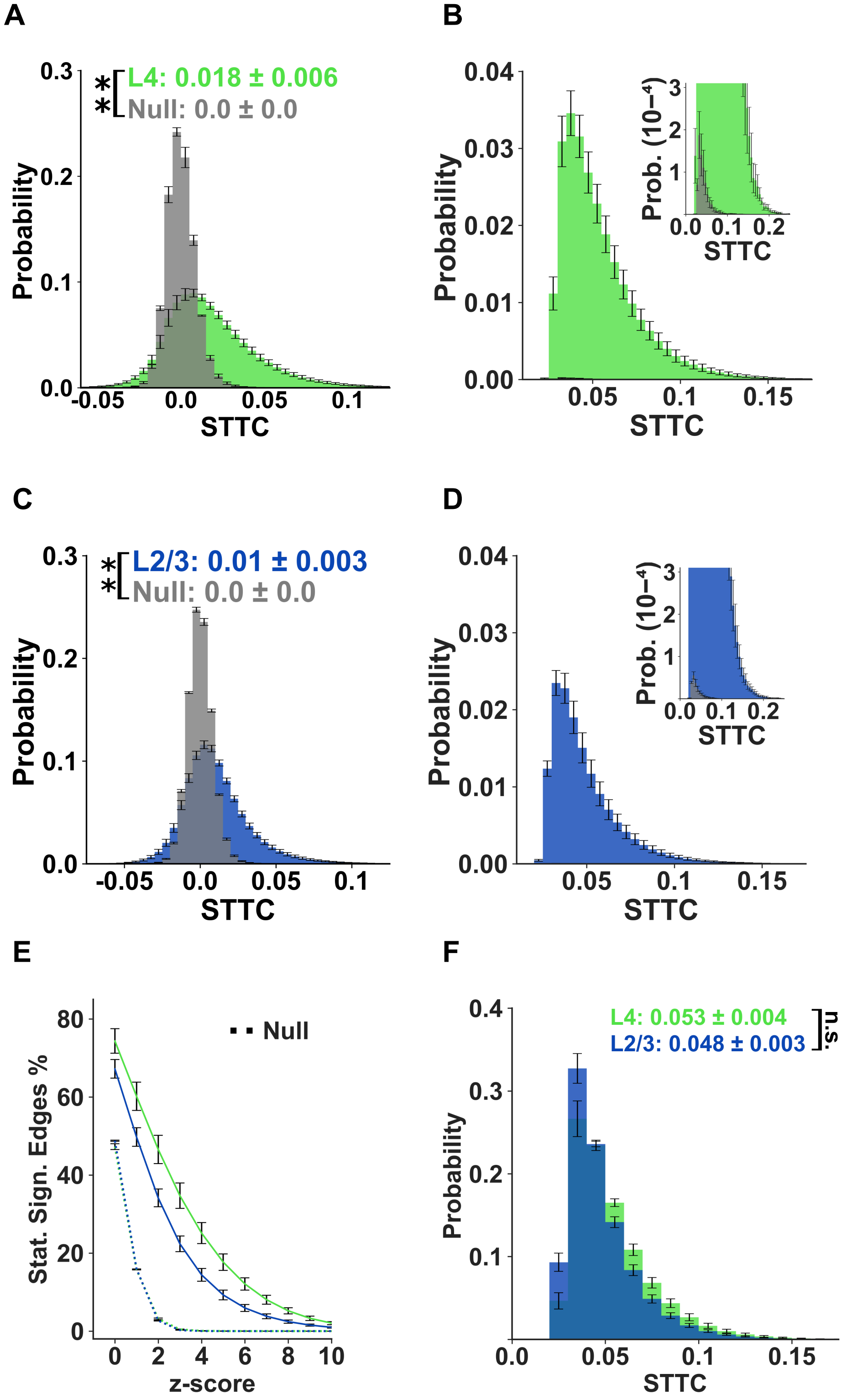


**Supplementary Figure 7. Intra-layer Pairwise Functional Connectivity under Spontaneous Activity, Quiet Wakefulness, Conditions. A,C.** Histograms of STTC values between pairs of neurons ("edges") in layer 4 (A; green) versus layer 2/3 (C; blue). *Gray:* corresponding null distributions obtained by random circular shifting. **B,D.** Histograms of STTC values between pairs of neurons with distributions (dotted lines). **F.** Histogram of STTC weights for observed statistically significant edges (z-score > 4). Pairwise STTC correlation strength is weak and distributed similarly across different layers. Values reported represent the mean ± standard deviation of the sample means across five mice (n=5), while error bars correspond to the standard error of the mean (SEM) across mice (n=5). P-values: “**” < 0.01; “n.s.”: non statistically significant. The highest p-value obtained from the permutation of means, the Welch's t-test, and the ANOVA F-test is considered for the level-of-significance.

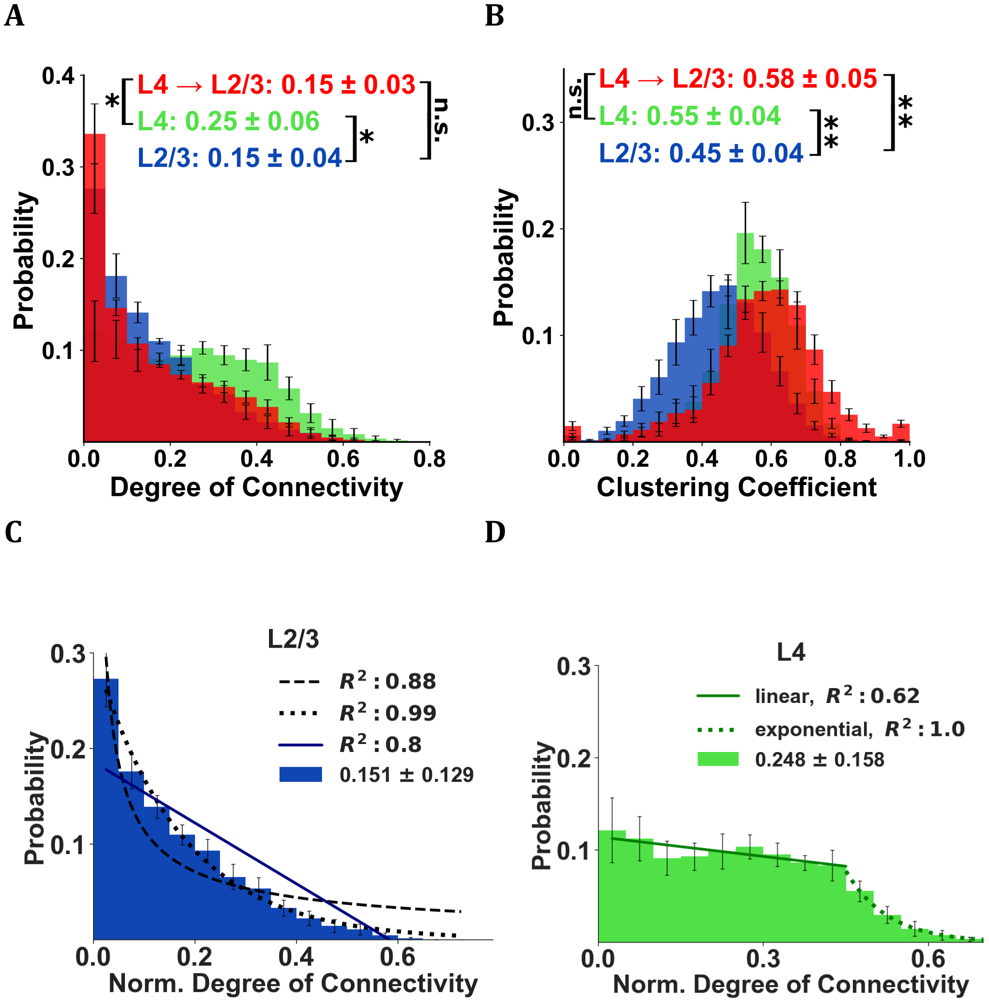


**Supplementary Figure 8. Degree of Connectivity and Clustering Coefficient Distributions**. **A.** Histograms of the degrees of connectivity (DoC) of neurons within L4 (green) within L2/3 (blue) as well as across L4-L2/3 (red) under resting state conditions, expressed as a fraction of the total number of neurons belonging to L4, L2/3 and L4 respectively. Only edges with statistically significant positive STTC (z-score > 4) were included in calculating the DoC. DoC distributions within L2/3 (blue) and across L4-L2/3 (red) fall rapidly as a function of the DoC, indicating a sparse functional network in which most neurons exhibited a small number of significant partners and high-degree nodes were relatively rare . In contrast, the DoC distribution within L4 (green) is nearly uniform up to considerably higher DoC values, indicating a more dense functional connectivity structure. **B.** The distributions of intra-layer clustering coefficients of one-degree functional connectivity groups within L4 (green), L2/3 (blue), and across L4-L2/3 (red). **C-D. Modeling the degree of functional connectivity in L4 and L2/3 across mice (n=5).** The fits of power law, exponential, and linear functions to the degree of connectivity *in resting-state* are denoted by dashed, dotted, and solid lines respectively. **C**. For the majority of mice, the degree of connectivity in Layer 2/3 follows an exponential distribution with $R^2$ above 0.97, although a power-law does provide a nice fit in the main part of the distribution up to its tail (with an $R^2$ of 0.88). We speculate that the ill-fit of the tail could be due to the fact that the data are "censored" (only a part of the neurons from L2/3 are recorded). **D.** The degree of connectivity in L4 follows a uniform distribution up to ~0.45 and after that, it falls exponentially (with an $R^2$ of 1). Error bars represent the standard error of the mean across mice (n=5). Insets: mean ± standard deviation of the sample means across mice (n=5). P-values: “*”< 0.05; “**”< 0.01; and “n.s.”: non statistically significant. The highest p-value obtained from the permutation of means, the Welch's t-test, and the ANOVA F-test, is considered for the level-of-significance.

##### Negative Functional Connectivity Analysis

We also computed statistically significant anticorrelated functional connections, including the STTC weights, and mapped the negative functional connectivity topography.

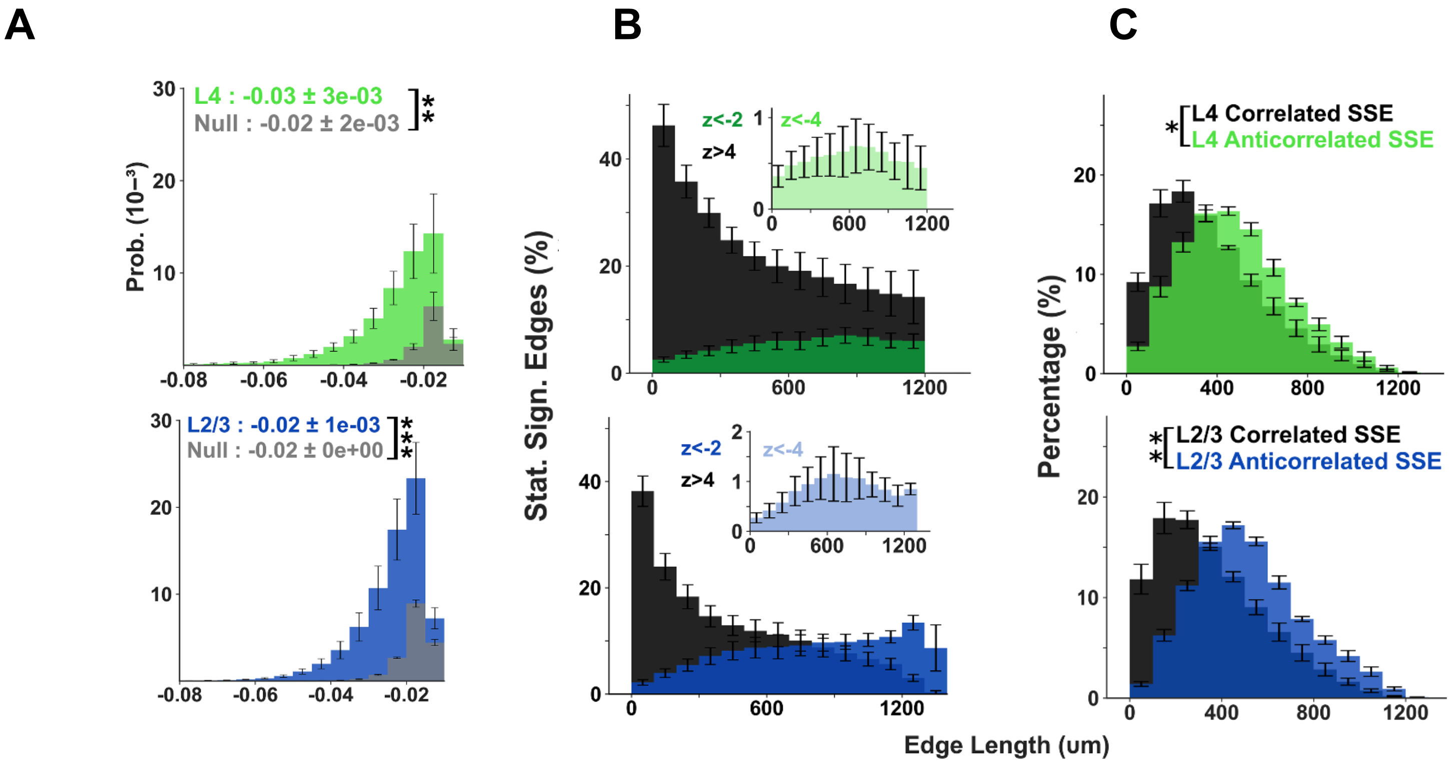


**Supplementary Figure 9. Negative functional connectivity as a function of distance during spontaneous activity. (A)** STTC weight distributions for negative edges significant at $z < -2$ for within-L4 pairs (green) and within-L2/3 pairs (blue). Gray denotes corresponding null distributions from circular time-shift controls; **(B)** Fraction of significant STTC edges per pyramidal neuron versus intersomatic distance (100-μm bins), shown for negative edges ($z < -4$, light; $z < -2$, dark) in comparison to positive edges ($z > 4$, black) in L4 (green) and L2/3 (blue). For each neuron, the fraction was computed as the number of significant edges in a distance bin divided by all possible edges in that bin; mean value per mouse was averaged across mice (n = 5; error bars, SEM). Negative edges are relatively rare at short range and increase with distance peaking at ~600–800 μm for $z < -4$ and >900 μm for $z < -2$. In contrast, positive functional correlation edges are concentrated at short distances. Note that the distance-dependent rise of negative-edge prevalence is slower in L4 than L2/3, while L2/3 exhibits a slightly larger percentage of negative stat. significant edges than L4. We also found that th**e** mean STTC weight of significant edges decrease only slightly with distance (not plotted here). **(C)** Distributions of significant positive ($z > 4$) and negative ($z < -2$) edges for L4 (top) and L2/3 (bottom) with distance, showing the longer-range bias for negative edges; $z < -4$ yields a distance distribution similar to $z < -2$ (not shown). Significance: permutation of means, Welch's *t* test, and ANOVA *F* test (largest *P* used); *$P < 0.05$, **$P < 0.01$.

##### Large Population Cofiring Events can be Achieved with Low Pairwise Correlations

Little pairwise correlation strength is needed to generate significant cofiring events among large aggregates of neurons [Shadlen & Newsome 1998]. A simple neuronal population model with mean firing rates commensurate to the ones we observed, shows that highly statistically significant cofiring events can be generated even when pairwise correlations remain low.

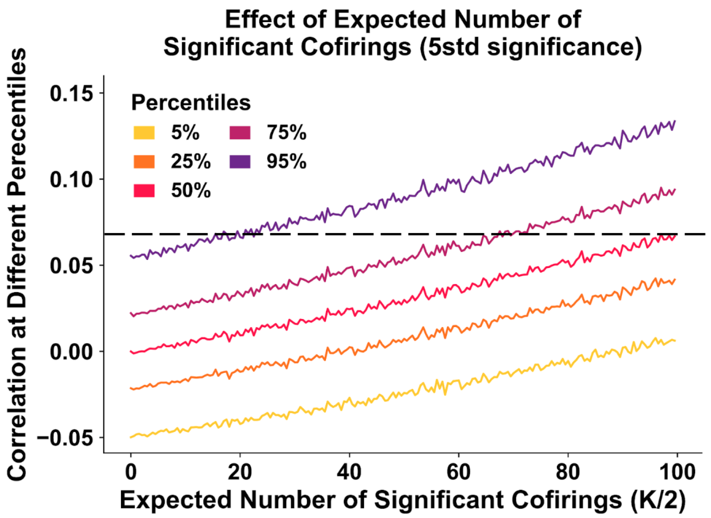


**Supplementary Figure 10. Large population co-firing bursts can occur despite weak pairwise correlations.** Toy Bernoulli model (N = 100 neurons; 1,000 frames) with matched mean firing rate (p = 0.1 per neuron per frame). In the ***independent*** condition (K = 0), neurons fire independently. In the ***burst*** condition, firing probability was increased on K randomly selected frames (p′) and reduced on all other frames to preserve the overall mean rate. p′ was chosen so that the expected population count on burst frames, Np′, was *5 SD above* the independent expectation: $Np' = (Np + 5\sqrt{(Np(1-p))})$. Because the population count on burst frames is approximately Gaussian around Np′, ~50% of burst frames exceed the 5-SD threshold; thus K/2 approximates the expected number of frames for which a significant number of co-firings occurs. Next, we compute all the pairwise correlations that emerge in the burst condition for each K, and plot the correlation values that correspond to various percentiles (5% to 95%) of the pairwise correlation distribution as a function of K/2. Notice that K=0 corresponds to the independent condition (first case), hence the median correlation (50% percentile) is ~zero for K/2=0. Even with ~100 significant co-firing frames per 1,000 frames (K/2 = 100), the median pairwise correlation remained low (~0.068, dashed black line), illustrating that prominent population co-firing can coexist with weak pairwise correlations.

#### 3. Relation of Tuning Properties to Spontaneous Connectivity

A natural question for us was to ask the degree to which neurons with similar tuning functions exhibit increased functional connectivity in resting-state (spontaneous conditions). The following Figures illustrate the relation of spontaneous function (positive and negative) connectivity to tuning properties.

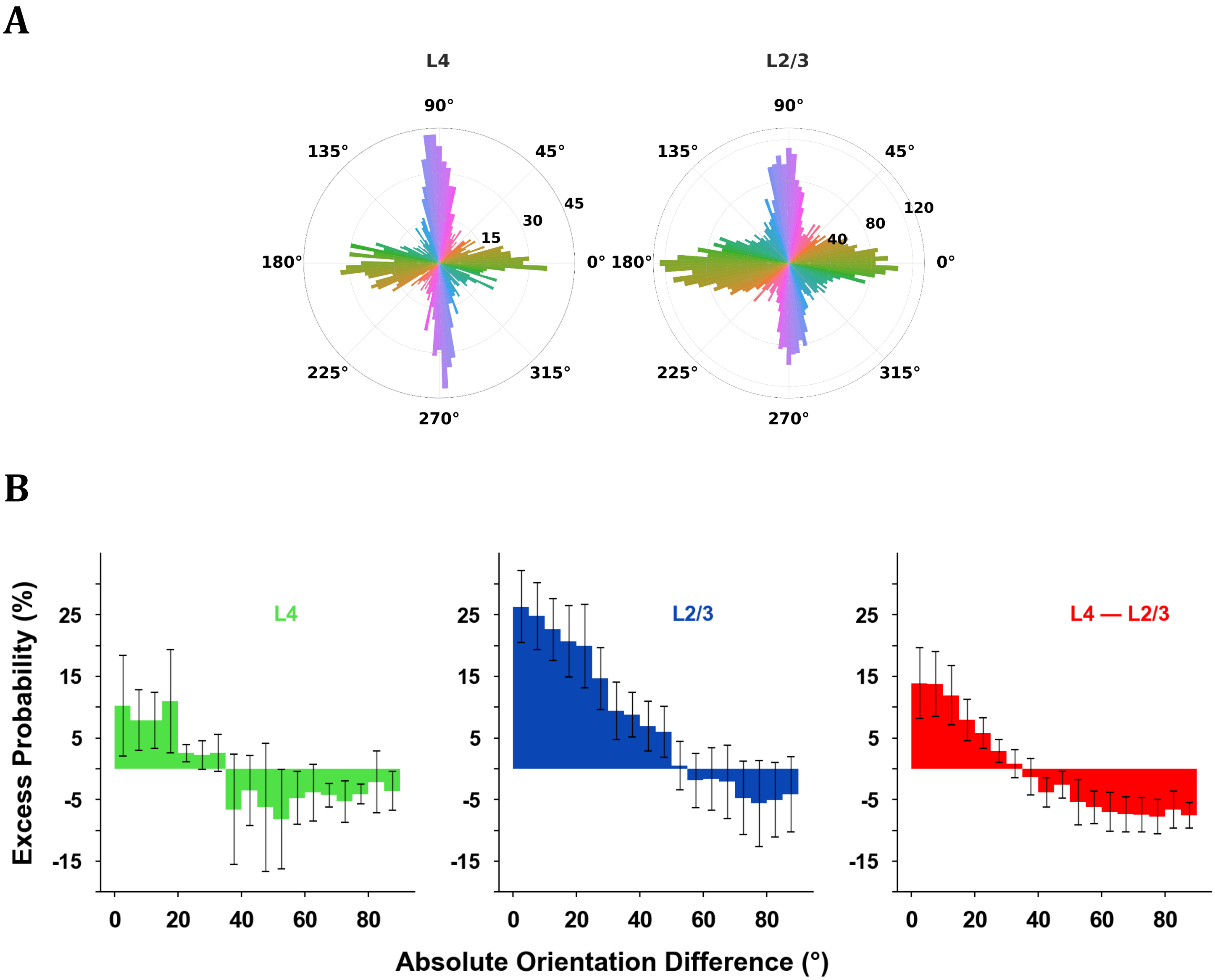


**Supplementary Fig. 11 | Spontaneous functional connectivity shows only a modest bias for tuning similarity.** Tuning function similarity implies a bias towards stronger functional connectivity but leaves enough room for neurons with disparate functional properties to also be strongly functionally connected. **(A)** Population direction/orientation tuning distribution pooled across mice (methods as in Fahey P. *et al*[1]). Polar bins are colored by preferred orientation; radial extent denotes neuron count. The known cardinal bias in mouse V1 is apparent; preferred-orientation angles were corrected for minor monitor/eye alignment offsets. **(B)** Enrichment of tuning similarity among significantly correlated pairs. Histogram shows, for each layer pairing (L2/3–L2/3, blue; L4–L4, green; L4–L2/3, red), the difference between the orientation-difference distribution of significantly correlated pairs (STTC z-score >4) and a null distribution defined by nonsignificant pairs (z-score [-2,2]), normalized by the

[1] Fahey, P. G., Muhammad, T., Smith, C., Froudarakis, E., Cobos, E., Fu, J., Walker, E. Y., Yatsenko, D., Sinz, F. H., Reimer, J., et al. A global map of orientation tuning in the mouse visual cortex. BioRXiv (2019).

null-bin probability. Orientation difference was computed as the minimal absolute difference between preferred orientations on [0,180), yielding a range of [0, 90] degrees (Methods). Error bars, SEM across mice (n=5).

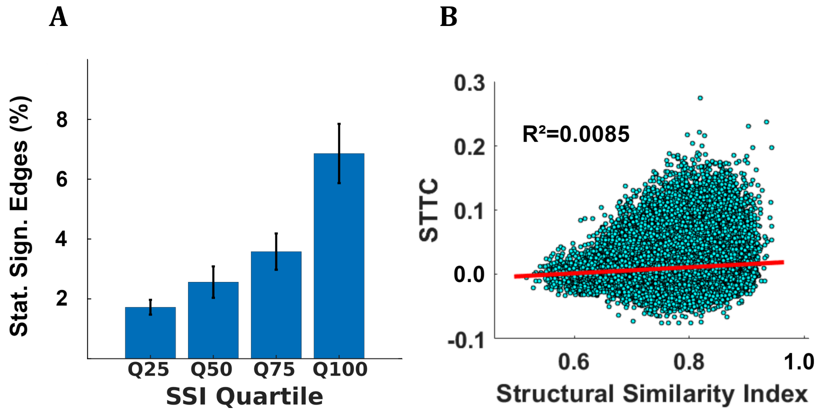


**Figure 12 Structural similarity between receptive fields and pairwise neuronal correlations. A.** To explore the trend in more detail, we grouped the cell pairs into quartiles according to the values of the **structural similarity (SSI)** between their receptive fields and calculated the percentage of pairs in each quartile having an STTC z-score ≥4. Bars represent the mean percentage of significant STTC pairs in each quartile, while error bars correspond to the SEM across mice (n=5). A clear difference exists between lowest and highest quartiles in terms of the percentage of functionally connected pairs (Y-axis: proportion between functionally connected (Z≥4) and non-connected pairs (Z<4)) (p = 0.0079, Wilcoxon ranksum test across mice; lowest quartile: 0.017±0.0025, highest quartile: 0.069±0.01). **B.** Receptive field similarity versus strength of functional connectivity: high similarity in receptive field maps does not imply stronger functional connectivity between cells on average across population. Relationship between STTC correlation coefficient and structural similarity of receptive field maps in visually responsive cells of an example mouse. Although there is a trend of cells with strongly similar receptive field maps also having a higher probability of stronger STTC connection, this trend is weak and non-linear ($R^2$=0.0085, red line shows linear fit).

#### 4. Characterization of the Functional Connectivity Structure

The following paragraphs identify the “persistent” functional connections that form the “core” network and its structure and spatial spread. Then, we assess the small worldness and network robustness of the functional connectivity architecture.

##### 4.2 Core Functional Network

The core functional network at a certain layer consists of the “persistent” edges, i.e. edges that are statistically significant (z-score>4) during ***all*** *four 15-minute non-overlapping consecutive* periods of the recording under resting state. Note that all persistent edges are statistically significant during the full hour. We found that they exhibit higher STTC weights and z-scores during the full hour, more prominent similarity in their tuning function, and shorter length than the other statistically significant edges. Similar results are observed when we define the core functional network based on *non-consecutive* frames.

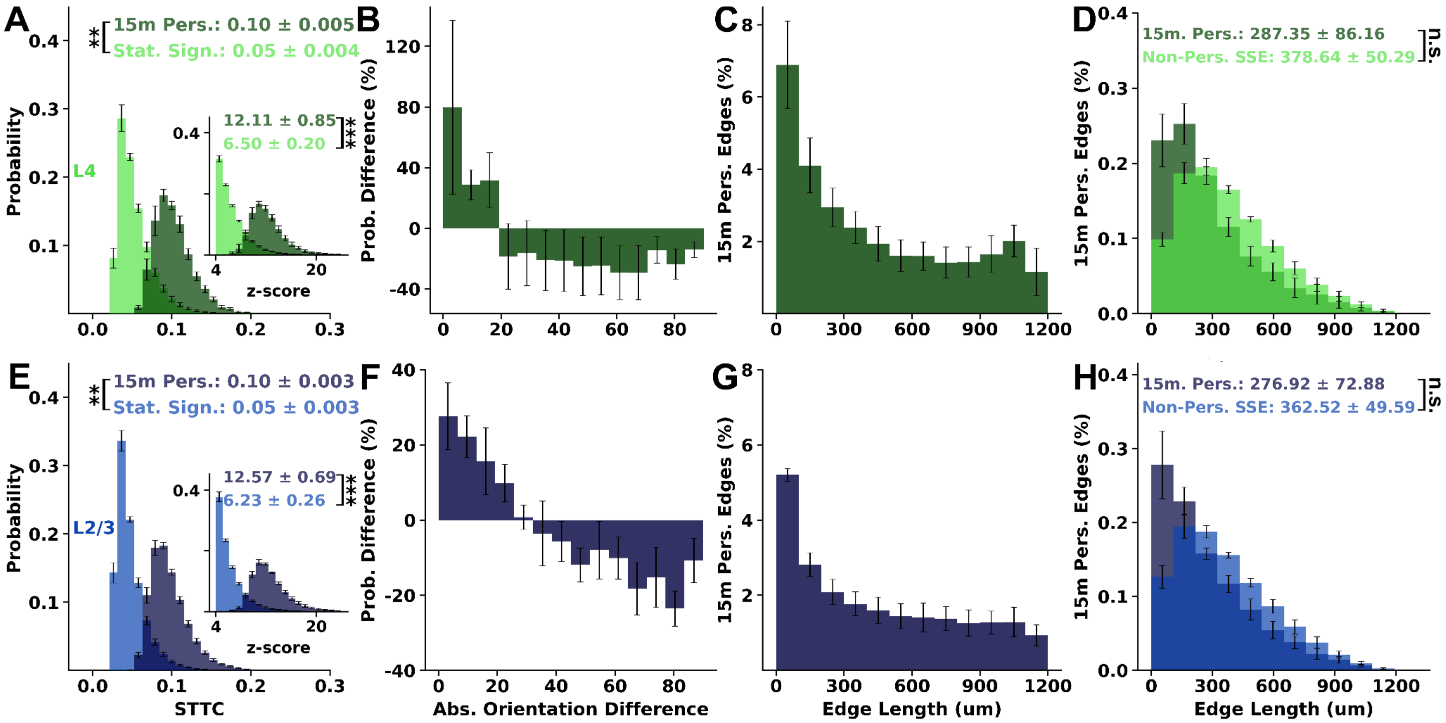


**Supplementary Fig. 13| Persistent functional edges are stronger, shorter-range, and more similarly tuned.** *Persistent* edges were defined as connections significant (**z** > 4) in **all four** non-overlapping 15-min spontaneous epochs; comparisons are to all significant edges identified from the full 1-h recording (STTC, **z** > 4). **(A–D) Intra-L4** edges: **(A)** distributions of STTC weights (and z-scores, inset) for persistent versus all significant edges; **(B)** enrichment of persistent edges as a function of the absolute preferred-orientation difference (Δθ, 0–90°) between neuronal pairs. For each absolute orientation difference bin, we computed the percentage of persistent edges in that bin (i.e., number of persistent connections with absolute orientation difference in that bin over the entire set of persistent L4 connections); Similarly, we computed the percentage of non-persistent statistically significant edges in that bin. We then report their difference over the percentage of statistically significant non-persistent edges in that bin. **(C)** fraction of significant edges that are persistent versus inter-somatic distance (100-μm bins); **(D)** edge-length distributions. **(E–H)** Same analyses for **intra-L2/3** edges. Values are mean ± SD of mouse means; error bars, SEM across mice (n = 5). Significance: P-values "**" < 0.01; "***" < 0.001 and "n.s.": non statistically significant; we report conservative *P* values (maximum across permutation-of-means, Welch's *t*, and ANOVA *F* tests; see Methods). For Figures B and F, only the OT neurons have been considered.

#### 4.3 Small-worldness

For each layer, namely L2/3 and L4, the observed graphs based on the pairwise statistically significant connections (with z-score >4) were formed and characterized in terms of clustering coefficient and average path length. We then compared them with their corresponding theoretical graph models, namely **1)** the **Erdős–Rényi** ER($N$,$p$), where $N$ is the total number of neurons in the layer of interest and $p$ the probability that two neurons are functionally connected, which is equal to the mean normalized degree of connectivity of the respective biological network, and **2)** a **regular ring** with $N$ nodes and fixed degree of connectivity for each node, equal to the nearest even integer of the mean degree of connectivity of the observed network. Specifically, in the regular ring, each node is placed in a circular topology, connected to its $k$ nearest neighbors on its left and to its $k$ nearest neighbors on its right. Each theoretical graph was constructed using the same number of neurons as their corresponding observed functional network. Note the small average path length of the observed functional connectivity, which is closer to the one estimated on the Erdős–Rényi graphs than that of the regular ring graphs. On the other hand, the clustering coefficient of the observed network is closer to that of the regular ring. Thus, the functional connectivity of L4 and L2/3 manifest small-worldness. L4 exhibits small-world qualities more prominently than L2/3, having significantly higher mean clustering coefficient (p-value 0.026) and significantly lower mean path length than L2/3 (p-value

0.049). Looking at L2 and L3 independently, we found that  the small-worldness index of L4 is significantly higher than that of L2, while no statistically significant difference was observed between L4 and L3.

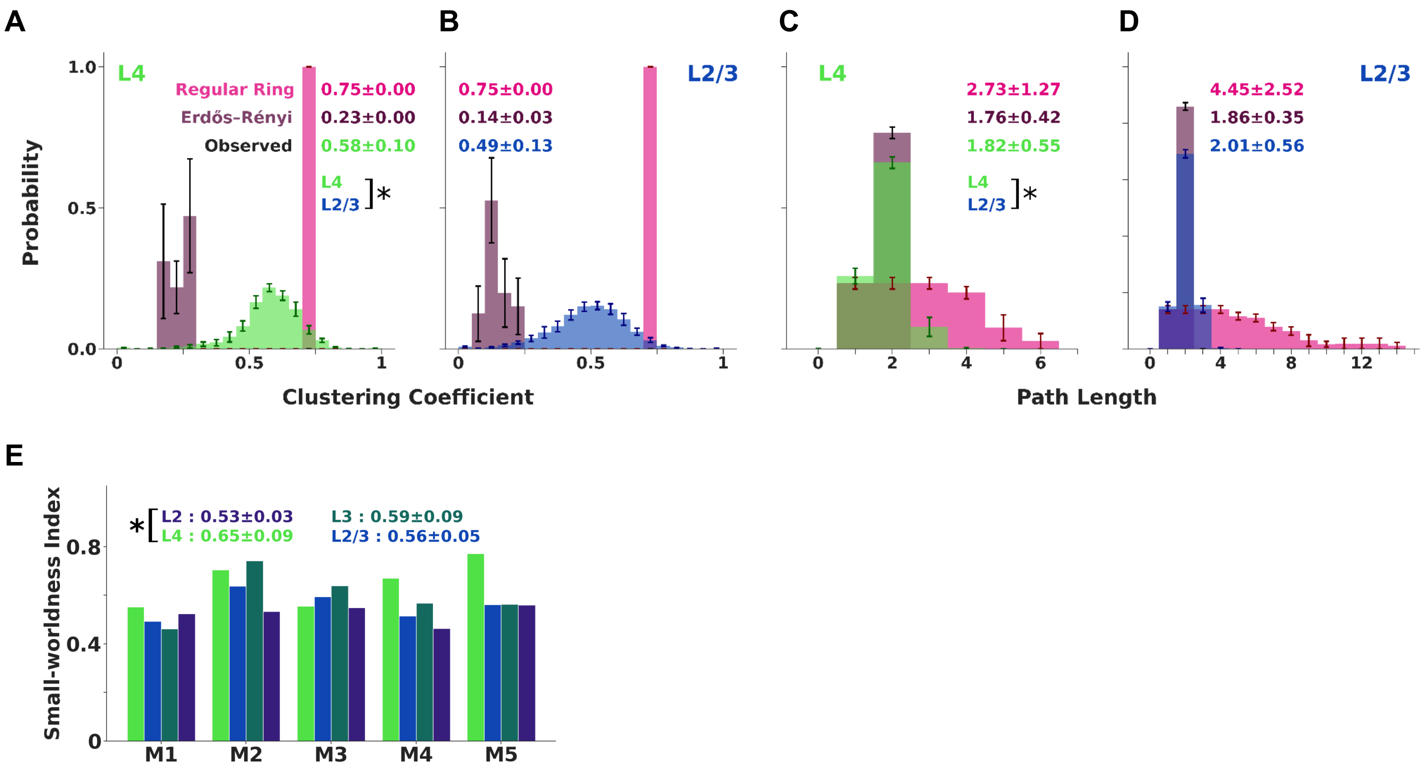


**Supplementary Figure 14 Functional connectivity in L4 and L2/3 exhibits small-world characteristics.** The layer network graphs based on the pairwise statistically significant connections (with z-score >4), estimated at resting-state, were formed and compared with their corresponding theoretical graph models, namely the **Erdős–Rényi** and the **regular ring**. **A)** Histogram of the clustering coefficient of L4 neurons (green) vs. the corresponding theoretical graphs random Erdős–Rényi (purple) and regular ring (pink), across mice (n=5). Here the clustering coefficient is estimated considering the entire set of frames without excluding the frames that the index neuron fires, but this does not alter the basic conclusion. **B)** Same as A, for Layer 2/3 (blue). **C-D)** As in A-B but now we histogram the shortest path length between pairs of neurons. Notice the small average path length of the observed functional connectivity in both layers, closer to the one estimated on the Erdős–Rényi graphs compared to regular ring graphs. On the other hand, the clustering coefficient distributions of the observed networks are closer to that of the regular ring graphs. This behavior is a signature of small worldness, manifest in both L4 and L2/3. Specifically, L4 exhibits small-world qualities more prominently than L2/3, having significantly higher mean clustering coefficient and significantly lower mean path length than L2/3. **E)** The mean small-worldness index across mice computed separately for each layer (green: L4, purple: L2, dark turquoise: L3, blue: L2/3)**.** L4 has a small-worldness index significantly higher than L2, while there is no difference between L4 and L3. Error bars correspond to SEM across mice (n=5). Insets report mean ± standard deviation across mice; P-values: "*" < 0.05. The highest p-value obtained from the permutation of means, the Welch's t-test, and the ANOVA F-test is considered for the level-of-significance**.**

#### Robustness

A simple graph-theoretical metric of *network robustness* (e.g., resilience to different types of node or edge failures) is based on the presence of a giant subnetwork component, with the property that any two of its nodes are connected via some path. A giant subnetwork component by definition contains a large number of nodes of the original network and their edges. The graph based on the L4 functional connectivity contains a giant component that persists under high z-score thresholds used to determine the statistically significant connections (up to Z-score 8) and includes approximately 80% of the L4 neuronal population (Suppl. Fig. 4.8A). L2/3 follows a similar trend but up to a somewhat lower z-score value.

Similar results are obtained when we employ the Molloy-Reed criterion of robustness. This criterion is derived for a particular family of random graphs with a specific distribution of normalized degrees of connectivity, to examine the robustness of the functional networks. **The Molloy-Reed criterion quantifies when a network is expected to lose its giant component.** The Molloy–Reed criterion is derived from the basic principle that in order for a giant component to exist, on average, each node in the network must have at least two links. Based on that criterion, one can obtain a critical threshold for the fraction of nodes needed to be removed for the breakdown of the giant component of the network, providing an indication of its robustness. According to the "Molloy-Reed"-based network robustness index, calculated as the ratio of the second moment of the degree of connectivity about zero over the mean degree of connectivity, the functional-connectivity of layers 2, 3 and 4 exhibits significant robustness (well above 2), much larger than the corresponding Erdős–Rényi and regular ring graphs. Even for very high z-score thresholds, the resulting functional networks at all layers remain well-connected.

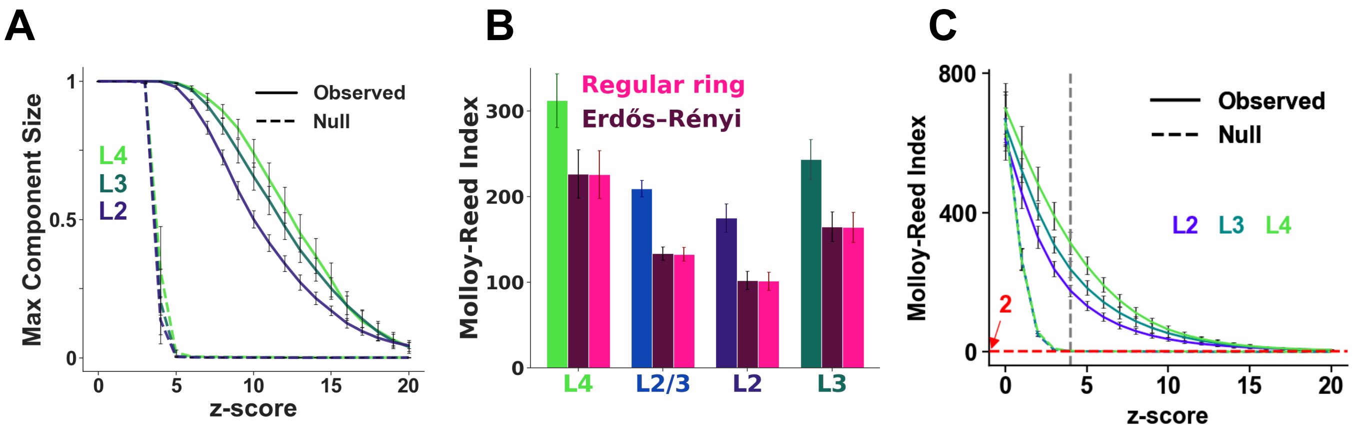


**Supplementary Figure 15| V1 functional networks in L4, L3, and L2 are highly robust.** Networks were constructed from resting-state activity (results were qualitatively similar during stimulus presentation). **(A)** Fraction of neurons contained in the largest connected component ("giant component") as a function of the z-score threshold used to define significant functional links: at $z \geq 4$, L4/L3/L2 networks contain a giant component spanning nearly all neurons and it persists at stricter thresholds. Null networks (constructed by independently circularly shifting each neuron's event/spike train by a random offset, recomputing pairwise statistics, and thresholding identically) lose their giant component rapidly. **(B)** Robustness quantified by the Molloy–Reed index, $\langle k^2 \rangle / \langle k \rangle$ (robust if >2), where k is the degree of connectivity, at $z \geq 4$; to control for layer-dependent neuron counts, neurons were subsampled within each mouse to match the smallest layer and indices were computed on the induced subgraphs. L4 is most robust, followed by L3 and then L2 (L2/3 intermediate); regular ring and Erdős–Rényi graphs show substantially lower robustness. **(C)** Molloy–Reed index versus z-score threshold for observed (solid) and null (dashed) networks; the red line marks index = 2 and the gray line marks $z = 4$.

#### 5. Predicting L2/3 pyramidal neuron activity from the cofiring of their L4 1FC group

This section focuses on the **putative communication modules across layers,** starting with the characterization of the main inter-layer neuronal ensemble unit of an L2/3 neuron in terms of degrees of connectivity and spatial spread of its L4-1FC connections. It then examines the relation between the independence in the firing of pairs of L2/3 putative recipient neurons and the sizes of their corresponding L4-1FC groups as well as the overlapping of their L4-1FC groups. In studying the response properties of L2/3-neurons with respect to their L4-1FC group size, we searched for a **normalization** by a power of the group size, to have similar response curves irrespective of the group size, based on a simple theoretical model (described below). Finally, we demonstrated that the accuracy of predicting L2/3 responses does not improve when, in addition to the aggregate number of cofiring events, we include information about the identity of L4-1FC group neurons that fire.

##### Resting State (i.e., Spontaneous Conditions)

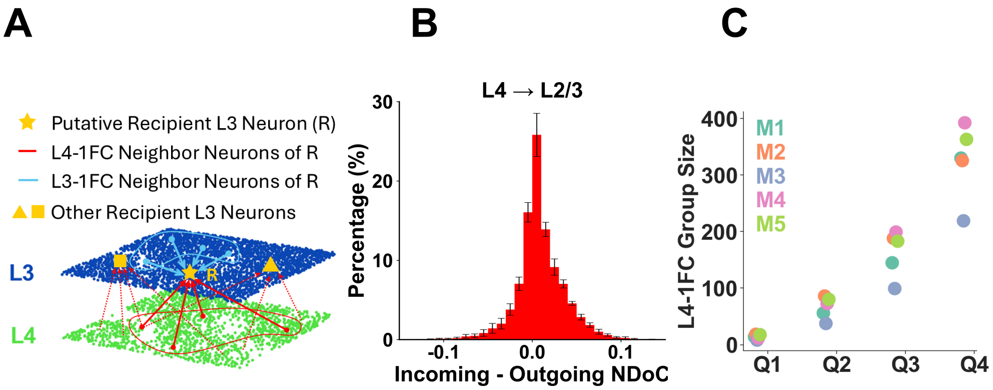


**D**

Number of Neurons in Layers

| | Layer 4 | Layer 3 | Layer 2 |
|---|---|---|---|
| M1 | 1192 | 1392 | 1278 |
| M2 | 803 | 1572 | 1577 |
| M3 | 612 | 1047 | 3039 |
| M4 | 1079 | 1171 | 1197 |
| M5 | 865 | 1241 | 1260 |

**Supplementary Fig. 16 | Putative cross-layer communication modules defined by functional connectivity. (A)** Schematic of an example L2/3 reference neuron (R; shown in L3, yellow star) and its **L4-1FC group**: L4 pyramidal neurons with significant positive STTC coupling to R (solid red lines). Dotted red lines from adjacent yellow stars indicate partially overlapping L4-1FC groups of neighboring L2/3 neurons; blue lines denote R's L2/3-1FC partners. Because calcium frames are ~158.7 ms long, STTC was generally computed at Δt = 0 (nondirectional), and we therefore make no directional information flow claims. Nevertheless, STTC at Δt = 0 identifies cross-layer neuron groups whose activity is synchronized above chance, defining putative interlaminar communication pathways. **(B)** Notably, a *directional* STTC analysis at a *two-frame lag* (~317 ms) does show a **positive bias in incoming** (L4→L2/3) minus outgoing (L2/3→L4) links (histogram), consistent with a predominantly feedforward drive from L4. For the directional STTC, we broadened the window of synchrony to 2 additional imaging frames, taking into consideration the temporal order of occurrence of spikes in A relative to B, i.e. reflect the probability that spikes of one neuron may systematically precede (or follow) spikes of the other (see the Methods for a more detailed description). **(C)** Mean L4-1FC group size (# of neurons) by quartile (Q1–Q4) for each mouse (M1–M5). **(D)** Neuron counts per layer (L4, L3, L2) imaged in each mouse.

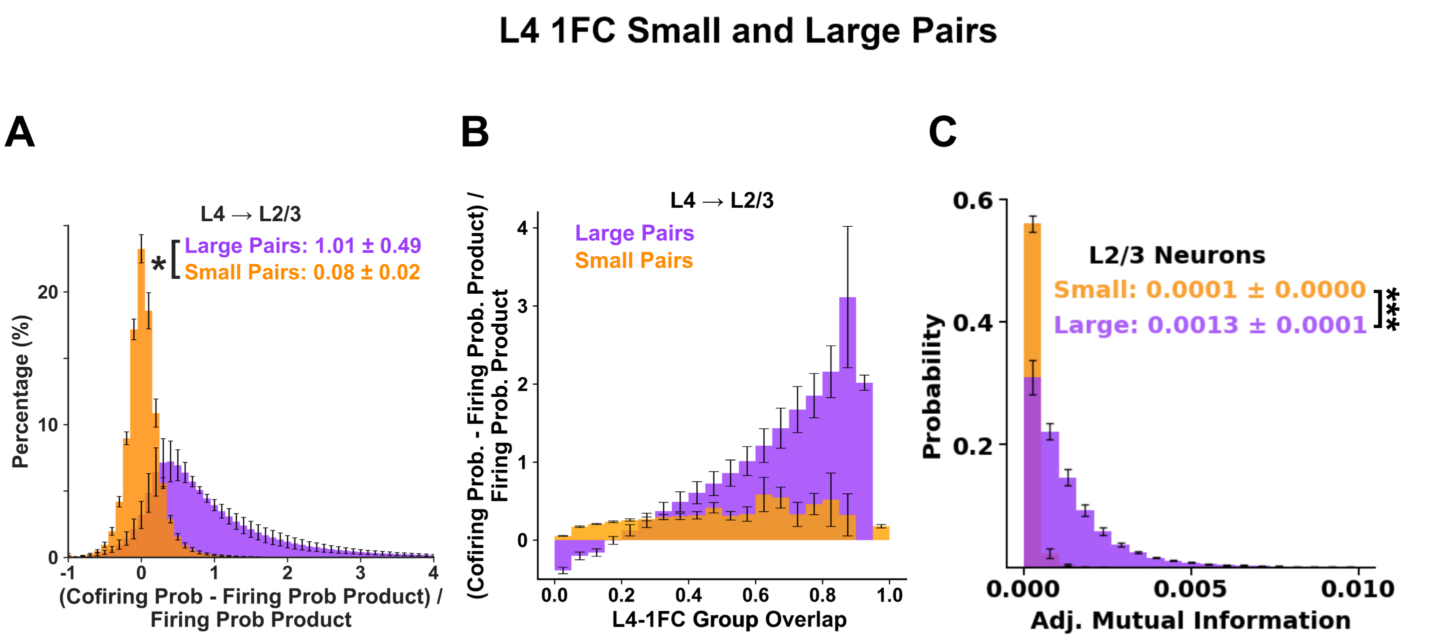


**Supplementary Figure 17: L2/3 inter-neuronal correlations depend on L4-1FC size.** L2/3 pyramidal neurons with small L4-1FC groups tend to fire *independently* during spontaneous activity, whereas neurons with large groups are highly correlated. L4-1FC groups were defined from resting-state recordings, in the absence of stimulus. Panels shown plot L2/3 pairs in which both neurons fall in the lowest degree of connectivity (DoC) quartile (*Q1, small L4-1FC groups*; orange) or the highest DoC quartile (*Q4, large L4-1FC groups*; purple). **(A)** Histogram of the normalized cofiring excess, $(P_{12}-P_1 \times P_2)/(P_1 \times P_2)$, demonstrating that for Q1 pairs cofiring excess is near zero. In contrast, the cofiring excess distribution for Q4 pairs is shifted to the right and has a long tail. **(B)** Same metric stratified by the fractional overlap between the pair's L4-1FC groups (see Methods). Q4 pairs show higher overlap and larger co-firing excess, whereas Q1 pairs remain near-independently firing even at high overlaps. Note that ≥65% of Q1 pairs have zero L4-1FC overlap, and these have particularly low normalized co-firing excess (range: 0.03-0.07). **(C)** Histograms of adjusted mutual information (see Methods) between resting-state eventograms for Q1 and Q4 pairs, again demonstrating Q1 pair independence. Error bars denote SEM across mice (n=5); inset shows overall mean ±SD of means across mice. P-values: * < 0.05, ** <0.01, *** <0.001. P is taken as the most conservative value across the permutation test, Welch's t-test, and one-way ANOVA (see Methods).

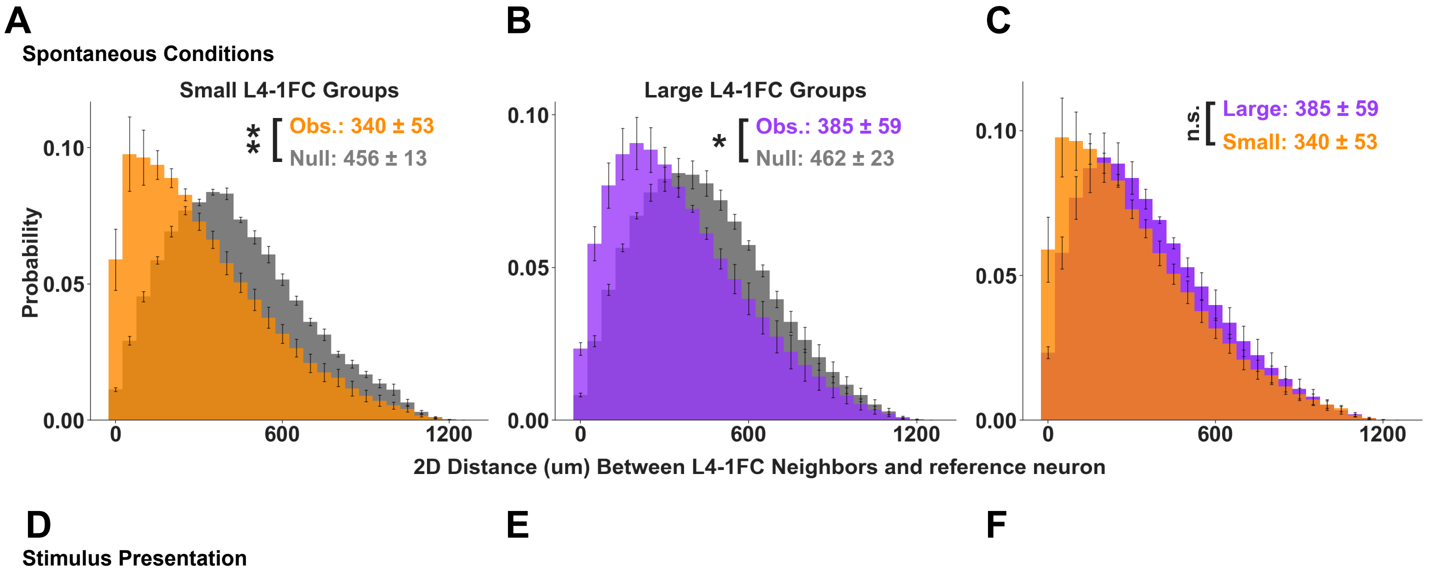

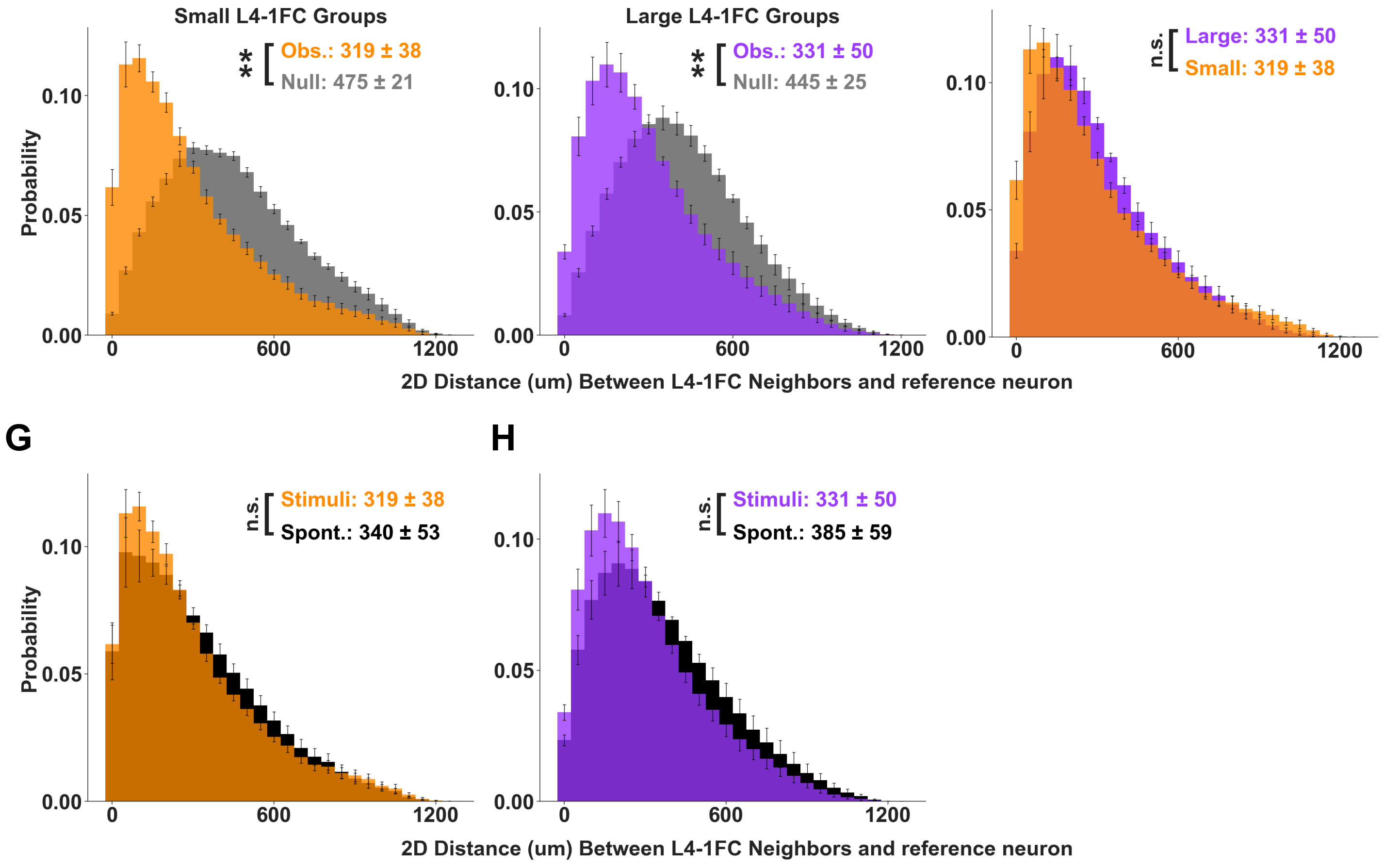


**Supplementary Figure 18 | Spatial extent of L4-1FC groups relative to their L2/3 index neuron.** Two-dimensional (x–y) distances were computed between each L2/3 neuron and all L4 neurons in its 1FC group and compared to size-matched control groups of randomly selected, non-1FC-L4 neurons. **(A–C)** Spontaneous activity: histogram of these distances for small (A) vs large (B) 1FC groups and their comparison (C). **(D–F)** Same analysis during stimulus presentation. **(G,H)** Cross-condition comparison (spontaneous vs stimulus) for small (G) vs large (H) groups. Functionally connected L4 neighbors lie significantly closer to the L2/3 index neuron than controls, with no significant differences between small vs large groups or between spontaneous vs stimulus conditions. Mean ± SD of mouse means; error bars, SEM across mice (n = 5). P-values: "*" < 0.05; "**" < 0.01; and "n.s.": non statistically significant. *P* values are chosen conservatively (maximum across permutation-of-means, Welch's *t*-test, and ANOVA *F*-test; see Methods).

#### Normalization of the impact of the size of the 1FC group in L4 on the prediction of the response of the L2/3 neuron

Let's suppose for simplicity that the L4 connectivity group consists of **N** independent neurons firing each with probability ***p*** within one frame. In this case, the mean firing is **μ=Np** and the firing standard deviation is **σ =** $\sqrt{Np(1-p)}$. If we denote by ***F*** the L4 connectivity group firing random variable, then, by the central limit theorem, we have

$$P\left(f \leq \frac{(F-Np)}{\sqrt{N}} \leq f + \delta f\right) \approx \frac{1}{\sqrt{2\pi p(1-p)}} e^{-\frac{f^2}{2p(1-p)}} \delta f$$

What is important about this distribution is that it is ***independent*** of ***N***. Hence if L2/3 neurons have response probability proportional to the normalized group firing above the mean, $p_R = \alpha\left(\frac{F-Np}{\sqrt{N}}\right) + b$, for the same constants ***a, b,*** they would respond under the same distribution ***irrespective of the size of their L4 group***, permitting a more homogeneous response in L2/3. In this case, the slope of the L2/3 neuron response against their L4 1FC group cofiring normalized by $N^{\frac{1}{2}}$ would be the same irrespective of the L4 1FC group size. This is slightly less than the 0.6-0.7 found experimentally. This difference might be attributed to either the existence of correlations between L4 neurons or differences in the probability of firing between different neurons in L4.

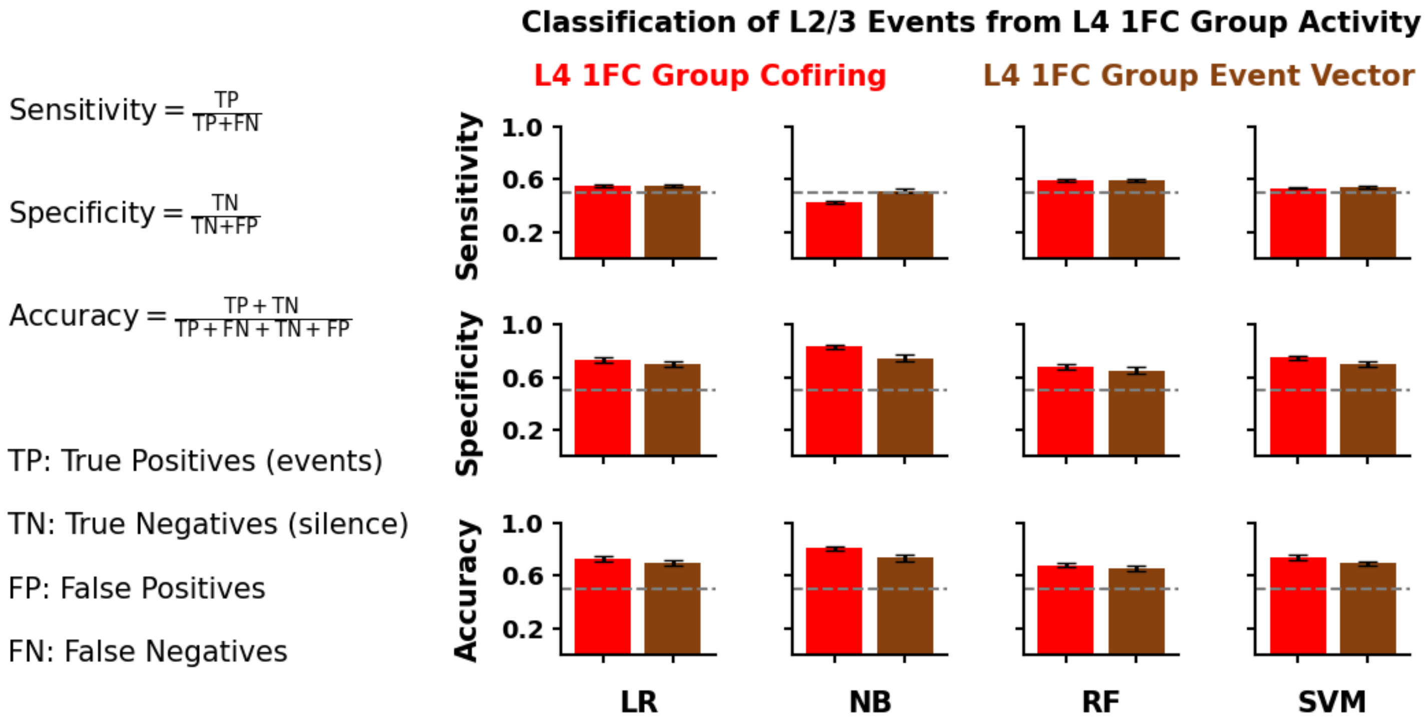


**Supplementary Fig. 19 | Prediction of L2/3 events from L4-1FC group cofiring activity during spontaneous conditions.** We trained four classifiers—logistic regression (LR), Gaussian naïve Bayes (NB), random forest (RF), and support vector machine (SVM)—to predict L2/3 neuron firing per imaging frame, using two input representations: **(i)** the number of cofiring events within the neuron's L4-1FC group (red), and **(ii)** the identity-resolved L4-1FC activity vector, which corresponds to the binary firing pattern across all L4-1FC group members (brown). Performance, reported as sensitivity, specificity and accuracy (see Methods), was evaluated with a 70/30 train/test split of each one-hour long recording: 1FC groups were computed from the first 40 min (training epoch) and performance was tested on the last 20 minutes (test epoch). Analysis included neurons with L4-1FC size > 15, the training set was class-balanced (firing versus no firing frames) by subsampling non-firing frames. Error bars show SEM across mice (n=5).

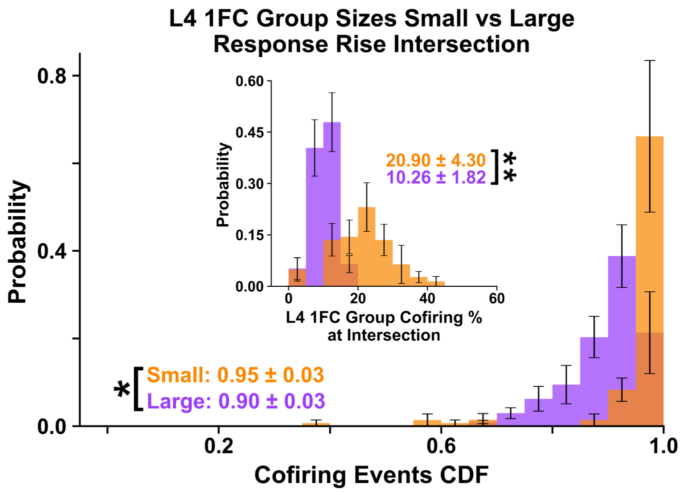


**Supplementary Figure 20| Transition point between two L2/3 response regimes as a function of L4-1FC group size.** Histogram of the **cumulative probability** of L4-1FC group cofiring at the point where an L2/3 neuron's response switches between its two response regimes, shown for neurons with **small** L4-1FC groups (1st quartile; orange) versus **large** L4-1FC groups (4th quartile; purple), as in Figures. 6C and 3G-H. For each L2/3 neuron, response probability (y-axis) was plotted against the regime-transition CDF value (CDF probability (x-axis) computed over the number of cofiring events in its L4-1FC group (defined under resting-state conditions) and fit with **two straight lines**; their intersection defines the regime-transition CDF value). *Inset:* histogram of the corresponding **L4-1FC cofiring percentage** at the transition point. Summary statistics report mean ± SD of mouse means; error bars denote SEM across mice (n = 5). Significance: P-values: * < 0.05, ** < 0.01, "n.s.": non statistically significant. Reported *P* values are conservative (maximum across permutation-of-means, Welch's *t*-test, and ANOVA *F*-test).

#### 6. Dependence of functional connectivity on brain state

The dynamic changes observed in the **pupillary diameter** over time have been associated with the state of alertness and attentional effort of the animal. **Aggregate population** activity in neuronal networks is also thought to reflect the general state of alertness/attention. We, therefore, asked how functional connectivity is modulated by pupillary and aggregate population activity dynamics, parameters that reflect the animal's internal state. We first present the neuronal firing (calcium event rate) at different states, namely during epochs of low population firing vs. high population firing activity, and focus on L2/3 neurons with different L4-1FC group sizes. We demonstrate that the distribution of *functional connectivity weights* under spontaneous conditions remains relatively invariant across different aggregate activity epochs, while the architecture of the functional connectivity does change.

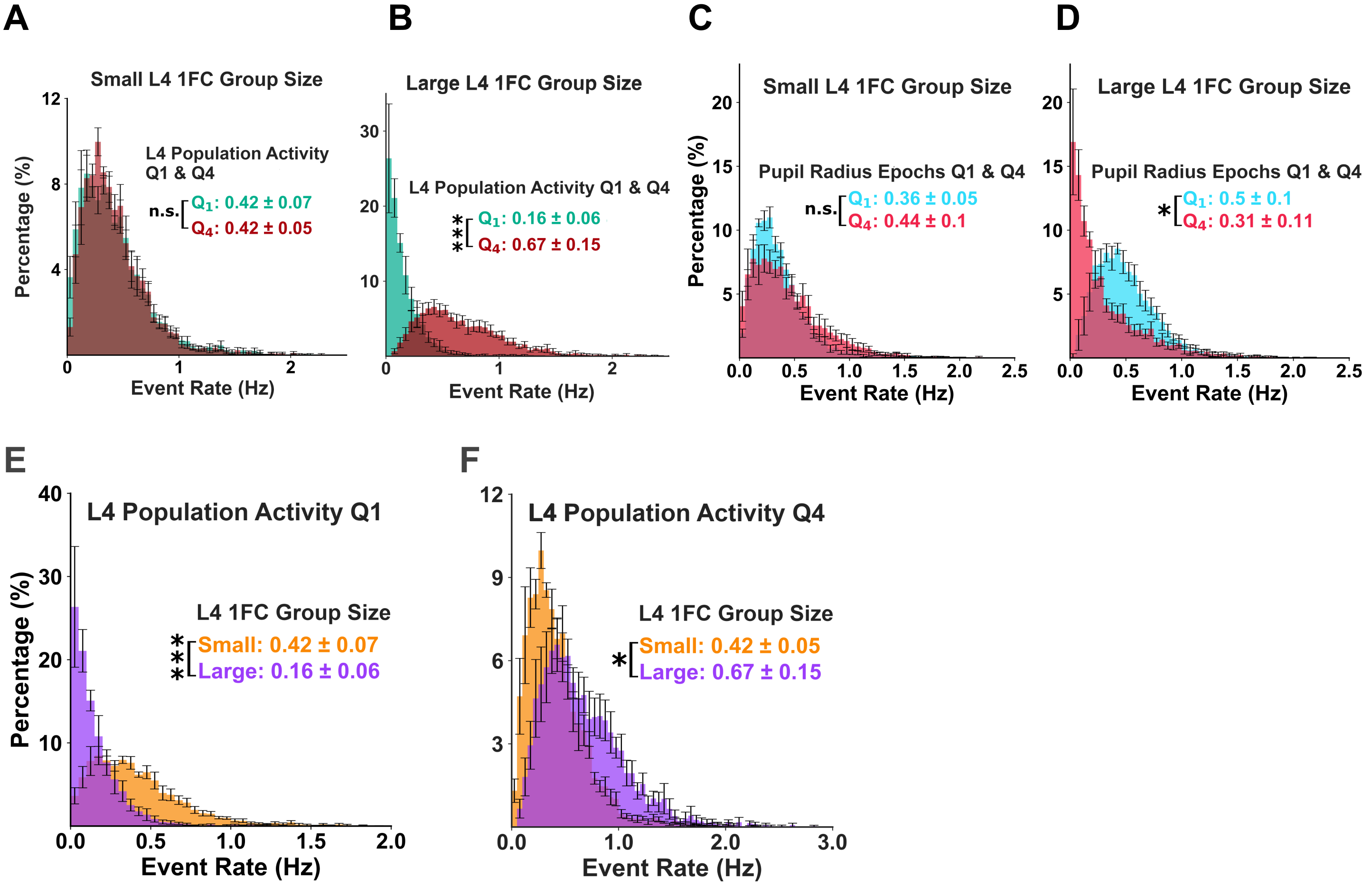


**Supplementary Figure 21: Firing rate of L2/3 neurons depends on L4-1FC group size.** L2/3 neurons that are functionally connected to smaller L4 groups exhibit different firing rates than those of L2/3 neurons functionally connected to larger L4 groups. **A.** Histogram of mean calcium event rates (Hz) of L2/3 neurons with small L4 1FC groups (0-25 percentile of L4-1FC group sizes) under two conditions: 1) low aggregate L4 population activity (colored green-turquoise; 0-25 percentile of overall population activity) and 2) high aggregate L4 population activity (colored brown-maroon; 75-100 percentile of aggregate activity). Note the distribution of event rates is essentially identical under the two conditions. **B.** Similar to (A), now for the mean calcium event rates (Hz) of L2/3 neurons with large L4-1FC groups (75-100 percentile of L4-1FC group sizes). Note that in contrast to neurons with small groups the distribution of event rates for L2/3 neurons connected with large L4 groups shifts significantly to the right (higher values) when the overall population activity increases (Q1 vs Q4). **C.** As in (A), but comparing the calcium event rate of L2/3 neurons with small L4-1FC groups (0-25 group size percentile) under different epochs of pupil radius size: 1) low pupil radius (0-25 percentile; cyan), versus 2) high pupil radius (75-100 percentile; pink). Note that the calcium event rate distributions remain similar, and their means are not significantly different.

**D.** As in (C) for the calcium event rates of L2/3 neurons with large L4-1FC groups (75-100 group size percentile). In contrast to L2/3 neurons with small groups, the firing-rate distribution of L2/3 neurons connected with large L4-1FC groups is higher in small pupil radius epochs. **E.** When aggregate activity in L4 is low (Q1: 0-25%, i.e. first quartile) the histogram of mean event rates (Hz) for L2/3 neurons with large L4-1FC groups (purple; Q4: 75-100 percentile of L4-1FC sizes) is to the left (towards smaller values) of that of L2/3 neurons with small L4-1FC groups (orange; Q1: 0-25 percentile of L4-1FC sizes). **F.** Conversely, when aggregate activity in L4 is high (Q4: 75-100%, i.e. highest aggregate firing rate quartile), the firing rate histogram of L2/3 neurons with large L4-1FC groups shifts to the right, at higher values. Note that the histogram of mean event rates (Hz) of L2/3 neurons with small L4-1FC groups (orange; Q1: 0-25 percentile of L4-1FC sizes) remains approximately independent of aggregate layer 4 activity. In Figs. A-F the L4-1FC groups as well as the aggregate activity are estimated under resting state conditions. Inset values reflect mean ± standard deviation of the sample means across mice (n=5). Error bars correspond to SEM across mice (n=5). "*": p-value < 0.05; "***": p-value < 0.001 and "n.s.": non statistically significant. The highest p-value obtained from the permutation of means, the Welch's t-test, and the ANOVA F-test is considered for the level-of-significance (see Methods for more details).

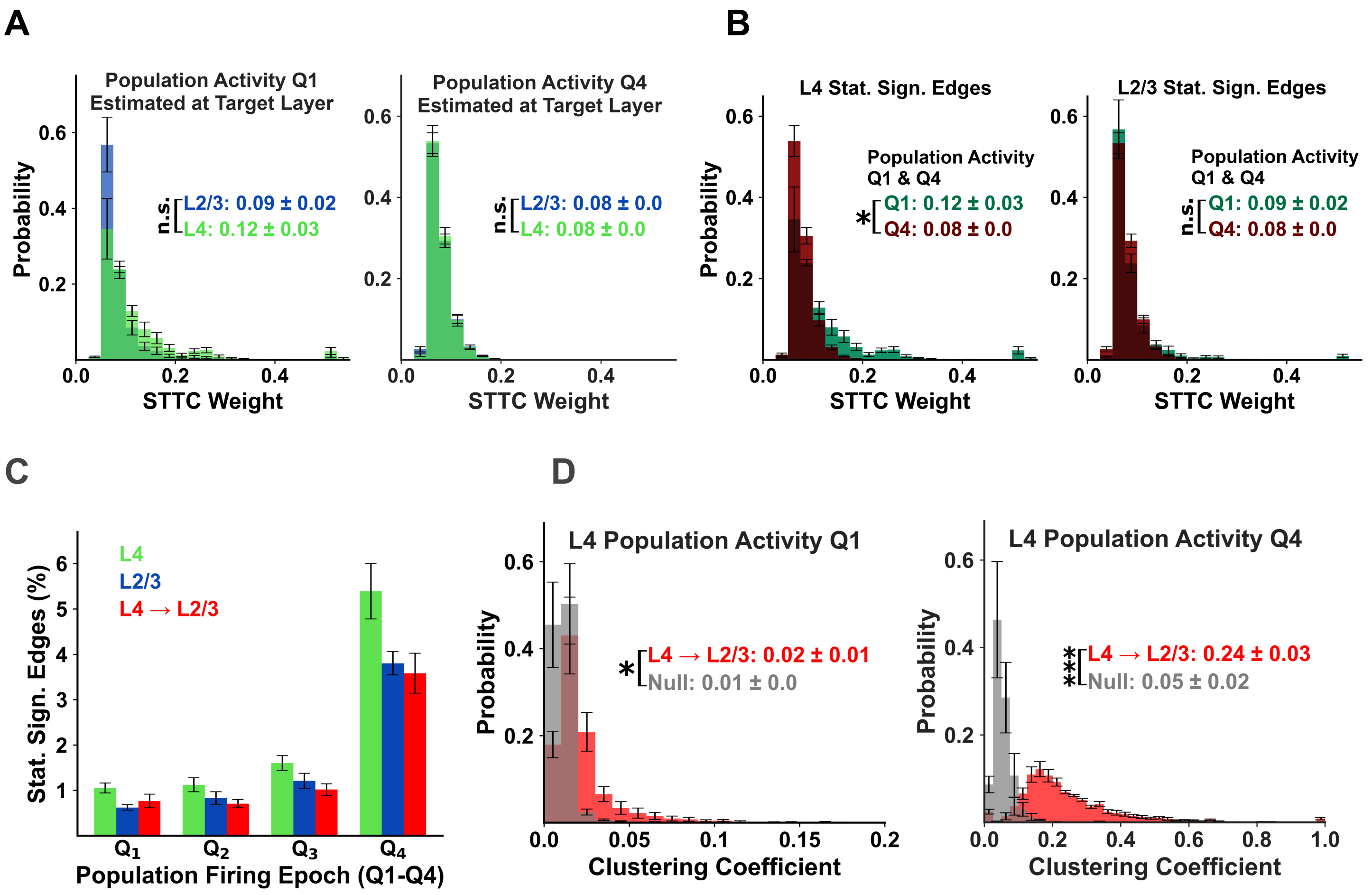


**Supplementary Figure 22: Functional connectivity in resting state across different aggregate activity epochs. A-B)** The distribution of functional connectivity ***weights*** under spontaneous conditions remains relatively invariant across different aggregate activity epochs. **A.** Histogram of STTC weights of functional connections, which are statistically significant during the epochs of low (Q1) vs. high population activity (Q4) quartiles. L2/3 vs L4 intra-layer connection weights are compared. During the epoch of high population activity, the distribution of STTC weights is identical across layers, whereas during low population activity STTC weights in L4 are slightly higher. **B.** Similar to A, except now the histograms across the low (Q1, dark green) vs the high (Q4, brown) quartiles of activity are compared within each layer. For L2/3 (right), the distributions are identical, while for L4 (left), the weight distribution has a slightly longer tail towards higher values when the aggregate activity is low. It is important to note that although the number of statistically significant edges and magnitude of clustering coefficients changes significantly and by a considerable percentage, the distribution of functional connectivity weights remains relatively invariant across different aggregate activity epochs. **C-D) The architecture of the functional connectivity during epochs of different aggregate firing activity. C.** Percentage of statistically significant edges calculated during epochs of different population firing activity quartiles (Q1-Q4) under spontaneous conditions. Intra-layer L4 (green) and L2/3 (blue) as well as inter-layer L4->L2/3 (red) statistically significant functional connection (i.e., edge) percentages are presented. As expected, in epochs of high aggregate activity the percentage of significant functional connections increases in all compartments. **D.** Left: Histogram of the clustering coefficients of L4-1FC groups of L2/3 neurons, computed during the low L4 population activity quartile (Q1). The 1FC neighbors of a reference neuron are computed during the entire resting-state period however the statistically significant edges were considered based on their values at the specific epoch of interest (Q1, Q4). Right: Similarly for the highest L4 population activity quartile (Q4). It is clear that clustering coefficients are much higher during the high aggregate activity quartiles. Inset values reflect mean ± standard deviation across mice (n=5). P-values: "*"<0.05; "n.s.": non statistically significant. The highest p-value obtained from the permutation of means, the Welch's t-test, and the ANOVA F-test is considered for the level-of-significance (see Methods for more details).

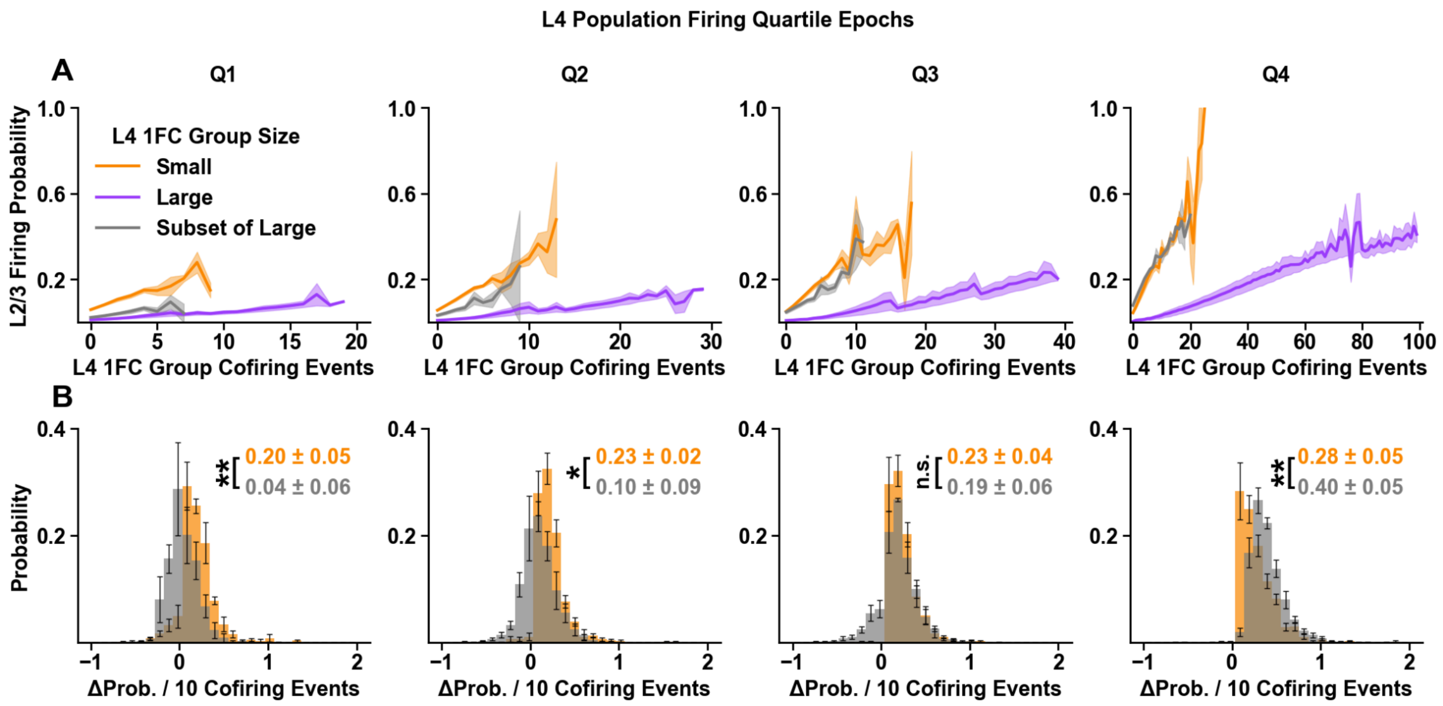


**Supplementary Figure 23. Small L4-1FC group properties are not explained by subsampling large groups.** Resting-state response functions were computed as L2/3 firing probability versus within-frame co-firing count in the associated rL4-1FC group, stratified by rL4-1FC size quartiles: *small* (Q1, orange; 0–25%) and *large* (Q4, purple; 75–100%). Panels (left to right) show epochs binned by *aggregate L4 population activity, Q1 (low) to Q4 (high)*. **(A)** Solid lines: empirical response functions for Q1 (orange: small groups) and Q4 (purple: large groups). *Gray lines*: subset (control) responses of Q4 neurons (neurons with large groups) whose responses are plotted as a function of the co-firing within randomly selected subsets of their rL4-1FC groups, with subset size matched to that of Q1 neurons (neurons with small groups). Note that actual small-group L2/3 neuron response functions (orange line; Q1) differ from the subset control (gray line), particularly during low-to-moderate population activity (L4 activity quartiles: Q1–Q3), converging together only at high activity (activity quartile Q4; rightmost plot). **(B)** Distributions of ReLU-fit slopes (good fits only, $R^2 \geq 0.8$) comparing true small groups (orange) to subset controls (gray), showing significant differences during L4 activity quartiles Q1–Q3. Insets, mean ± SD of mouse means; error bars and shading, SEM across mice (n = 5). Significance: permutation of means, Welch's *t* test, and ANOVA *F* test (largest *P* reported); *: $P < 0.05$, **: $P < 0.01$, n.s.: not significant.

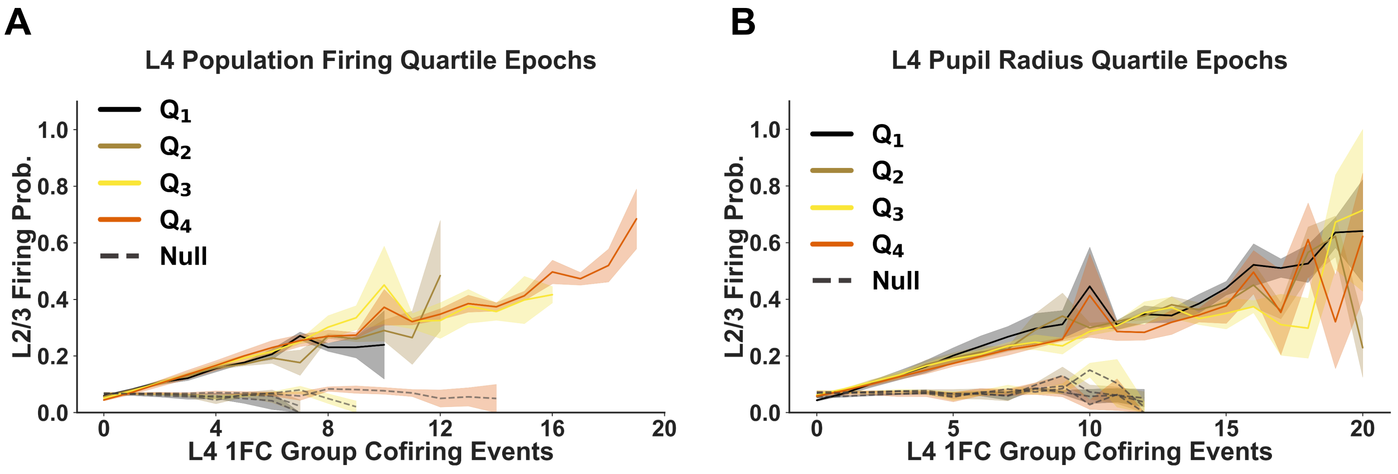

**Supplementary Figure 24: Resting-state L2/3 response slopes vary little with aggregate L4 population activity or pupil size.** L2/3 neuron firing probability is plotted as a function of the within-frame co-firing count of the neuron's rL4-1FC group (defined during spontaneous activity). **(A)** Response functions binned by aggregate L4 activity quartiles (Q1-low to Q4-high). L2/3 neuron response functions during epochs of low (quartile Q1) to high (Q4) aggregate L4 activity show similar slopes. **(B)** Response functions binned by pupil-size quartiles (Q1–Q4; 0–25% to 75–100%) likewise overlap, indicating minimal dependence on pupil-linked state. Shading indicates SEM across mice (n = 5). Plots show L2/3 neurons with small L4-1FC groups (size-quartile Q1); comparable slope stability is observed for large groups (not shown).

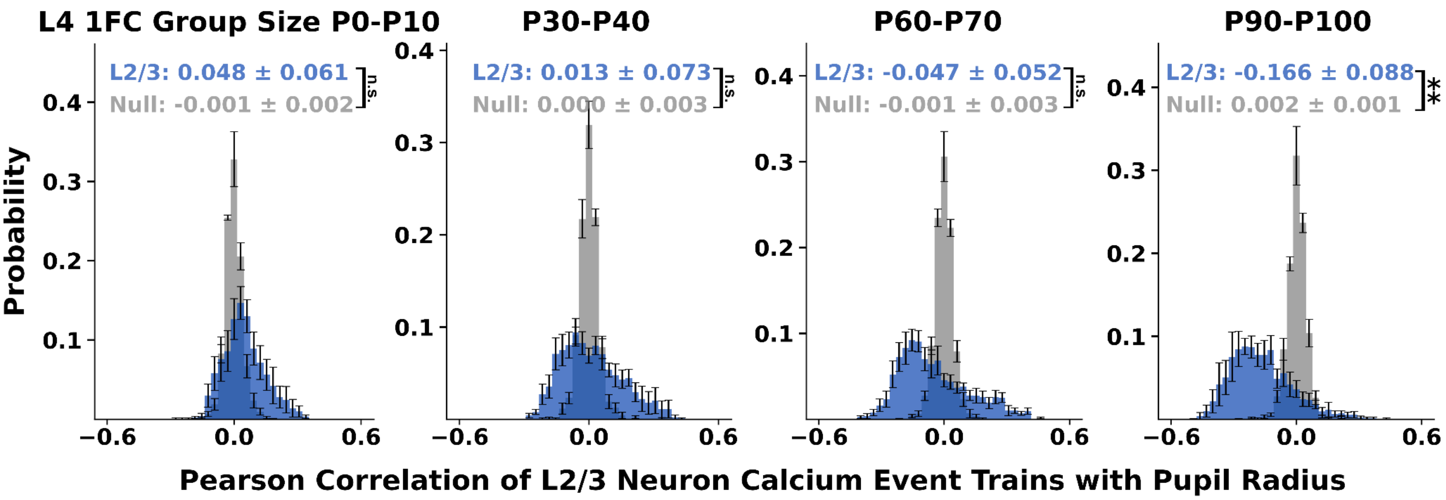


**Supplementary Figure 25: Correlation of calcium eventograms of L2/3 neurons with different L4-1FC group sizes with pupil radius.** Histogram of the Pearson correlation of the calcium eventogram of an L2/3 neuron (binned in 5-frame intervals) with the pupillary radius, considering L2/3 neurons that are connected with L4 1FC groups of different sizes, namely in the decile [0, 10]% (Q0-Q10), [30, 40]% (Q30-Q40), [60, 70]% (Q60, Q70), and [90, 100]% (Q90, Q100). Only frames of *quiet wakefulness* were considered here. Similar results were obtained during the entire spontaneous period. Error bars represent the standard error of the mean across mice, while the statistics on the plot indicate the mean ± standard deviation of the sample means across mice (n=5). P-values: "**"<0.01; "n.s.": non statistically significant. The highest p-value obtained from the permutation of means, the Welch's t-test, and the ANOVA F-test is considered for the level-of-significance (see Methods for more details).

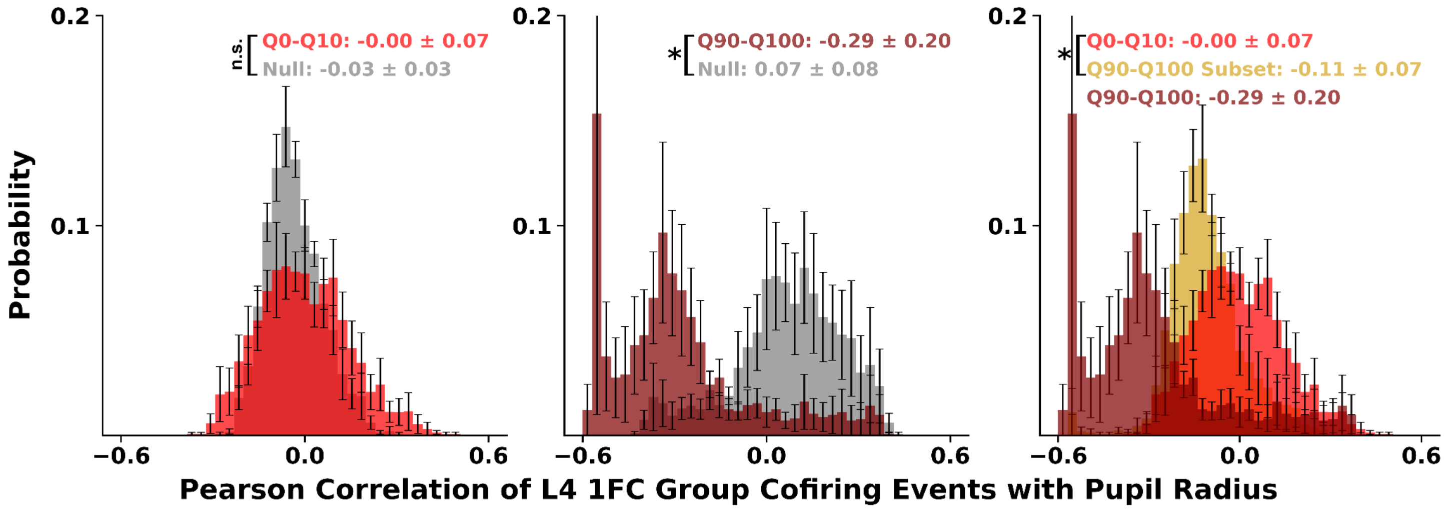


**Supplementary Figure 26| Pupil-size linked modulation of L4-1FC group cofiring depends on group size.** Histograms of Pearson correlation coefficients ($r$) between pupil radius and L4-1FC group activity (defined as the per-frame sum of cofiring events across all L4 neurons in the L4-1FC group associated with each L2/3 neuron). Correlations are shown for L2/3 neurons with **small** L4-1FC groups (0–10th percentile; bright red; left panel),

**large** groups (90–100th percentile; dark red, middle panel), and **size-matched subsets** sampled from the large groups to match the small-group size (yellow; right panel). Larger group cofiring is anticorrelated with pupil radius, and size-matched subsets retain the anticorrelation, resulting in a distribution that is distinct from that of genuinely small groups (mean average correlation ~0; right panel). Error bars, SEM across mice (n = 5); inset, histogram mean ± SD of means across mice. Significance: P-values: "*"*<0.05;* "n.s.": non statistically significant. Reported $P$ values are conservative (maximum across permutation-of-means, Welch's $t$-test, and ANOVA $F$-test; see Methods).

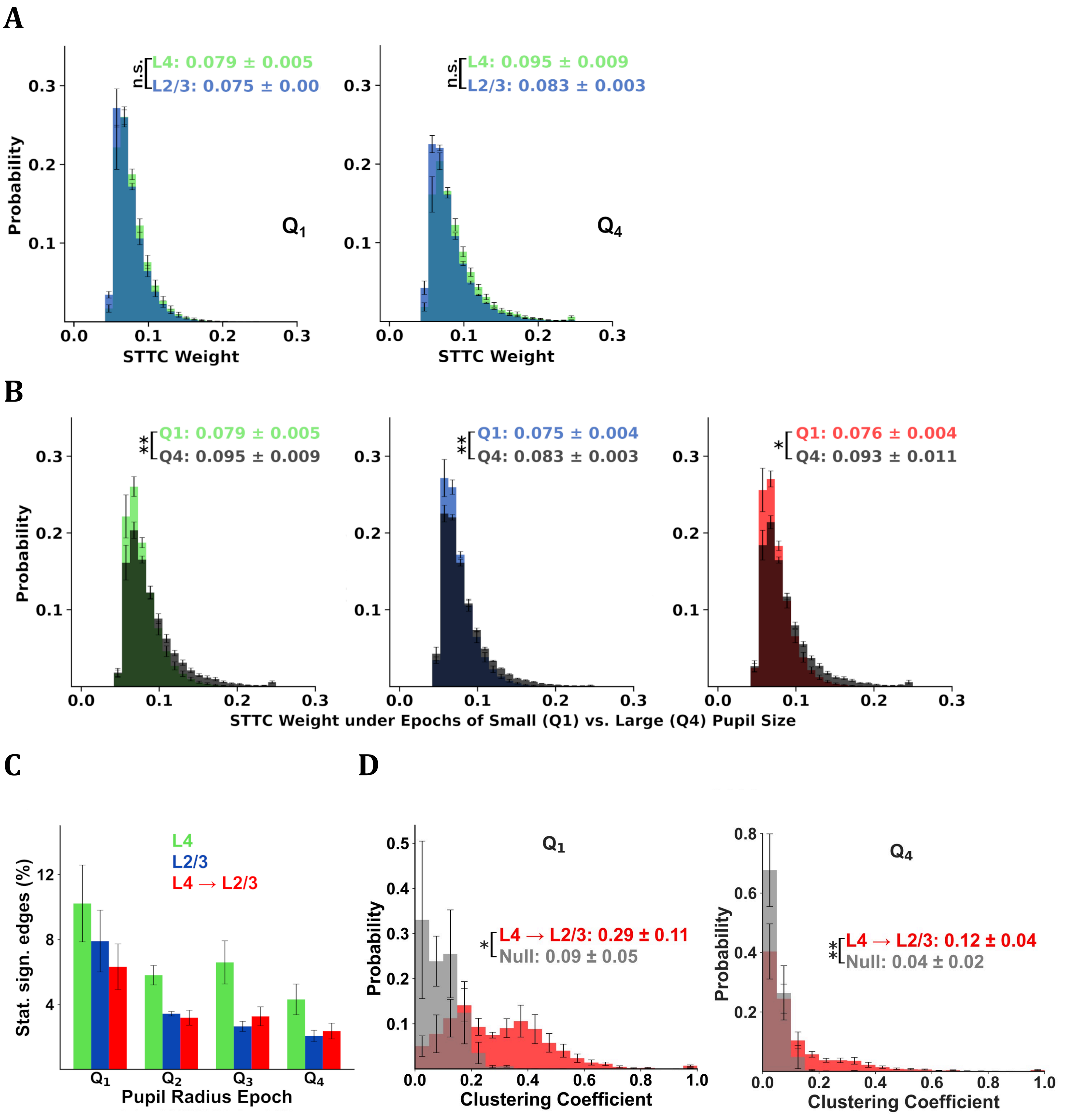


**Supplementary Figure 27: Epochs of smaller pupil size yield a higher percentage of functionally connected pairs under spontaneous conditions. A)** *Left:* Histogram of STTC weights of statistically significant functional connections within L4 (green) versus L2/3 (blue) measured during the epochs of small pupillary radius (Q1: first quartile, corresponding to 0-25% lowest values of pupillary radius). *Right:* as in left but during epochs of high pupillary radius (Q4: $4^{th}$ quartile, corresponding to 75-100% highest values of pupillary radius). As can be seen, the STTC weight distributions in L2/3 vs L4 are similar in either condition. **B)** Histograms as in (A), comparing epochs of small (Q1; colored) versus big (Q4; black) pupillary radius quartiles. *Left to right panels:* L4, L2/3, and L4 to L2/3. Functional connectivity distributions are slightly shifted to higher weights at epochs of high pupillary radius in all cases. **C)** Percentage of statistically significant edges during the different quartiles of pupillary radius, Q1-Q4, for the intra-layer cases L4 and L2/3 as well as from L4 to L2/3. The higher the pupillary radius the *fewer* significant functional connectivity pairs are found both within and across layers. **D)** *Left:* Histogram of the clustering coefficients in L4-1FC groups of L2/3 neurons computed using the connections that were statistically significant during the low pupillary radius epoch Q1. *Right:* Similarly for the high pupillary radius epoch Q4. For the purposes of defining the L4-1FC group, functional connectivity was computed across the entire 60-minute spontaneous imaging period (i.e. not distinguishing between epochs of high or low pupillary radius). Overall, there is a sharp increase in the percentage of functional connectivity edges and clustering coefficients during small pupillary size epochs. This goes together with only a small decrease in the magnitude of functional connectivity weights. Inset values indicate the mean ± standard deviation of the sample means across mice, while error bars correspond to SEM across mice (n=5). P-values: * < 0.05, ** < 0.01, "n.s.": non statistically significant. The highest p-value obtained from the permutation of means, the Welch's t-test, and the ANOVA F-test is considered for the level-of-significance (see methods for more details).

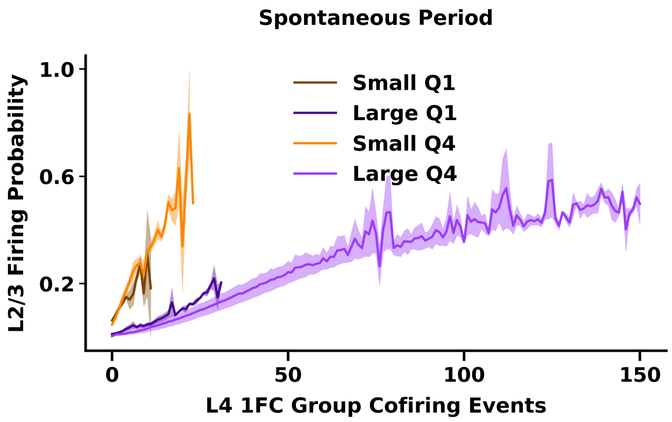


**Supplementary Figure 28 | L2/3 response functions are weakly, if at all, modulated by the global population firing state during spontaneous activity.** Response probability of L2/3 neurons is plotted versus L4-1FC group cofiring count, computed separately during **high** (Q4; 75-100%) versus **low** (Q1; 0-25%) aggregate-population firing epochs. *Group size strongly shapes the slope of the response*: contrast responses of small (orange/brown) versus large (dark/light-purple) groups. In contrast, within the same range of group sizes, slopes remain similar across markedly different population firing states: contrast yellow vs brown curves for small group sizes to light vs dark purple curves for large group sizes. High-firing epochs mainly extend responses to higher probabilities as they exhibit larger cofiring counts. Shading, SEM across mice (n = 5).

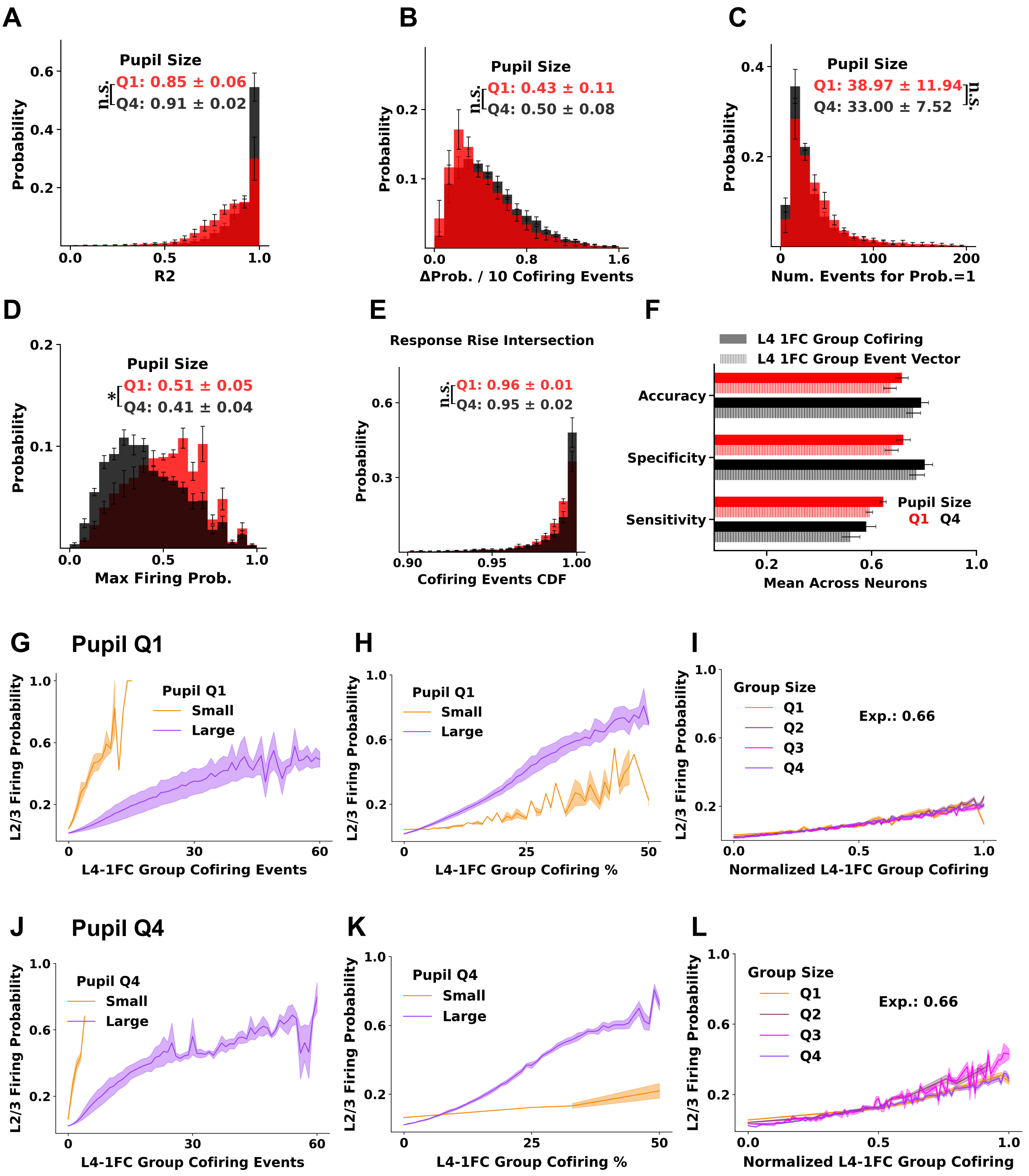


**Supplementary Fig. 29 | The relationship between L2/3 neuronal responses and L4 ensemble cofiring is preserved across alertness states during resting state. (A–E)** ReLU-based characterization of L2/3 responses as a function of L4-1FC cofiring during epochs of **small pupil size** (1st quartile: Q1, red) versus **large pupil size** (4th quartile: Q4, black). ReLU fits were appropriately weighted by the standard error of each point. **(A)** Distribution of ReLU fit quality ($R^2$). L4-1FC groups were defined within the corresponding pupil-size epoch. **(B)** Distribution of response slopes (increase in firing probability per 10 additional cofiring events per frame) for fits with $R^2 \geq 0.8$. **(C)** Extrapolated number of cofiring events required to reach firing probability = 1 (for fits with $R^2 \geq 0.8$). **(D)** Actual maximum firing probability of the response function. **(E)** Distribution of the cofiring cumulative distribution function (cdf) values at the onset of the steep response regime ("knee"), defined by the intersection of the two fitted response regimes (neurons plotted had $R^2 > 0.7$). **(F)** Sensitivity, specificity, and accuracy of a linear support vector machine (SVM) in predicting L2/3 firing during small-pupil epochs using as input: either aggregate L4-1FC cofiring counts (solid) or a binary vector encoding the identities of firing L4-1FC neurons (hatched). L4-1FC groups used were computed consistently, with data obtained within the small-pupil (Q1, red) and large-pupil (Q4, black) epochs. **(G–I)** Average L2/3 firing probability as a function of **raw** L4-1FC cofiring (G), **percentage** of the L4-1FC group that cofires (H), and **normalized** cofiring (raw cofiring divided by group size raised to a fitted exponent, which here is 0.66) (I). Neurons are grouped by L4-1FC size quartiles: smallest (Q0–Q25, orange) and largest (Q75–Q100, purple). **(J–L)** Same analyses as **(G–I)**, computed during epochs of large pupil size. Note the similarity of the plots. Across all panels, statistics report mean ± SD of mouse-wise means; error bars denote SEM across mice (n=5).

#### 7. Relation of L4 1FC Groups of L2/3 Neuronal Pairs

This section examines the relation between the overlap of the L4-1FC groups of pairs of L2/3 functionally connected neurons vs. their orientation preference and distance. The L4 1FC groups of functionally-connected L2/3 neurons exhibit much higher overlap than expected by chance, suggesting they share significant (putative) common input from L4.

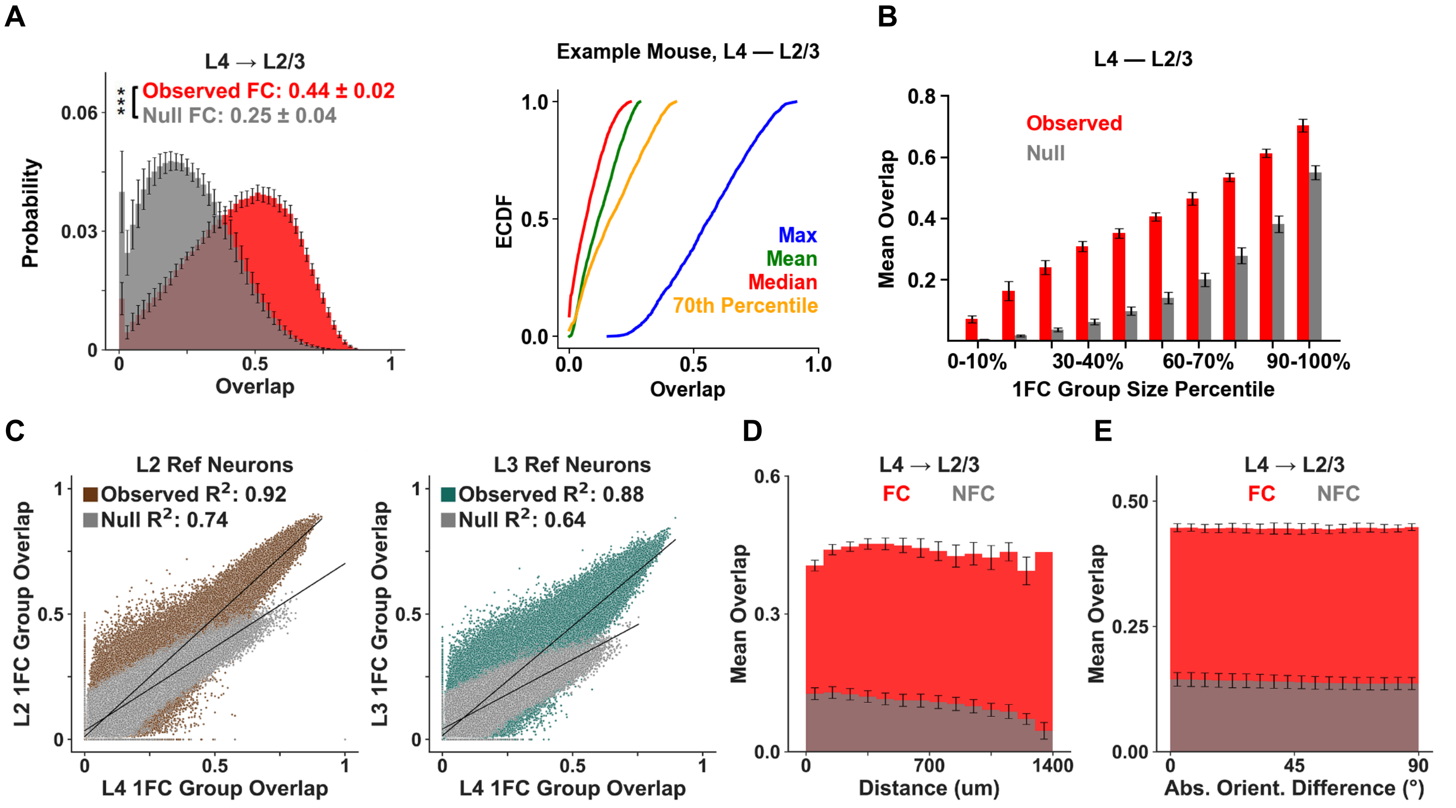


**Supplementary Fig. 30| Shared L4 input predicts functional coupling between L2/3 neurons at rest.**
Overlap between two 1FC groups was quantified as |A∩B|/sqrt(|A||B|). **(A)** *Left:* Functionally connected L2/3 neuron pairs exhibit greater overlap of their observed L4-1FC groups (red distribution) than size-matched null groups drawn from unconnected L4 neurons (gray). *Right*: Distribution of overlap statistics (max, mean, median, 70-percentile overlap) across all L2/3 pairs in a typical example mouse: for each L2/3 neuron, we compute the overlap of its L4-1FC group with the L4-1FC group *of all other* L2/3 neurons and plot the corresponding statistic. Overlaps >80% are rare. **(B)** Mean L4-1FC overlap among L4-1FC groups of the same size, binned by group size. Note percent overlap increases with L4-1FC group size. **(C)** Overlap of L4-1FC groups covaries with the overlap of the corresponding intra-layer L2/3-1FC groups. Note the stronger, steeper, linear correlation for observed pairs

versus controls. **(D)** Mean L4-1FC overlap versus X-Y plane intersomatic distance for same-depth L2/3 pairs: overlap for functionally connected pairs tends to be distance-independent, whereas unconnected pairs show somewhat higher overlap at shorter distances. **(E)** Mean L4-1FC overlap versus absolute orientation difference of the "index" L2/3 neurons for functionally connected L2/3 pairs shows no dependence on tuning difference. Unconnected control pairs, as expected, show also no dependence on tuning difference, but have much lower mean overlap. Plots show mean ± SD; error bars denote SEM across mice (n=5).

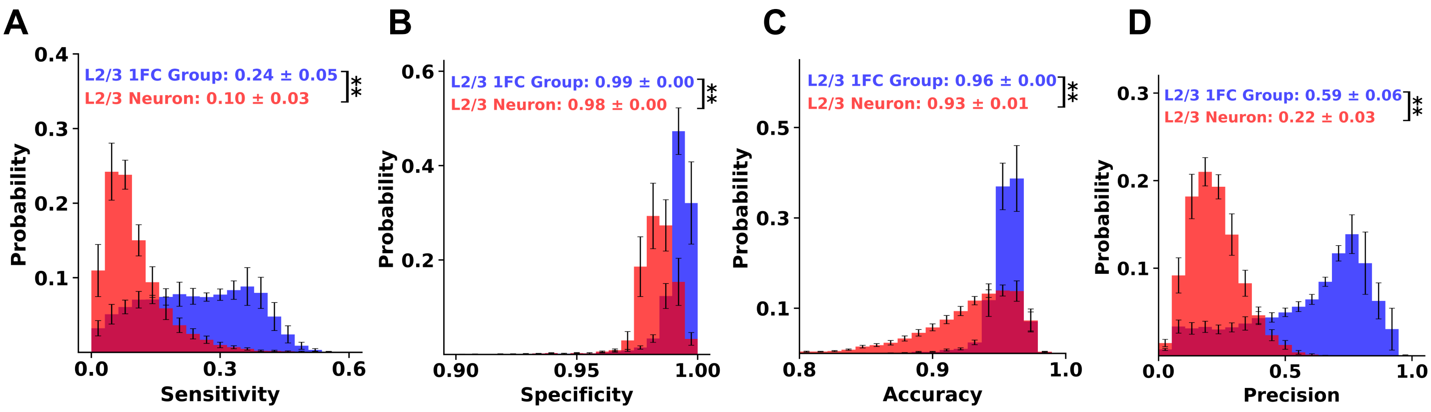


**Supplementary Fig. 31 | Ensemble-to-ensemble outperforms ensemble-to-neuron activity under resting state. A.** Histogram of the sensitivity of activation transfer from the resting-state defined *rL4-1FC groups* of L2/3 neurons to the corresponding *rL2/3-1FC groups* (blue) versus to the *individual L2/3* "index" neuron (red). Sensitivity is defined as the fraction of frames in which the rL4-1FC group is active given that the L2/3 target (group or neuron) is active. *Active frames* are defined as imaging frames in which the group cofiring z-score is above 3 for the L4-1FC group and 2 for the L2/3-1FC group. The latter ensured the number of frames in which rL2/3-1FC groups are active is similar on average to the number of events occurring in individual L2/3-neurons, allowing for a fair comparison. Ensemble-to-ensemble transmission shows markedly higher sensitivity. **B.** Specificity, defined as the fraction of frames in which the rL4-1FC group is inactive given that the L2/3 target is inactive, is likewise higher for ensemble-to-ensemble transmission. **C.** Accuracy, defined as the fraction of frames in which L4 and L2/3 targets are concordant (both active or both inactive). **D.** Precision, defined as the fraction of frames in which the L2/3 target is active given that the rL4-1FC group is active. *Note that across all metrics, ensemble-to-ensemble (rL4-1FC to rL2/3-1FC) communication outperforms ensemble-to-neuron transmission, indicating that interlaminar information transfer under stimulation is more reliably captured at the ensemble level.*

This is consistent with what we observed under stimulation Suppl. Fig. 55**.** Insets report mean ± SD across mice (n=5); error bars denote SEM. P-values: "*"< 0.05, "**"< 0.01,"***"< 0.001; "n.s.": non statistically significant. The highest p-value obtained from the permutation of means, the Welch's t-test, and the one-way ANOVA F-test is reported as the level-of-significance. Note that the results we report here are robust to an extensive sensitivity analysis performed over a *range of different thresholds*.

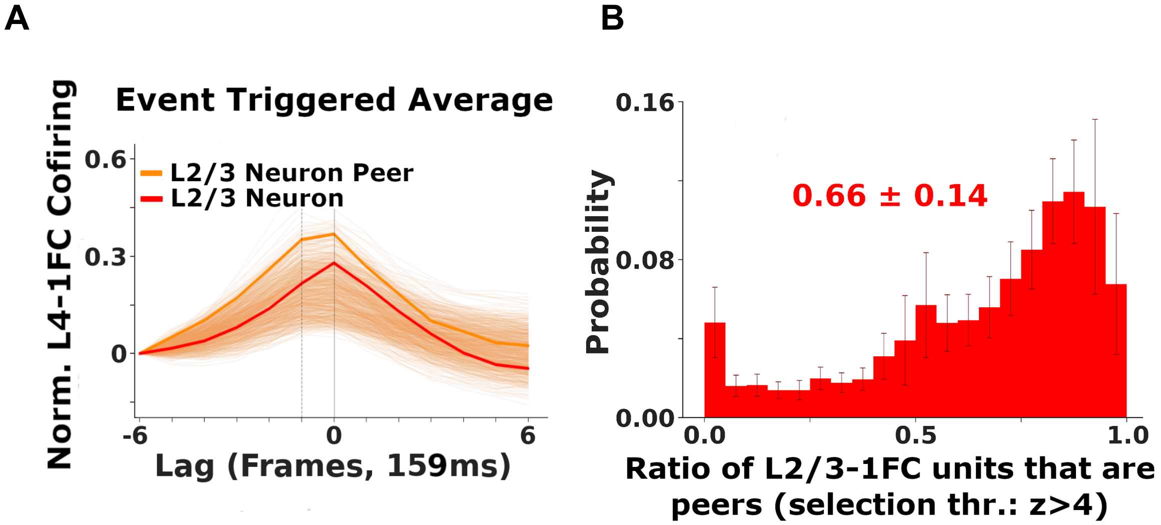


**Supplementary Figure 32 | L2/3-1FC neuron firing is time-locked to the index L2/3 neuron's L4-1FC group cofiring. (A)** For a given reference L2/3 neuron (say *R)*, another L2/3 neuron (say *P*) was defined as a "***peer"***, if its event-triggered average (ETA), computed relative to cofiring events in R's L4-1FC group, exhibited a significant peak (**z** > 4 vs baseline) at lag ≤ 0 (at or before P's firing). Example ETAs are shown (red, R's L4-1FC cofiring; gold, strongest peer; brown, remaining peers). **(B)** Fraction of R's L2/3-1FC group (L2/3 neurons functionally-connected to R) that also satisfies the peer criterion (**z** > 4 for both), revealing substantial overlap consistent with L4–L2/3 information-coupled ensembles.

#### 8. Impact of Global Modes of Activity on Functional Connectivity

The impact of the behavior on functional-connectivity was re-examined using PCA, based on the edges that are statistically significant **after the removal of the first two principal components** (that are strongly correlated with the population and pupil size), demonstrating that the central results remain essentially identical.

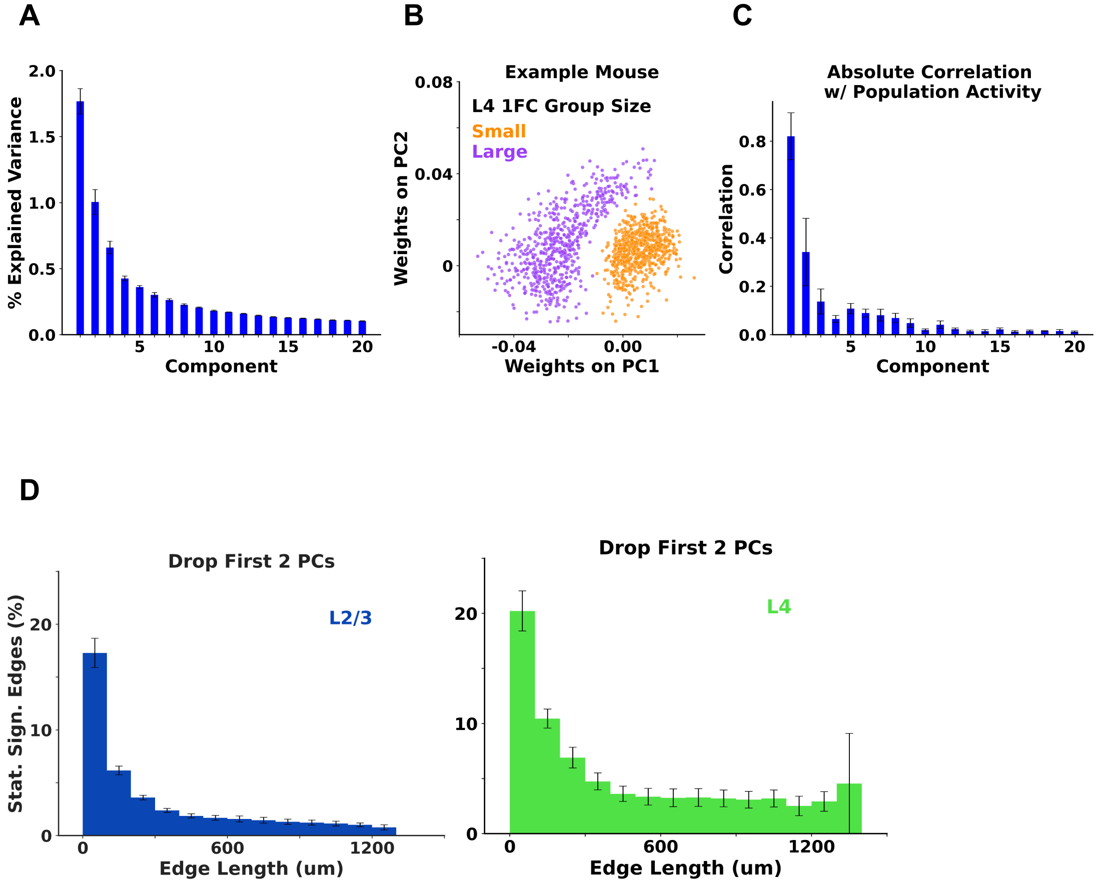

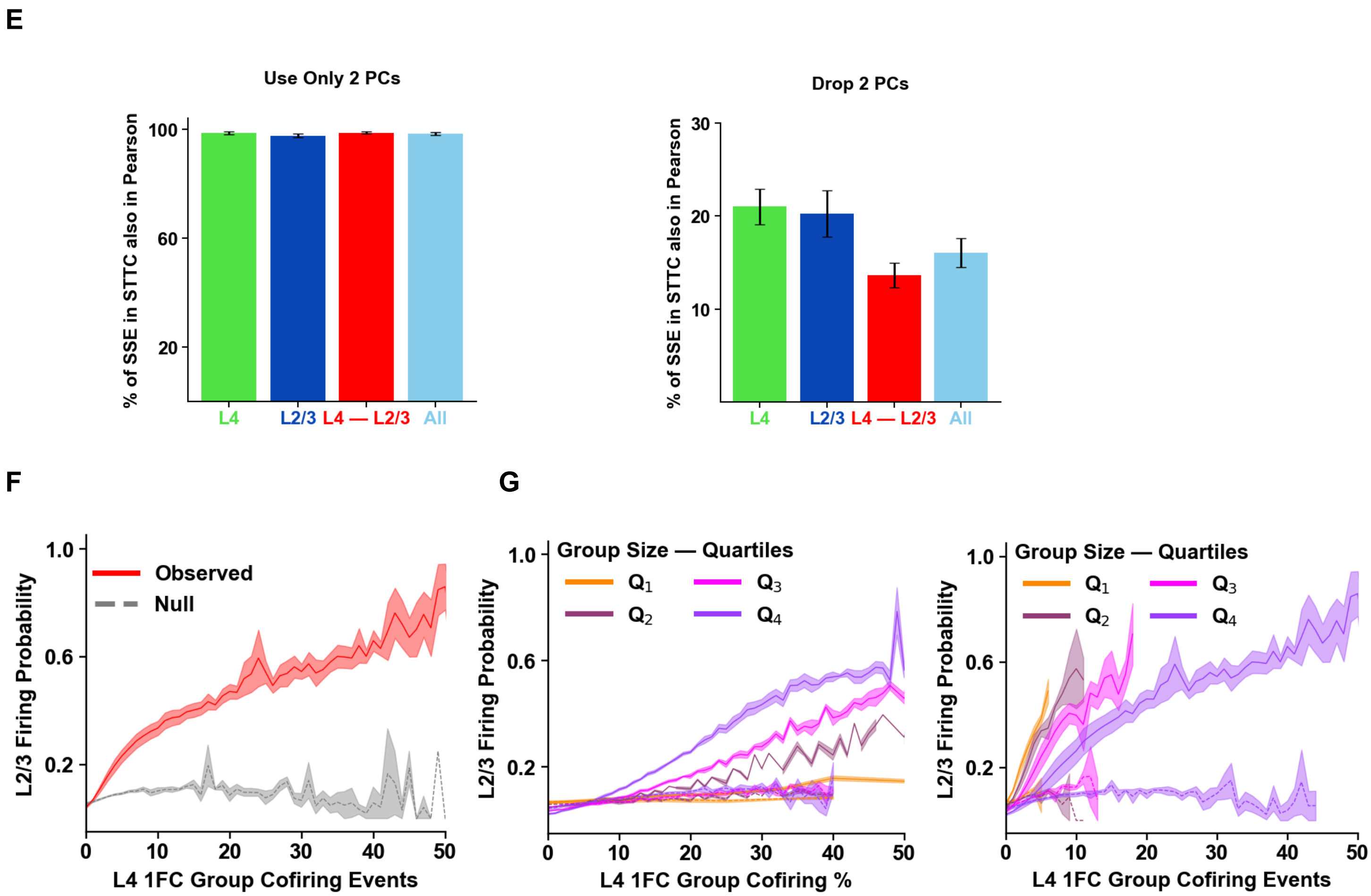


**Supplementary Fig. 33 | Removing global correlation structure preserves main results. (A)** Principal Component Analysis (PCA) performed on resting-state eventograms from V1 L4 and L2/3 shows a dominant first principal component (PC). Variance explained is shown for PCs 1–20. **(B)** L2/3 neurons with small (Q1; orange) versus large (Q4; purple) L4-1FC groups separate in the space of PC1–PC2 (shown for one example mouse; similar across mice), chiefly because of different PC1 projections. **(C)** PC1–PC2 are the components most correlated with global population activity and pupil radius respectively. **(D)** Distance-dependent fraction of significant positive edges ($z > 4$) computed using Pearson correlations after reconstructing signals with PC1–PC2 removed, thereby attenuating population/pupil-linked correlations. **(E)** Overlap between STTC-defined significant edges and Pearson-defined significant edges using either the full reconstruction (retaining PC1–PC2) or the residual reconstruction (PC1–PC2 removed) (stat. sign. definition based on z-score > 4). **(F, G)** L2/3 response functions (as in main Fig. 4A), computed with respect to L4-1FC groups defined from the residual (PC1–PC2-removed) Pearson network. reproduce the manuscript's trends despite reduced edge counts and show clear separation from the null. Error bars/shading, SEM across mice ($n = 5$).

#### 9. Quiet Wakefulness versus Locomotion Periods

Another surrogate marker indicative of **different brain states is the status of locomotion**. We repeated the above analysis incorporating **only frames in which the animal is locomoting**, and found that cortical networks exhibit increased decorrelation, which results in a smaller number of statistically significant edges, degree of connectivity, and clustering coefficient, compared to equal durations of a randomly-selected interval of quiet wakefulness, where mice were not locomoting. However the differences are not statistically significant.

Fig. 34 compares various functional connectivity metrics estimated from the entire one-hour resting-state recording to those derived from resting-state frames *excluding periods of locomotion*, while Fig. 35 focuses on the period of locomotion and compares it with a randomly-selected period of equal size, under resting-state without any locomotion.

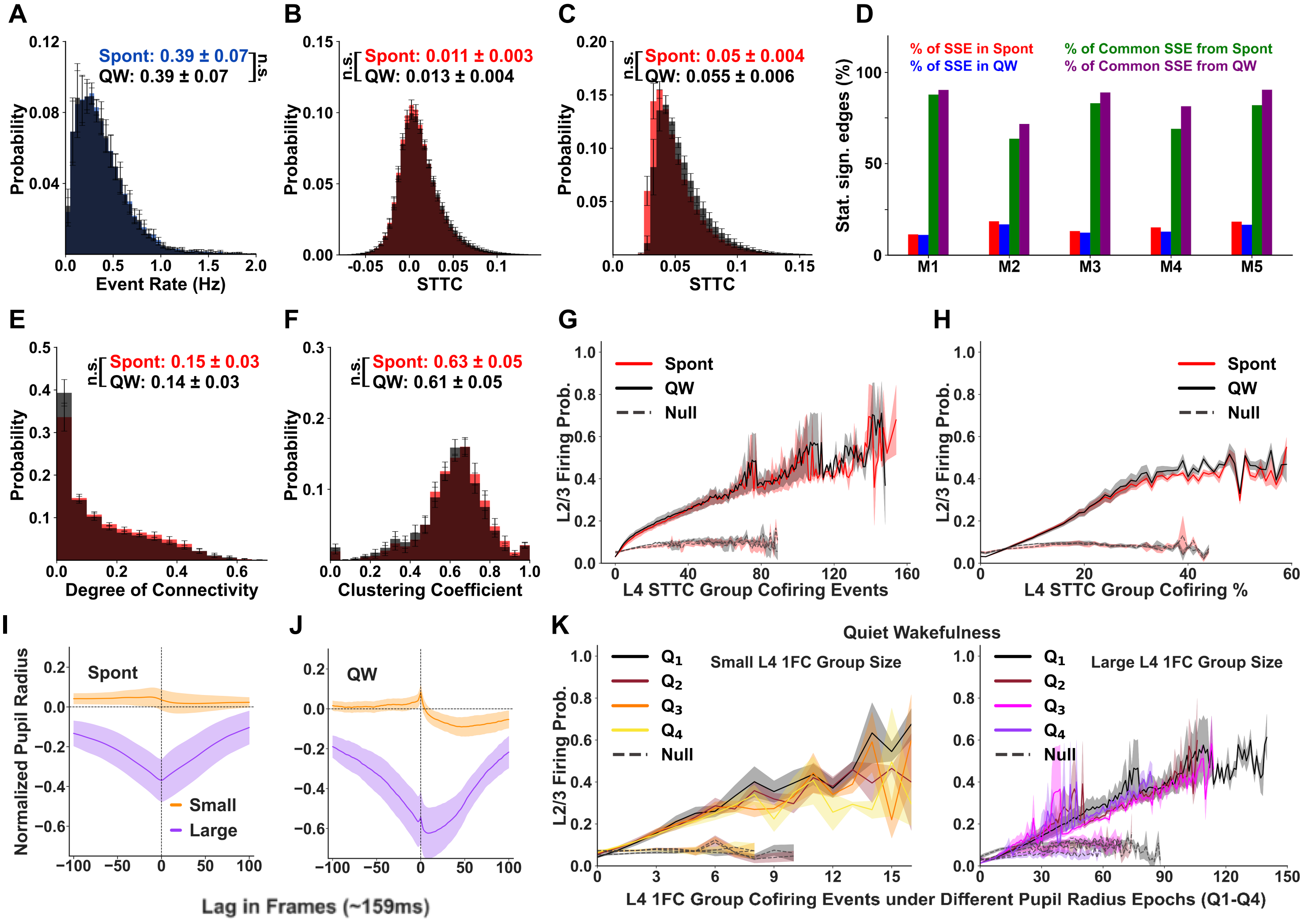


**Supplementary Figure 34: The *entire* resting-state/spontaneous period (Spont) and the period of quiet wakefulness (QW), where the mice are *not* moving, have similar properties: A.** Histogram of the event rates of the L2/3 neurons in the entire period (Spont, blue) and in the period of quiet wakefulness (QW, black). **B.** The distribution of the STTC weights of *all functional connections* from L4 to L2/3 in the entire period (Spont, red) and in the period of quiet wakefulness (QW, black). **C.** The distribution of the STTC weight of *all statistically significant (z-score>4) functional connections* from L4 to L2/3 in the entire spontaneous period (red) and in the period of quiet wakefulness (black). **D.** The percentage of statistically significant edges ("SSE") from L4 to L2/3 is similar in both periods (red, blue) and the percentage of the common ones is around 80% on average (green, purple). **E.** Histogram of the normalized degree of connectivity from L4 to L2/3 in the entire spontaneous period (red) and in the period of quiet wakefulness (black). **F.** Histogram of the clustering coefficient from L4 to L2/3 in the entire spontaneous period (red) and in the period of quiet wakefulness (black). **G.** The average probability of firing of L2/3 neurons as a function of the number of co-firing events in their L4-1FC groups, at the same imaging frame, for the entire spontaneous period (red) and the quiet wakefulness period (black). The L4-1FC groups are formed separately at each period. **H.** As in (G), except the x-axis is expressed as the percentage of neurons of the L4-1FC group that cofire. **I.** ETA on pupil radius of L2/3 recipient neurons that have small (first quartile; orange line) vs. large (fourth quartile; purple line) L4-1FC groups, respectively, under the entire spontaneous period. Note that L2/3 neurons with small L4-1FC groups tend to fire on average with a weak, if any, dependence on pupil size, while L2/3 neurons with large L4-1FC groups tend to fire when pupil size is very low. **J.** As in (I), but during the quiet wakefulness period. The L2/3 neurons with small and large L4 1FC groups exhibit similar behavior as in the entire spontaneous period, with the latter firing at an even lower pupil size. **K.** L2/3 neuron event firing probability as a function of the number of cofiring events in their L4-1FC groups, during epochs of different pupillary sizes (Q1-Q4) in the quiet wakefulness period. Note that the response is similar for all epochs irrespective of pupillary size for L2/3 neurons with small L4-1FC groups (left) and large L4-1FC groups (right), while for the latter as the pupillary size increases the observed cofiring events drops. Similar results have been observed under the entire spontaneous period (Suppl. Fig. 6.6B). The statistics in each plot indicate the mean ± standard deviation of the sample means across mice, while the shading corresponds to the standard error of the mean error bars across mice (n=5); Error bars throughout correspond to SEM across mice (n=5); "n.s.": non statistically significant. The highest p-value obtained from the permutation of means, the Welch's t-test, and the ANOVA F-test is considered for the level-of-significance (see Methods for more details).

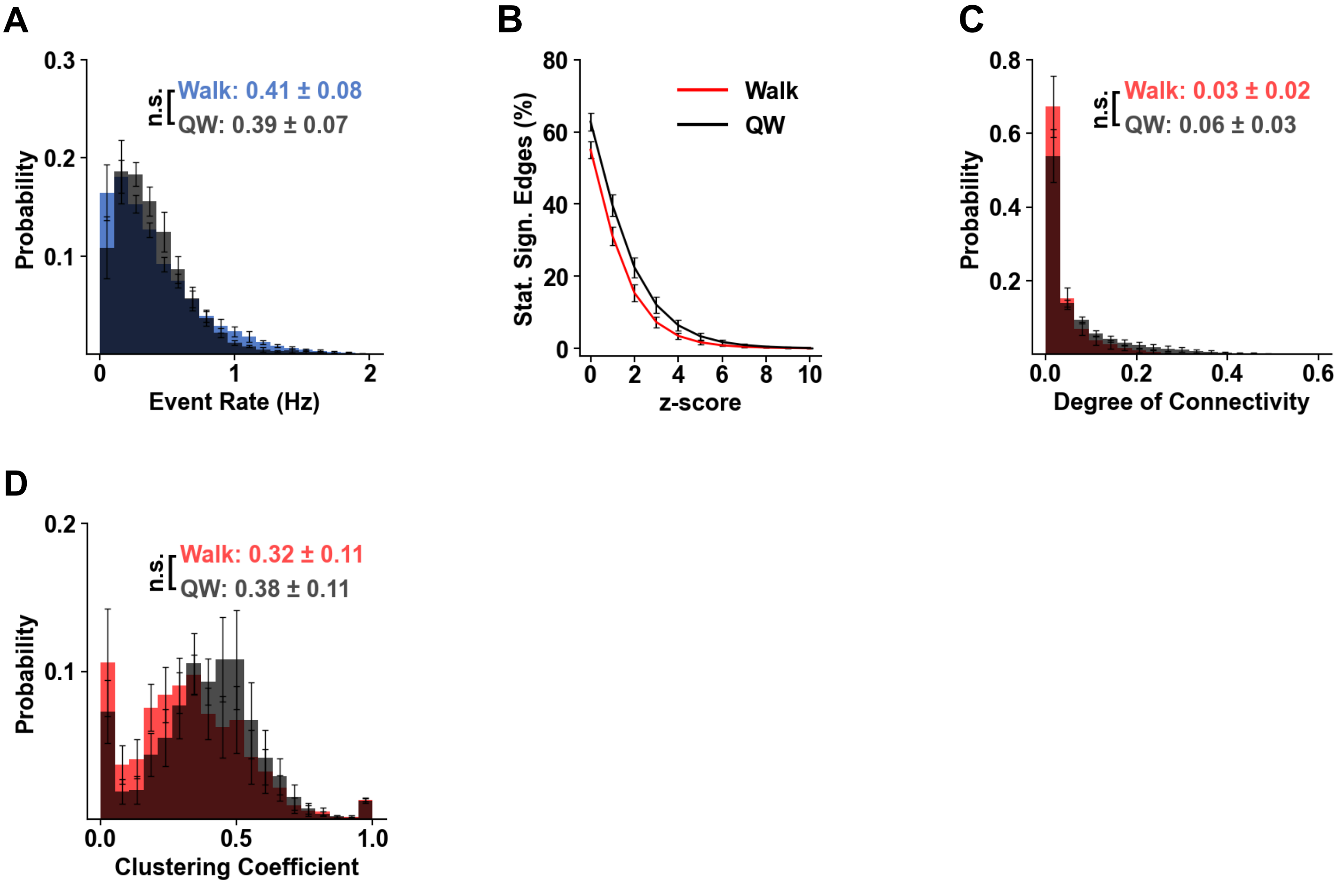


**Supplementary Figure 35. Locomotion modestly reduces L4—L2/3 functional coupling.** Walk epochs were compared with randomly selected, duration-matched quiet wakefulness (QW; resting-state, no movement) epochs from the same spontaneous recordings. L4-1FC groups are formed separately at each period. **(A)** L2/3 event-rate distributions during Walk (blue) and QW (black); mean event rates did not differ significantly across mice, though the firing rate distribution had a longer tail in the walking periods. **(B)** Fraction of significant L4—L2/3 connections versus z-score threshold during Walk (red) and QW (black) periods; QW shows a higher fraction of significant connections. **(C)** Mean normalized L4—L2/3 degree of connectivity is lower during Walk vs QW periods, but the trend does not reach significance. **(D)** L4—L2/3 clustering coefficient is lower during Walk than QW, but the trend does not reach significance. Error bars and shaded bands indicate SEM across mice (n = 5). "n.s.", not significant; significance assessed by permutation of means, Welch's t test, and ANOVA F test (largest p value used; see Methods).

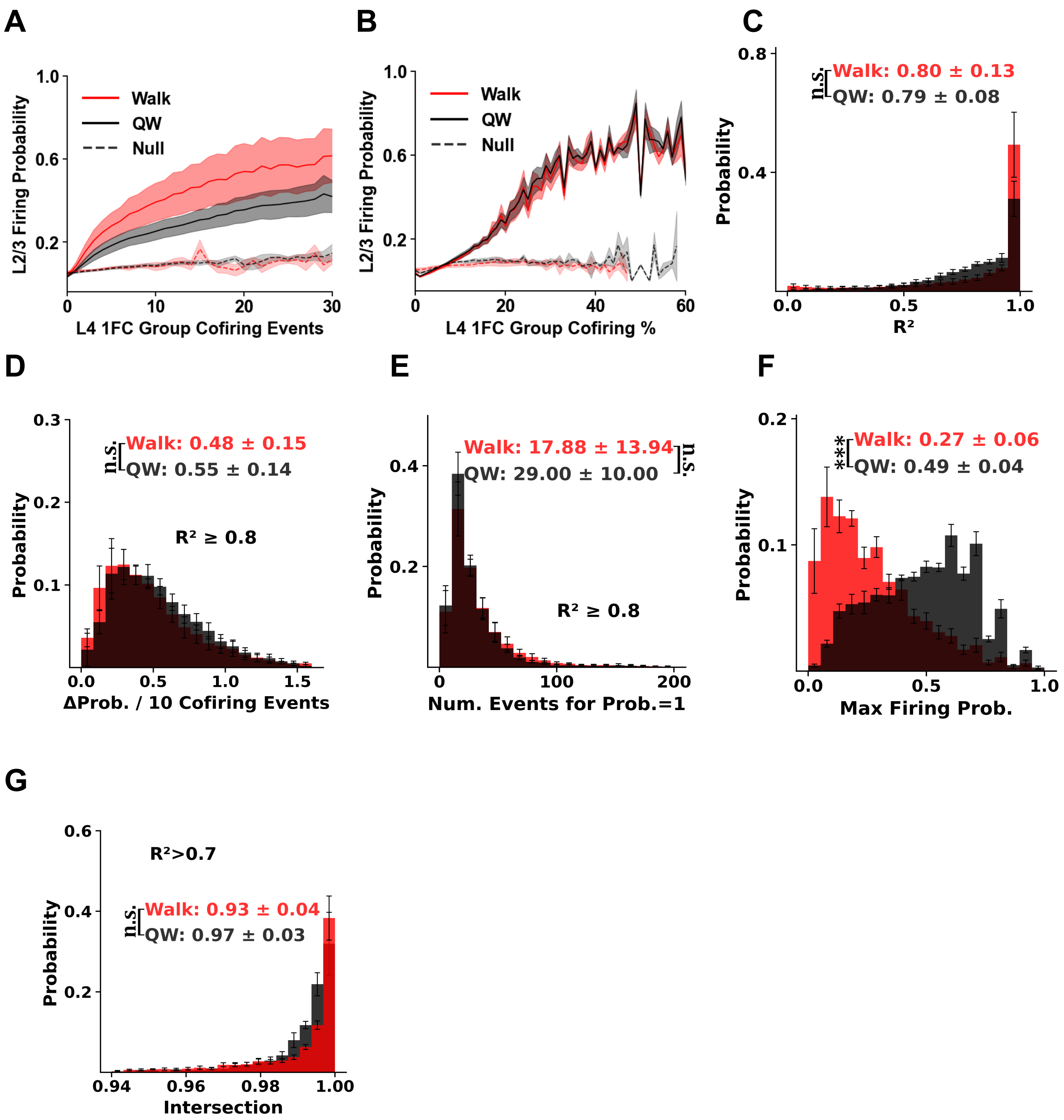


**Supplementary Figure 36. L4—L2/3 (ReLU function) response properties during locomotion versus quiet wakefulness.** Locomotion (walk) epochs were compared with duration-matched, randomly selected quiet wakefulness (QW) epochs from the same spontaneous recordings (resting-state). L4-1FC groups were defined separately within each state. **(A)** Mean L2/3 firing probability as a function of the number of within-frame co-firing events in the corresponding L4-1FC group (groups defined separately for Walk and QW); Walk exhibits a steeper response with fewer maximal co-firing events. **(B)** Same as (A), with co-firing expressed as a percentage of the L4-1FC group size; here Walking vs QW periods do not differ in slope. Note that the shared fluctuations between the two periods, and more specifically the "dips" at some ratios (e.g., 50%), may be explained by the arithmetic constraints imposed by normalization. Specifically, some percentages occur more often than others because they correspond to more common ratios. For example, any group with an even number of neurons can yield 50% cofiring; in a two-neuron group, this corresponds to just one neuron firing. By contrast, a value like 51% is far less frequent, requires larger group sizes (e.g., multiples of 51), and larger absolute number of neurons to fire together. This larger input increases the probability of firing. **(C)** Distribution of weighted $R^2$ values for ReLU fits (weighted by the standard error of each measurement). **(D–G)** Fit-derived response metrics computed from well-fit neurons ($R^2 \geq 0.8$): **(D)** slope (Δ firing probability per frame for every +10 co-firing events); The slope of the linear part of the ReLU fit; **(E)** extrapolated co-firing count at which firing probability reaches 1; **(F)** maximum actual firing probability, and **(G)** CDF value of the co-firing level at the onset ("knee") of the steep response regime. Error bars indicate SEM across mice (n = 5).

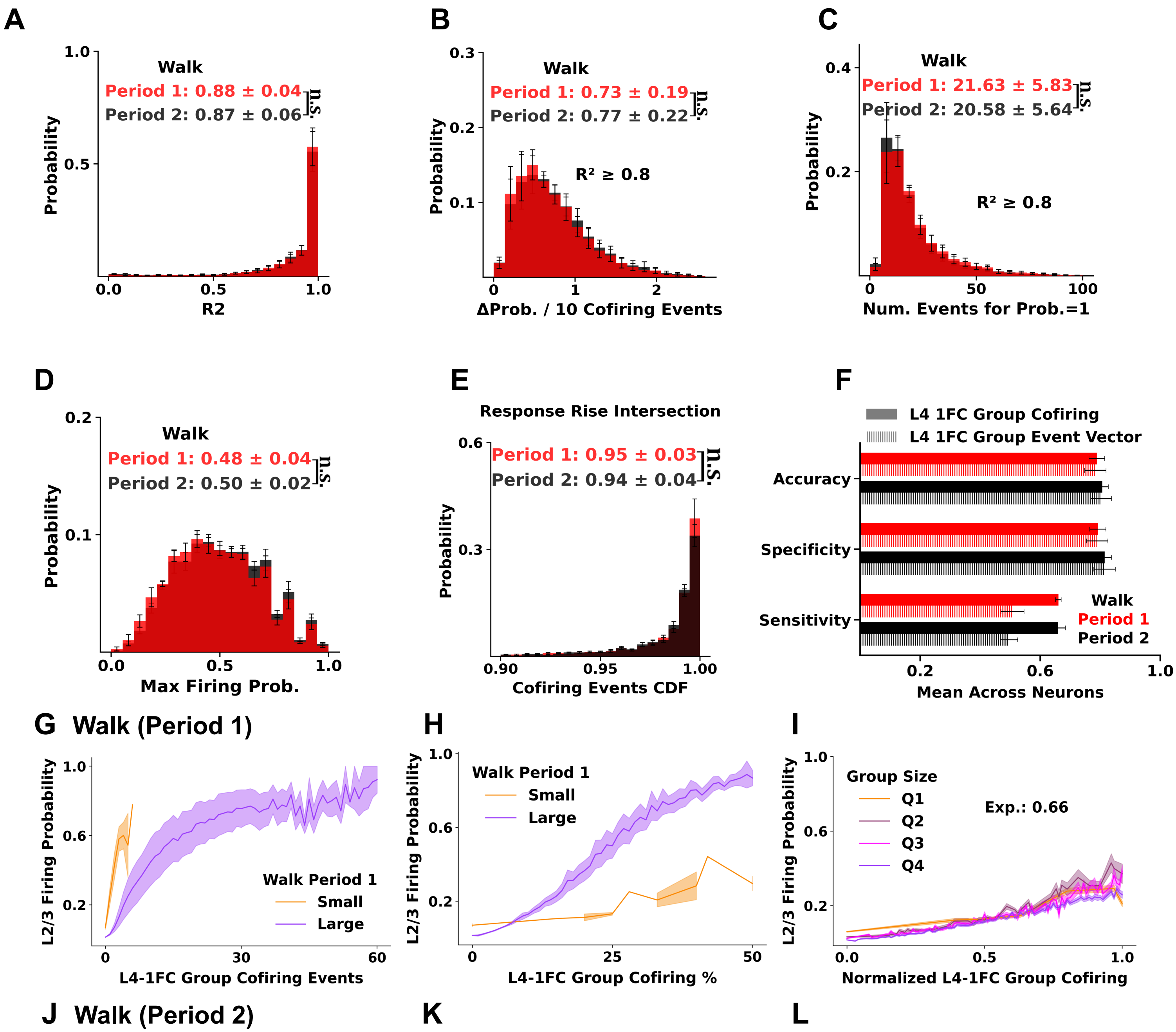

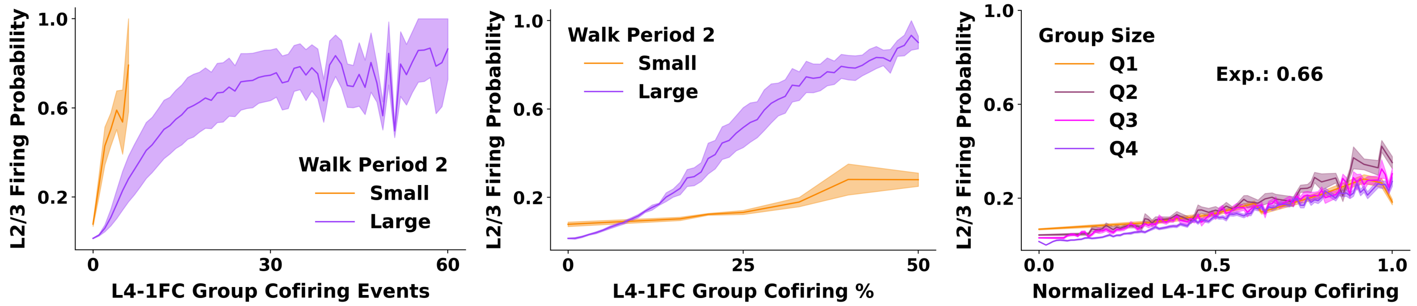


**Supplementary Figure 37. Response function stability across different periods with locomotion.** Locomotion frames were split into first (red) and second (black) halves; L4-1FC groups were computed separately within each half. **(A)** Distribution of weighted $R^2$ for ReLU fits (weights from pointwise SE). **(B–E)** Fit-derived response metrics from well-fit neurons ($R^2 \geq 0.8$): **(B)** slope (Δ firing probability of L2/3 neurons per frame for each +10 L4-1FC group co-firing events), **(C)** extrapolated co-firing count at firing probability = 1, **(D)** maximum firing probability, and **(E)** CDF value of the co-firing level at response onset ("knee"). **(F)** Linear SVM performance (sensitivity, specificity, accuracy) predicting L2/3 firing in the first (red) and second (black) half using either total L4-1FC co-firing count (solid) or the identity-resolved binary firing vector of L4-1FC members (hatched); groups defined from their respective periods. **(G–I)** L2/3 firing probability versus L4-1FC co-firing for first-half locomotion: **(G)** stratified by L4-1FC group size quartiles (smallest, orange; largest, purple), **(H)** expressed as percent of group co-firing, and **(I)** normalized co-firing (example mouse) using the exponent shown. Shading indicates SEM across mice (n = 5). **(J–L)** Same analyses as (G–I), computed for the second half of locomotion. The legends report the mean ± SD across the 5 mice. Error bars in histograms and in bar plots indicate SEM across mice (n = 5).

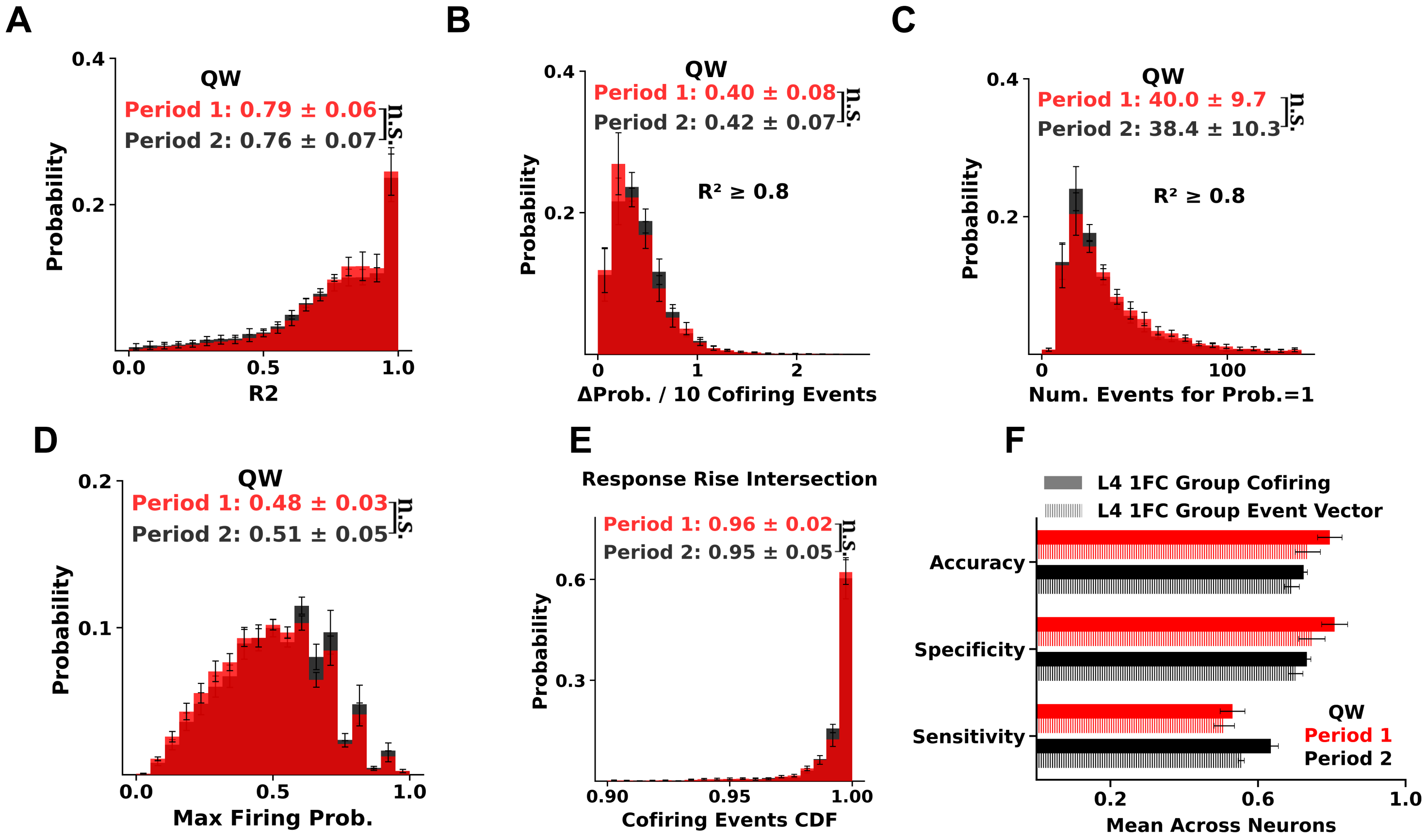


**G** QW (Period 1)

**H**

**I**

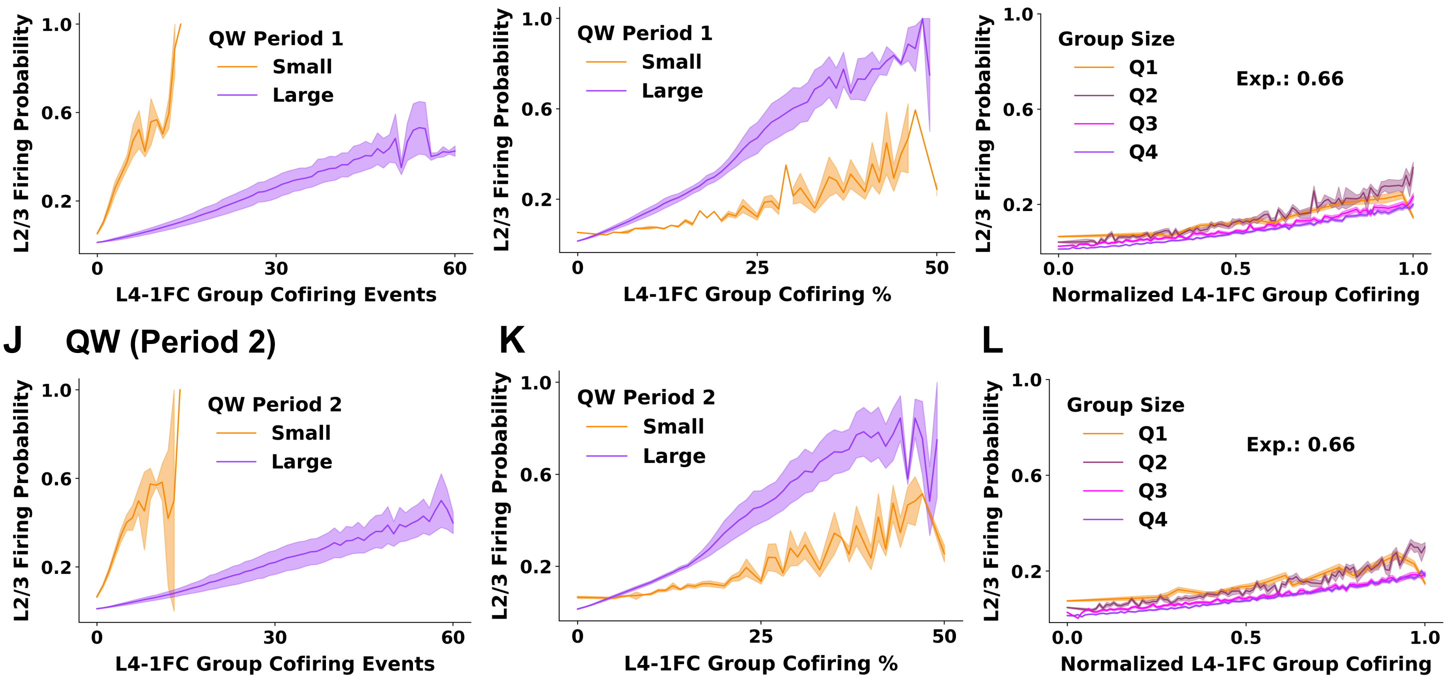


**Supplementary Figure 38. Response function stability across different quiet wakefulness periods.** Quiet wakefulness (QW; resting-state, no movement) frames were split into first (red) and second (black) halves; L4-1FC groups were computed separately within each half. **(A)** Distribution of weighted $R^2$ for ReLU fits (weights from pointwise SE). **(B–E)** Fit-derived response metrics from well-fit neurons ($R^2 \geq 0.8$): **(B)** slope (Δ firing probability per +10 co-firing events per frame), **(C)** extrapolated co-firing count at firing probability = 1, **(D)** maximum firing probability, and (E) CDF value of the co-firing level at response onset ("knee"). **(F)** Linear SVM performance (sensitivity, specificity, accuracy) predicting L2/3 firing in the first (red) and second (black) half using either total L4-1FC co-firing count (red) or the identity-resolved binary firing vector of L4-1FC members (black); groups defined from their respective periods. **(G–I)** L2/3 firing probability versus L4-1FC co-firing for first-half QW: **(G)** stratified by L4-1FC group size quartiles (smallest, orange; largest, purple), **(H)** expressed as percent of group co-firing, and **(I)** normalized co-firing (example mouse) using the exponent shown. Shading indicates SEM across mice (n = 5). **(J–L)** Same analyses as (G–I), computed for the second half of QW. Error bars indicate SEM across mice (n = 5).

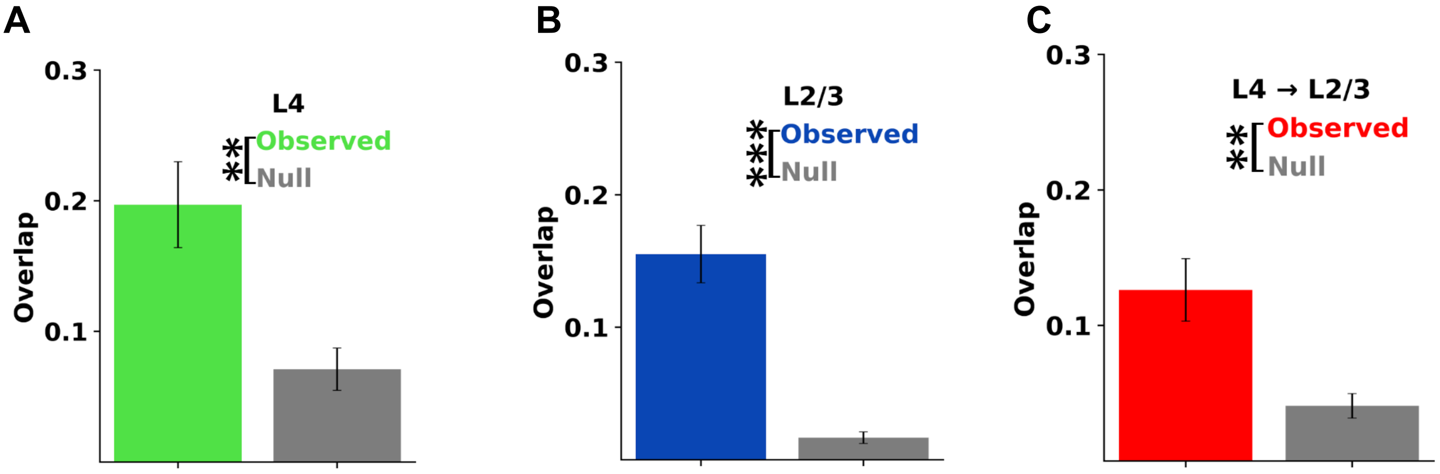

**Supplementary Figure 39: 1FC groups formed during walking and quiet wakefulness periods exhibit statistically significant overlap. A.** Number of common neurons between the L4-1FC groups of L4 neurons during walk and quiet wakefulness, divided by the square root of the factor of the two group sizes. The quiet wakefulness frames were sampled 10 times, ensuring that the number of frames is the same as in the walking period. The mean across the 10 iterations is reported for each mouse. The null is computed by selecting random neurons for the walk L4-1FC groups, ensuring that the group size doesn't change and that neurons in the observed group are excluded. **B.** Same as A, but for the L2/3-1FC groups of L2/3 neurons. **C.** Same as A, but for the L4-1FC groups of L2/3 neurons.

### 10. Functional Connectivity under Stimulus Presentation

The analysis reported so far was in the absence of visual stimulus, at resting-state, which is the focus of this work. Naturally, the presence of stimulus introduces signal correlations that impact functional-connectivity. Note that it is beyond the scope of this section to analyze fully the relation of signal correlations to resting-state (noise) correlations. Rather we focus on exploring whether the basic observations we made above remain valid in the signal correlation context.

#### Firing rate in resting state vs under stimulus presentation

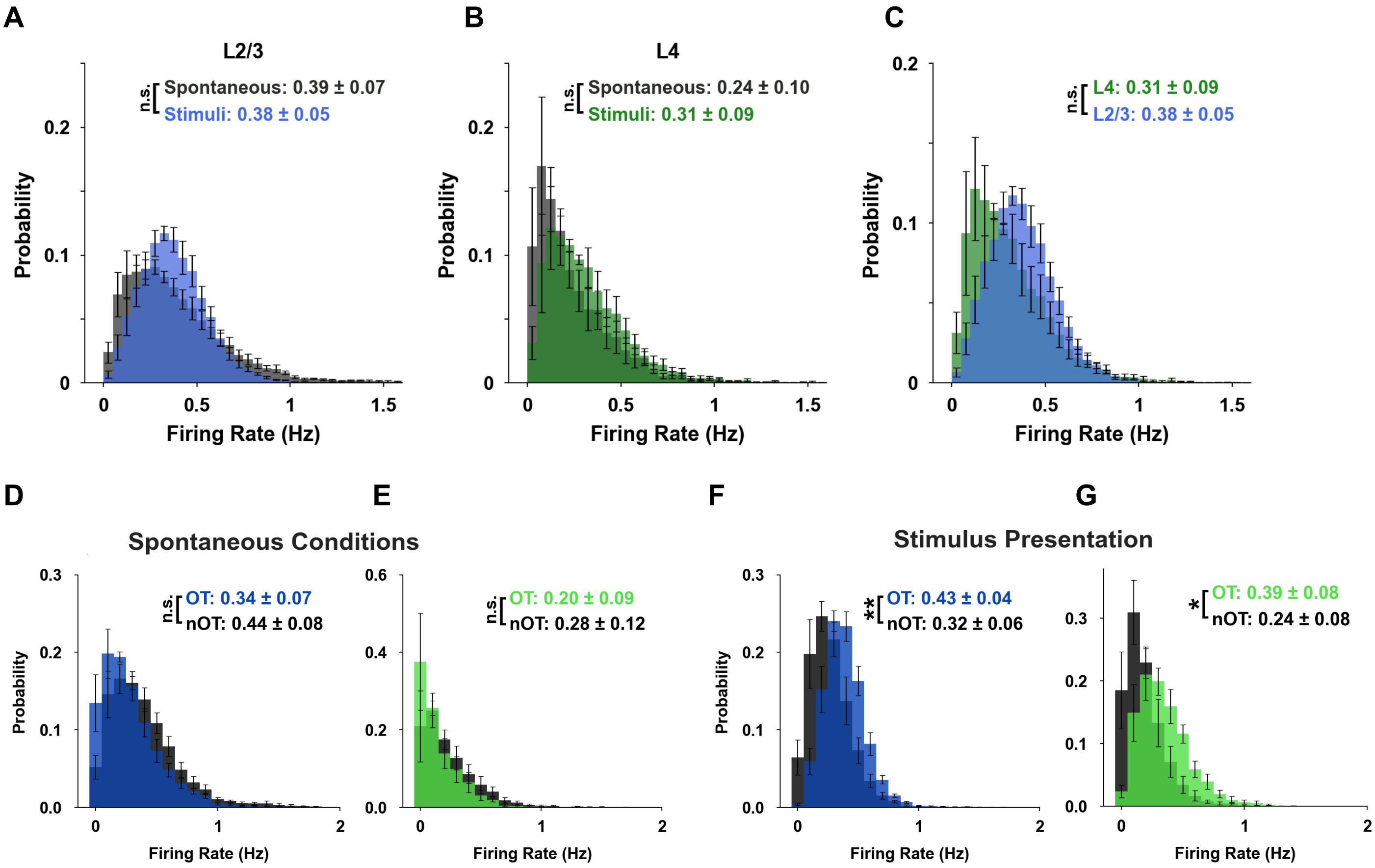

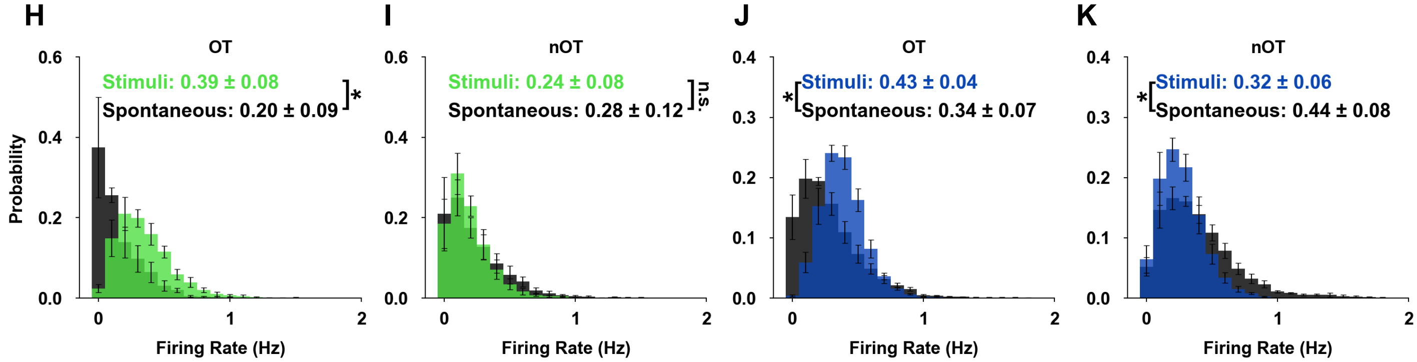


**Supplementary Figure 40. Pyramidal-neuron event-rate distributions: spontaneous vs. stimulus-evoked.** For each mouse, event-rate histograms were computed as the fraction of neurons per bin within each layer, then averaged across animals (n = 5; error bars, SEM across mice). **(A, B)** L2/3 (A) and L4 **(B)** event-rate distributions during spontaneous activity versus stimulus ("Monet", see methods) presentation. **(C)** L2/3 versus L4 event-rate distributions during stimulus presentation. **(D,E)** Event rates of orientation-tuned versus non–orientation-tuned neurons during spontaneous activity in L2/3 (D) and L4 (E). **(F, G)** Same as (D, E) during stimulus presentation. **(H,I)** L4 tuned (H) and non-tuned (I) neurons: spontaneous versus stimulus-evoked distributions. **(J, K)** Same as (H,I) for L2/3. Stimulus presentation shifts event-rate distributions rightward and alters tail structure; during stimulation, tuned neurons exhibit higher event rates than non-tuned neurons, whereas tuning has little effect during spontaneous activity. Insets show mean ± SD of mouse means across mice for each layer. Significance: permutation of means, Welch's $t$ test, and ANOVA $F$ test (largest $P$ used); $P < 0.05$ (*), $P < 0.01$ (**), n.s., not significant.

#### STTC weights, degree of connectivity, clustering coefficient, and length of functional connections

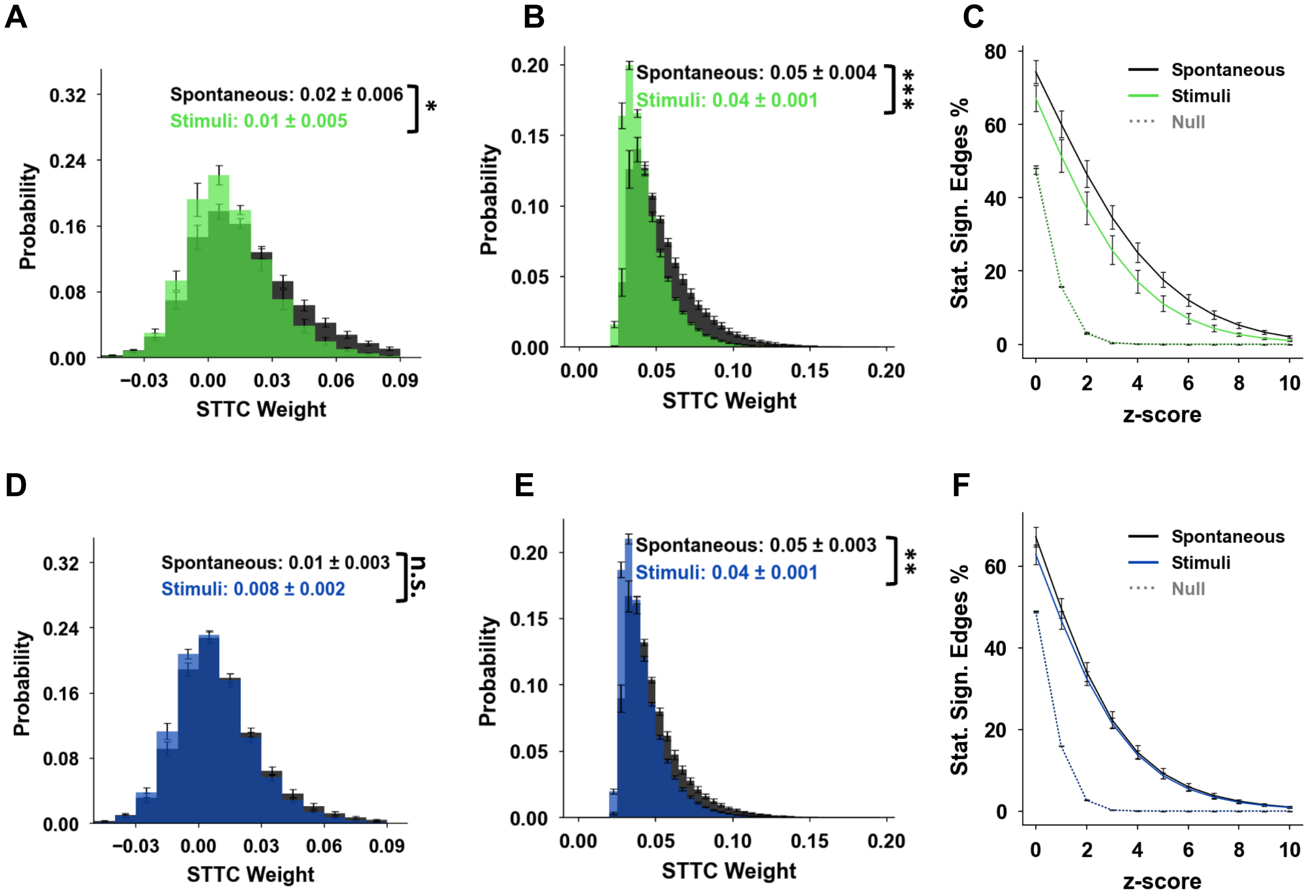

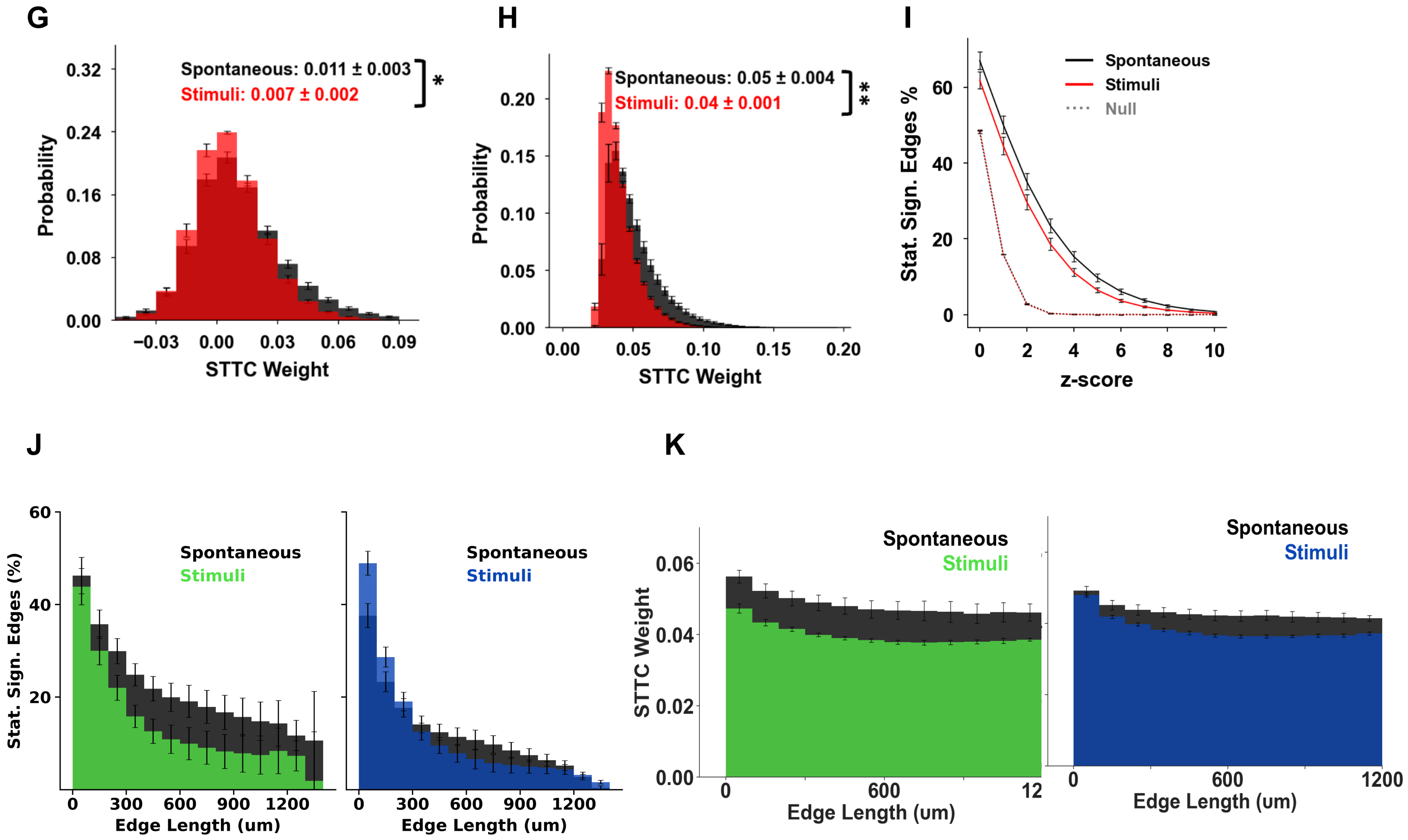


**Supplementary Figure 41 Pairwise correlations under stimulus presentation vs resting-state. A.** Histograms of STTC values between pairs of L4 neurons under stimulus presentation (green) vs. under spontaneous (resting-state) conditions (black); **B.** As in A, but focusing on *statistically significant connections* (z-score >4). **C.** Percentage of statistically significant functional connections between L4 neurons as a function of the z-score threshold. Dotted lines: corresponding null distributions obtained by random circular shifting. **D, E, F.** As in A, B and C, for the functional connectivity between L2/3 neuronal pairs, respectively. **G, H, I.** As in A, B and C, for the functional connectivity between L4 → L2/3 neuronal pairs, respectively. **J-K) Pairwise functional connectivity decays with intersomatic distance.** The prevalence of significant positive correlations decreases sharply with distance, whereas the strength of surviving correlations decreases much more weakly. Stimulus-evoked activity (blue/green histogram) exhibits a faster, larger, distance-dependent falloff than spontaneous activity (black). **J.** Fraction of positive STTC edges exceeding threshold (z-score > 4) as a function of intersomatic distance (100-μm bins). For each bin, we computed per-neuron proportions of significant pairs among all possible within-layer pairs at that distance, then averaged across mice (n = 5). Significant long-range correlations extend to ~1 mm and are more prominent in L4; the 0–100 to 100–200 μm drop is steeper in L2/3 than in L4 (right-tailed two-sample *t* test, *P* = 0.001; n = 5 mice). Distances were measured in the x–y plane; correlations were computed within the layer (L2 and L3 analyzed separately and plotted together). **K.** Mean STTC weight of significant edges from (A) versus distance, showing only a modest decline despite the larger reduction in edge prevalence. Insets: Mean STTC weight ± their standard deviation across animals (n=5), for each layer. Error bars represent the standard error of the mean across animals (n=5). P-values: "*"< 0.05; "**"< 0.01; "***"< 0.001 and "n.s.": non statistically significant. The highest p-value obtained from the permutation of means, the Welch's t-test, and the ANOVA F-test is considered for the level-of-significance.

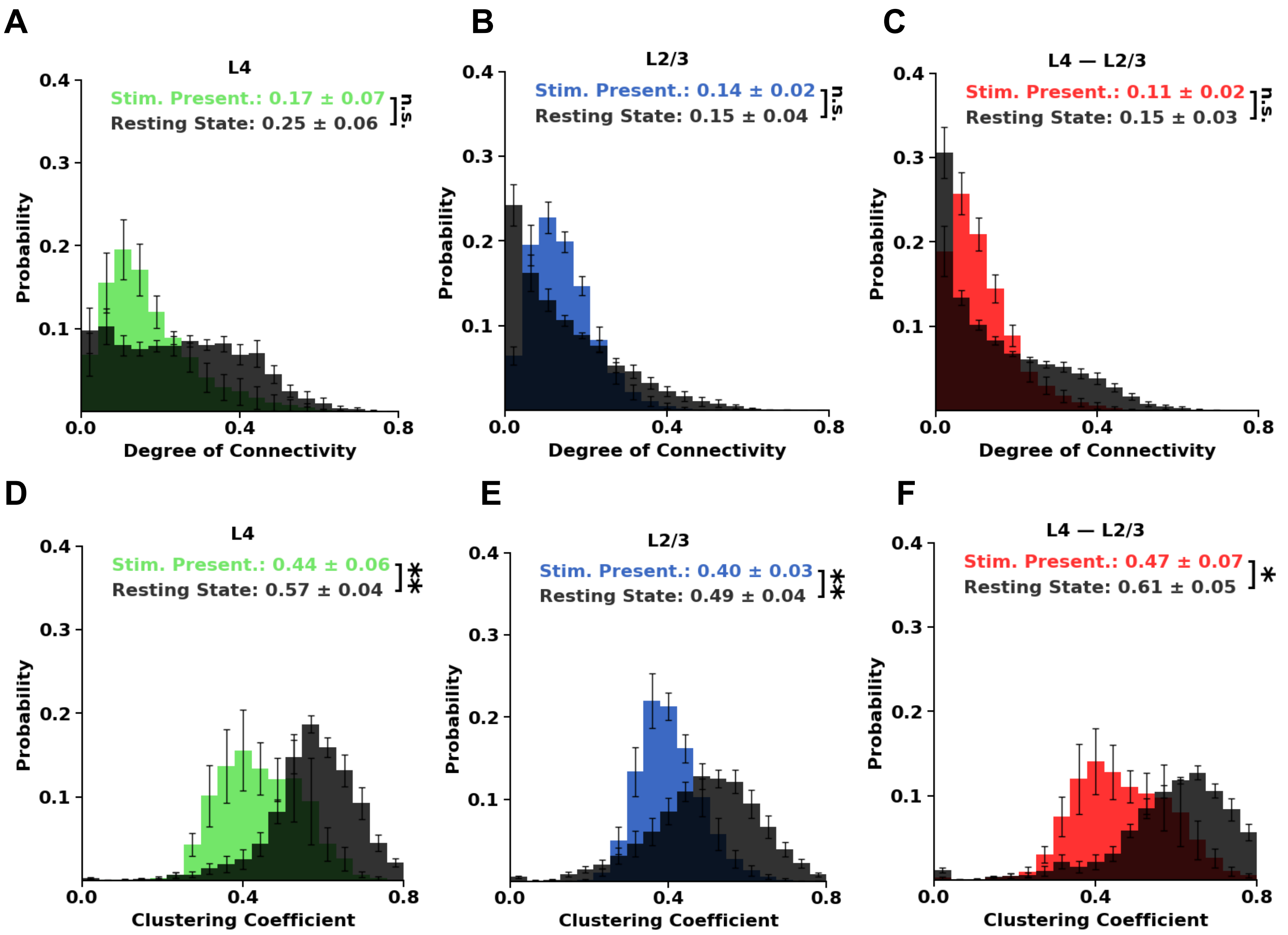


**Supplementary Fig. 42 | Stimulus presentation reduces functional degree of connectivity and clustering.** **(A–C)** Histograms of the *degree of connectivity* (DoC), expressed as a fraction of the total number of neurons in the field of view, for L4 (A), L2/3 (B), and the L4-1FC groups defined from "index" neurons in L2/3 (C), computed during *spontaneous* (black) versus *stimulus-driven* (green, blue, red) conditions. Only edges with statistically significant positive STTC (z-score >4) were included. **(D–F)** Corresponding distributions of the *clustering coefficient* for L4 neurons (D), L2/3 neurons (E), and L2/3-index neurons, considering their L4-1FC neighbor neurons (F) i.e., neurons in L4 that were significantly functionally connected to that L2/3 neuron. Both within and across layers, stimulus presentation is associated with reduced DoC and clustering compared with spontaneous activity, indicating a sparser functional organization during stimulus presentation. Insets show mean ± SD across animals (n=5). Error bars denote SEM across animals (n=5). P-values:; "**"< 0.01; and "n.s.": non-statistically significant (maximum p-value across permutation test, Welch's t-test, and one-way ANOVA).

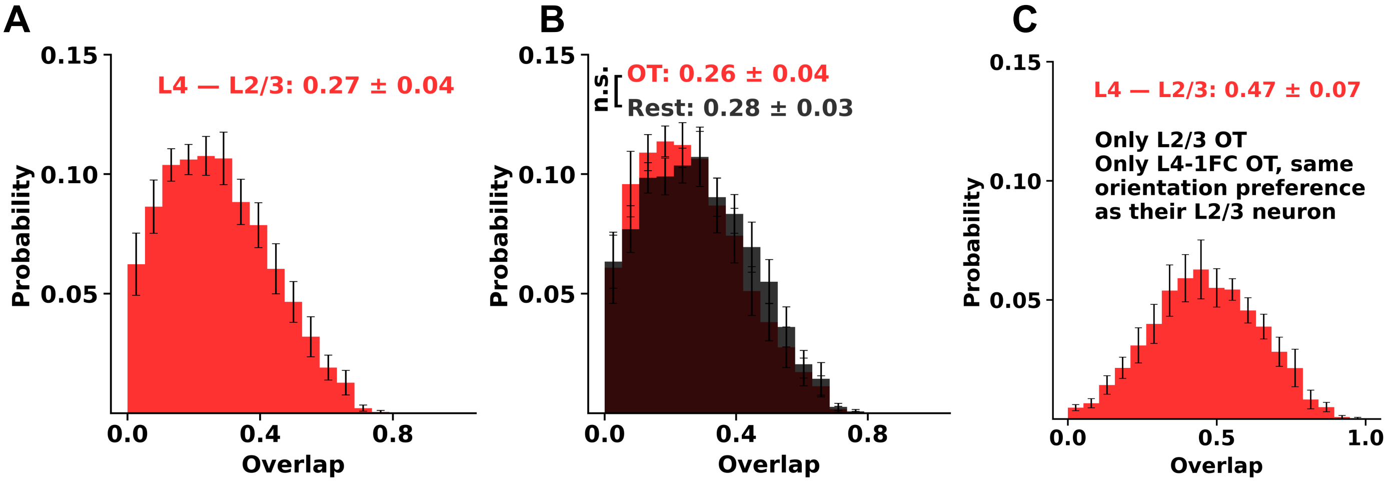


**Supplementary Figure 43. Distributions of L4-1FC group overlaps identified under stimulus presentation versus spontaneous conditions. (A)** Distribution of overlap between L4-1FC groups of a L2/3 neuron defined during resting-state (spontaneous condition; red) versus during stimulus presentation ("Monet"; black). Overlap was quantified as |Gspont∩Gstim|/|Gspont||Gstim|. **(B)** Same as (A), separated by L2/3 orientation-tuned (OT; red) versus non–orientation-tuned (nOT; black) neurons. **(C)** For OT L2/3 neurons, the overlap computed after restricting each L4-1FC group to orientation-tuned L4 neurons whose preferred orientation matches that of the L2/3 index neuron exhibits a relatively small increase. Analyses include only pairs in which both L4-1FC groups had size ≥ 5. Significance: permutation of means, Welch's *t* test, and ANOVA *F* test (largest *P* used); n.s., not significant.

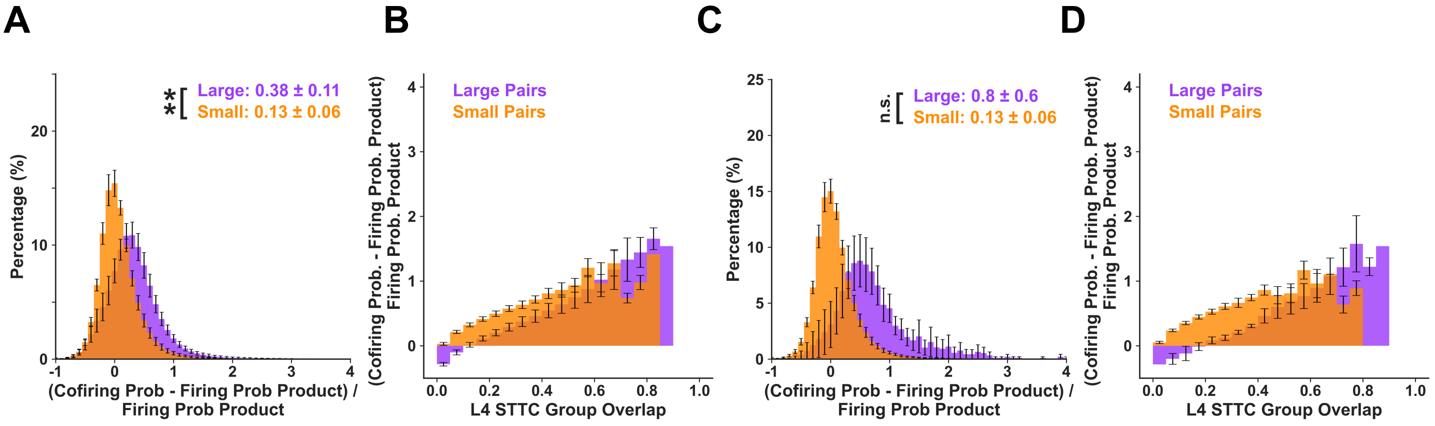


**Supplementary Figure 44 | Independence of firing of L2/3 neurons under stimulus presentation conditions for different L4-1FC group sizes. A.** Histogram of the probability that both L2/3 neurons cofire minus the product of their individual probabilities of firing, normalized by the product of the individual probability of firing. **B.** Histogram of this ratio (presented in A) under different L4 1FC group overlap sizes. The higher the overlap the less the independence in the firing of the L2/3 neurons, reinforcing the idea of collaboration between the L4-1FC ensembles. This is true for both L2/3 neurons with large L4-1FC groups as well as small ones. **C.** The figure is created under stimulus presentation, similarly to A but for a ***subset of the L2/3 reference neurons.*** The subset, for a specific quartile Q (Q1: Small, Q4: Large), is composed of the L2/3 index neurons whose group size, as defined in the stimuli-period, is within the range of group sizes of quartile Q as defined during the *spontaneous period*. **D.** As in B, but for the subset of L2/3 neurons identified in Fig. C. The error bars correspond to SEM across mice (n=5); Histogram inset reports the mean ± standard deviation of the means across mice (n=5); P-values: "*" < 0.05. The highest p-value obtained from the permutation of means, the Welch's t-test, and the ANOVA F-test is considered for the level-of-significance (see Methods for more details).

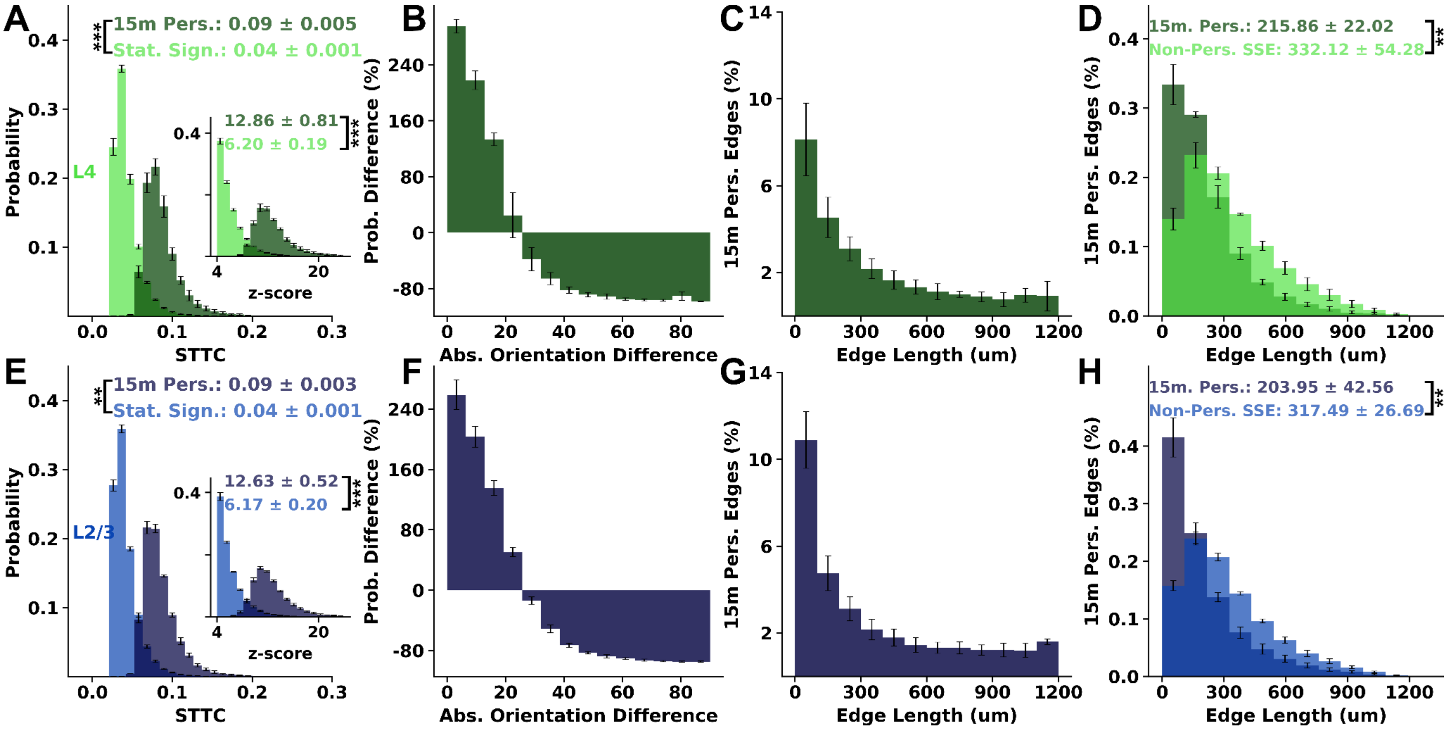


**Supplementary Figure 45 | Persistent functional edges during stimulus presentation.** *Persistent edges* were defined as STTC connections that remained statistically significant in each of four non-overlapping 15-min epochs sampled across the stimulus session. **(A)** Distributions of STTC weights (main) and z-scores (inset) computed over the full session for *persistent* intra-L4 edges (dark green) versus *all significant* intra-L4 edges (light green); persistent edges are a subset of the significant edge set. **(B)** Histogram of the orientation preference difference among L4 neuronal pairs with persistent functional correlations minus that formed by pairs with statistically significant non-persistent functional connections, normalized bin by bin by the nonpersistent distribution. Orientation difference was computed as the minimal absolute difference on the circular [0,180) space (0 ≡ 180), yielding $\Delta\theta \in [0,90]$ (see Methods). **(C)** Fraction of significant intra-L4 edges that were persistent versus intersomatic distance (in 100-µm bins), computed per index neuron as #persistent/#significant within each distance bin and then averaged across mice. **(D)** Edge-length distributions for persistent versus nonpersistent significant intra-L4 edges. **(E–H)** Same as (A–D) for intra-L2/3 edges. In both L4 and L2/3, persistent edges show *higher STTC weights/z-scores, greater tuning similarity, and shorter characteristic lengths* than other significant edges. Summary values are mean ± SD of mouse means; error bars, SEM across mice (n = 5). P-values: "**" < 0.01; "***" < 0.001. Significance was assessed by permutation of means, Welch's *t* test, and ANOVA *F* test (largest *P* reported).

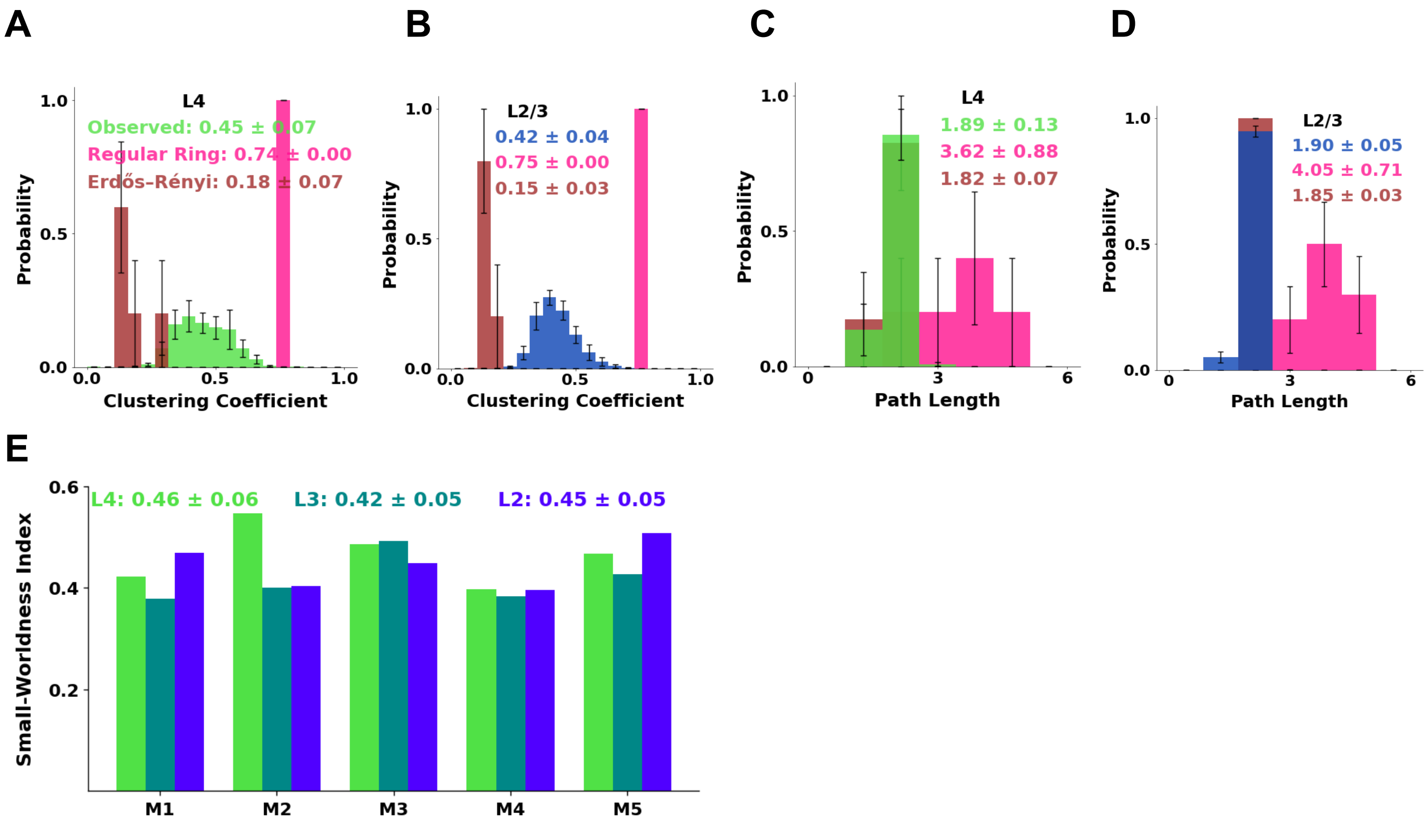


**Supplementary Figure 46 | Functional connectivity under stimulus presentation exhibits small-world characteristics.** The layer network graphs based on the pairwise statistically significant connections (with z-score >4), estimated *during stimulus presentation*, were formed and compared with their corresponding theoretical graph models, namely the Erdős–Rényi and the regular ring (see methods). **A)** Histogram of the clustering coefficient of L4 neurons (green) vs. the corresponding theoretical graphs random Erdős–Rényi (purple) and regular ring (pink), across mice (n=5). Here the clustering coefficient is estimated considering the entire set of frames without excluding the frames that the index neuron fires. **B)** Same as A, for Layer 2/3 (blue). **C-D)** As in A-B but now we histogram the shortest path length between pairs of neurons. Notice the small average path length of the observed functional connectivity, closer to the one estimated on the Erdős–Rényi graphs compared to regular ring graphs. On the other hand, the clustering coefficient distributions of the observed networks are closer to that of the regular ring graphs. Thus, the functional connectivity networks of L4 and L2/3 computed *under stimulus presentation* do manifest small-worldness (when compared with the Erdős–Rényi and regular ring, in terms of clustering coefficient and path length). **E)** The small-worldness index per mouse (M1-M5) for each layer case separately (green: L4, purple: L2, dark turquoise: L3). The error bars correspond to SEM across mice (n=5); Plot insets report the mean ± standard deviation across mice.

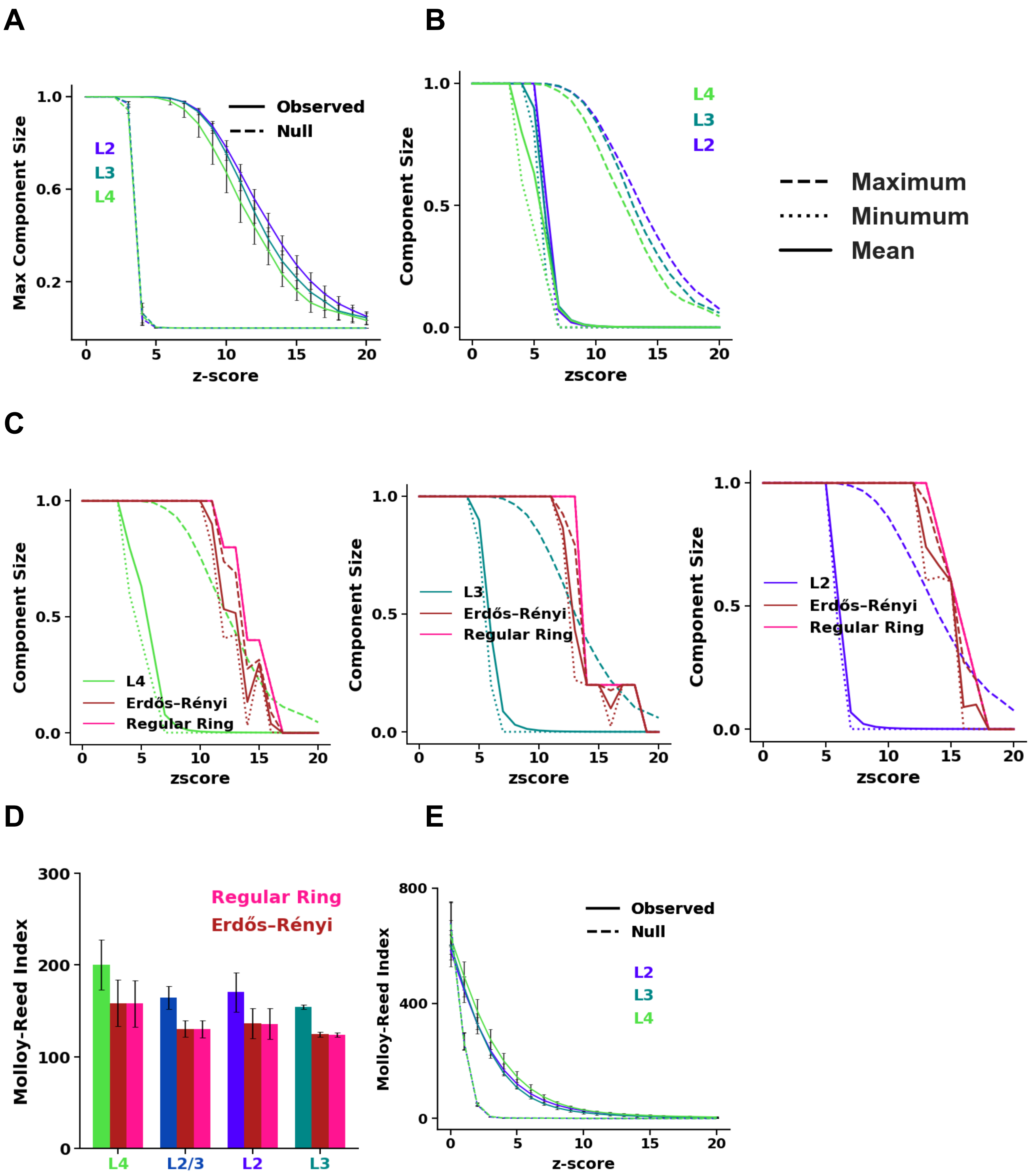


**Supplementary Fig. 47 | Functional networks in V1 remain robust during stimulus presentation. (A)** Fraction of neurons in the largest connected component versus z-score threshold for significant functional links; at $z \geq 4$, L4, L3, and L2 networks exhibit a giant component spanning nearly the entire layer and it persists at stricter thresholds. **(B) The functional connectivity architecture at each layer exhibits network robustness during stimulus presentation: up to large z-score values, the network consists of a giant component.** For each layer (namely L4, L3, L2), the size of the mean (solid), maximum (dashed), and minimum (dotted) connected component of the network, formed at each z-score threshold, is reported as a function of the z-score threshold (y-axis: fraction of the total size of the layer). The mean, minimum, and maximum component size for the observed functional network *under stimulus presentation*, versus theoretical graphs, namely *regular ring* (fuchsia) and *Erdős–Rényi*

(dark brown/grenat). **(C)** In the observed network up to z-score threshold 4, there is a single connected component. **(D)** Network robustness quantified by the Molloy–Reed index, $\langle k^2 \rangle / \langle k \rangle$ (robust if >2), where K is the degree of connectivity, computed at $z = 4$ on within-mouse size-matched subnetworks (neurons subsampled to match the smallest layer); L4 is most robust, followed by L3 and then L2, and all exceed matched regular-ring and Erdős–Rényi graphs. **(E)** Molloy–Reed index versus z-score threshold for observed (solid) and null (dashed) networks; nulls were generated by circularly shifting each neuron's event/spike train before recomputing links (see Methods), with size-matching performed at each threshold.

#### 11. On the Impact of Tuning Properties on Functional Connectivity under Stimulus Conditions

This section examines the impact of tuning properties on functional connectivity under stimulus presentation and comparatively analyzes it with the functional connectivity under resting-state. We focused on the role of the pyramidal orientation-tuned neurons in the functional connectivity architecture. Specifically, we compared the orientation-tuned neurons vs. rest in terms of firing rate, degree of connectivity, and clustering coefficient under resting state (spontaneous) vs stimulus presentation.

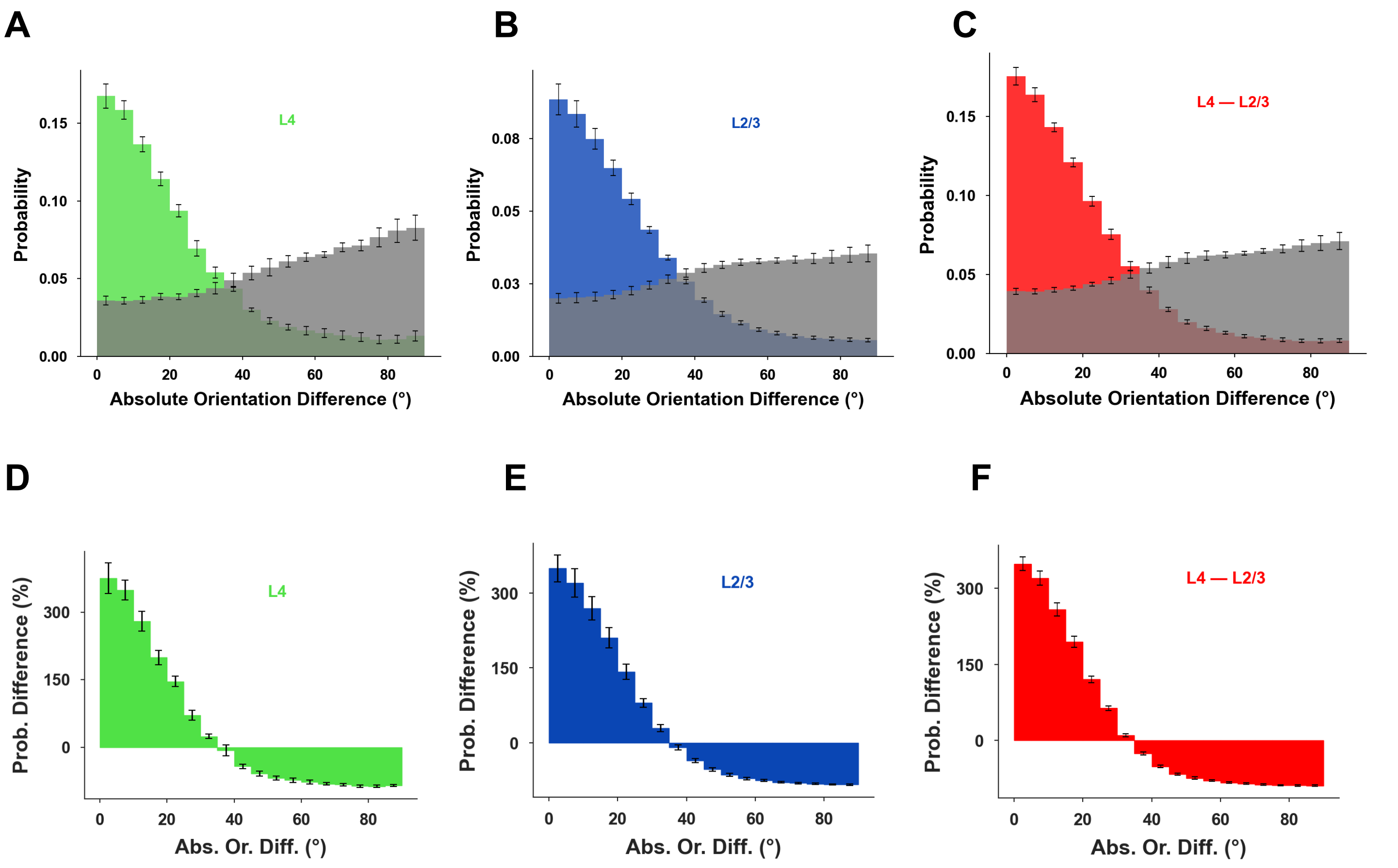


**Supplementary Fig. 48 | Stimulus-epoch functional connectivity is strongly biased by tuning similarity. (A–C)** Fraction of _positive_, statistically significant (STTC z-score >4) functional-correlation edges identified during stimulus presentation, binned by absolute orientation-preference difference for L4–L4 (A; green), L2/3–L2/3 (B; blue), and L4–L2/3(C; red) neuronal pairs. Gray bars show the corresponding null edge (z-score in [−2,2]) fraction. Orientation difference was computed as the minimal absolute difference between preferred orientations on [0,180), yielding a range of [0,90] degrees (Methods). **(D–F)** Enrichment relative to null: (significant – null) /

(null-bin probability), expressed as percentage, for L4–L4 (D; green), L2/3–L2/3 (E; blue), and L4–L2/3 (F; red). Error bars denote SEM across mice (n=5). Interestingly, a strong tuning-dependent functional connectivity structure is observed under stimulus-driven conditions, in contrast to an order of magnitude weaker bias seen in the absence of stimulus, at resting state.

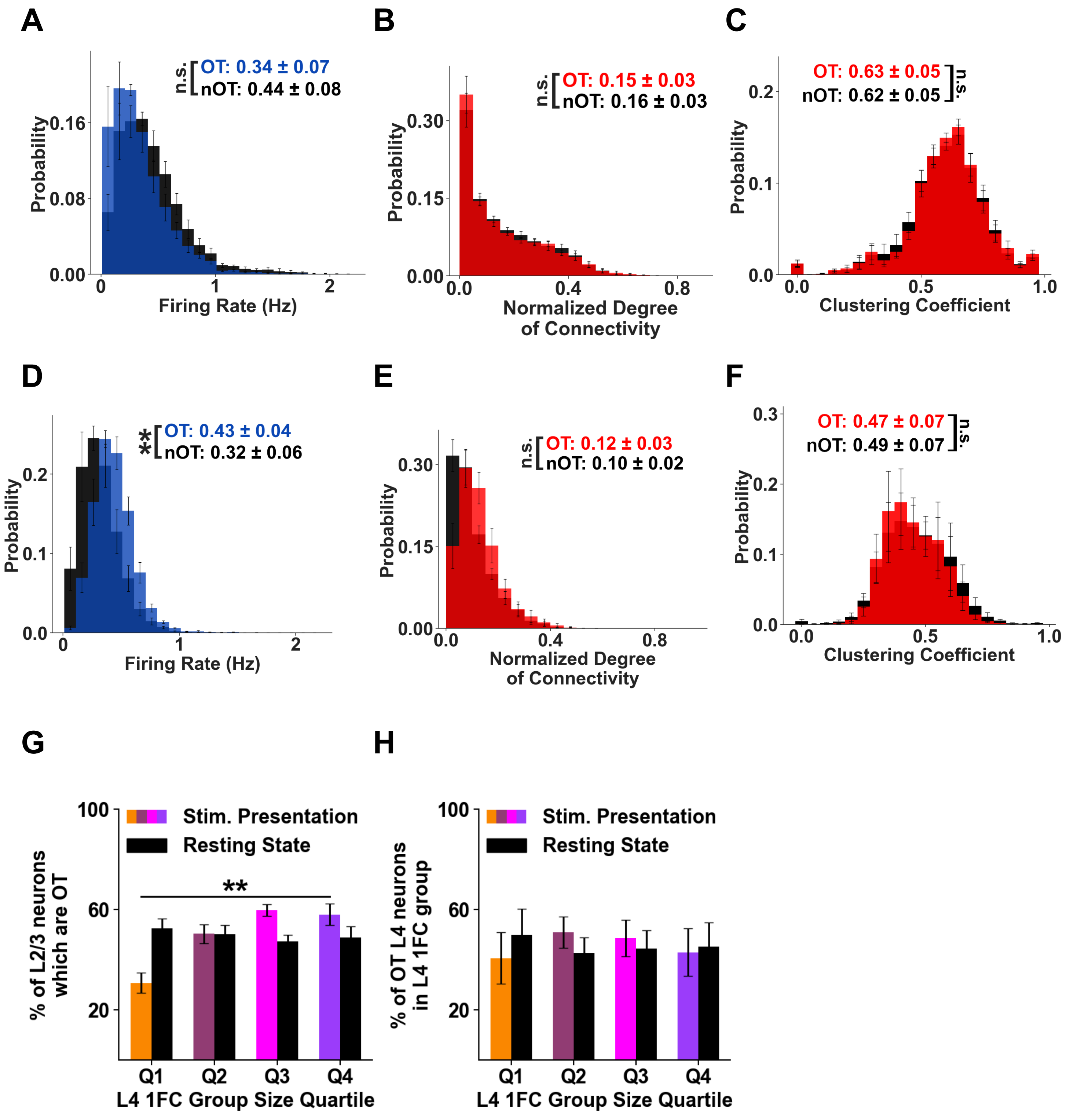


**Supplementary Fig. 49 | Orientation tuning versus functional connectivity coupling.** Orientation-tuned (OT) and non–orientation-tuned (nOT) L2/3 pyramidal neurons show comparable distributions of event rate, L4 degree of connectivity (L4-1FC size), and L4-1FC clustering under spontaneous conditions. **(A–C) Resting state:** histograms of L2/3 event rate (A), normalized L4-1FC size (B), and L4-1FC clustering coefficient (C) for OT (colored) and nOT (black) L2/3 neurons. **(D–F)** Same metrics during **stimulus presentation**. As expected, firing event rate of orientation tuned neurons is higher under stimulation conditions, but otherwise degree of connectivity and clustering coefficient distributions remain similar. **(G)** Fraction of L2/3 neurons that were OT as a function of their corresponding their L4-1FC group size quartile (Q1–Q4). Note that at resting state (rL4-1FC group size) there is no bias (black bars). In contrast, during stimulation, L2/3 neurons with larger sL4-1FC groups are more likely to be orientation tuned, as expected, since functional connectivity is now driven in part by stimulus correlations. (colored; e.g., ~58% OT in Q4 versus ~30% in Q1. This difference is statistically significant). **(H)** Fraction of OT L4 neurons within L4-1FC groups across quartiles, shown for rest (rL4-1FC groups; black) and stimulation (sL4-1FC groups; colored). Error bars denote SEM across mice (n=5). Significance: P-values: "**" < 0.01; and "n.s.": non statistically significant. The highest p-value obtained from the permutation of means, the Welch's t-test, and the ANOVA F-test is considered for the level-of-significance.

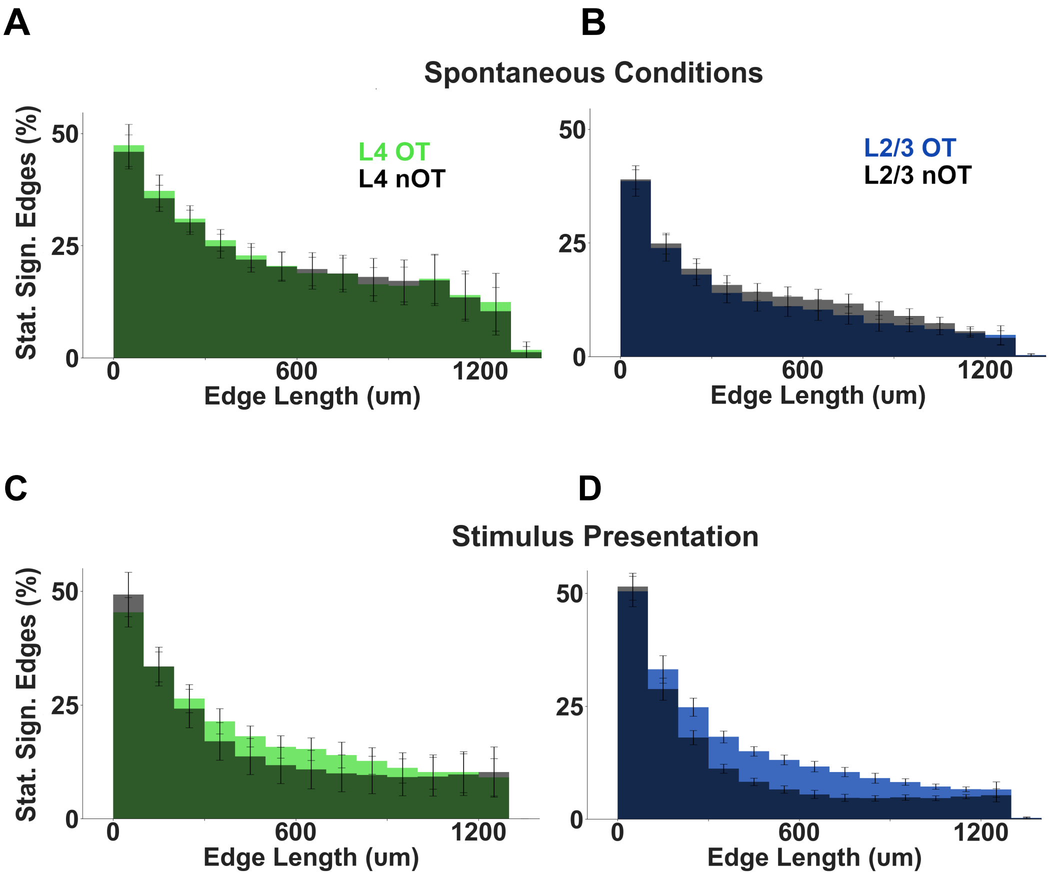


**Supplementary Figure 50 | Prominence of significant connections between OT neurons vs between nOT neurons as a function of distance. A.** Positive inter-neuronal functional correlations with z-score >4 ("edges") *between L4 orientation-tuned neurons* (light green) and between L4 neurons *that are not* orientation tuned (dark green) as a function of distance from the soma, expressed as a fraction of total possible pairwise connections between L4 orientation-tuned and non orientation-tuned neurons, respectively, at that distance plotted in bins of 100μm (light green). The histogram reflects the mean and the standard error of the mean across animals under *spontaneous conditions*. The darker histograms correspond to the same distribution but for the inter-neuronal functional correlations between L4 neurons *without* orientation tuning. **B.** As in A but for L2/3. **C-D.** As in A-B but under *stimulus presentation*. Orientation-tuned L2/3 neurons (lighter blue color, as in A-B) have a larger percentage of statistically significant edges with mid-range length, such as between 200μm-1mm. Error bars: SEM across mice (n=5).

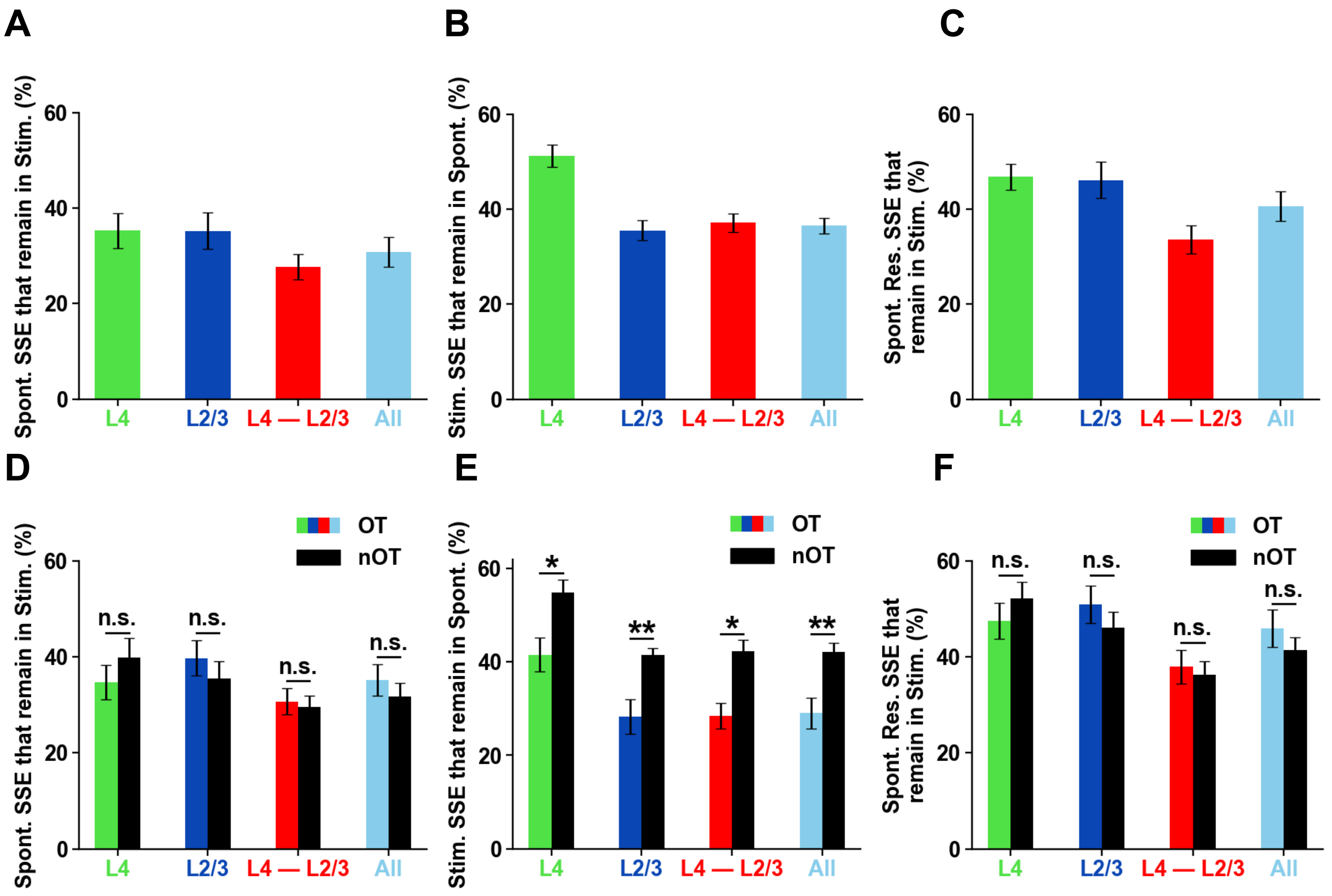


**Supplementary Fig. 51| Partial overlap of functional connectivity between spontaneous and stimulus-driven conditions. A.** Fraction of statistically significant positive functional connections (STTC z-score > 4) identified during spontaneous activity that remain significant during stimulus presentation. In L4, ~35% of resting-state edges persist under stimulation, with comparable fractions observed for intra-layer L2/3 and interlaminar L4-L2/3 connections. **B.** Reciprocal analysis showing the fraction of stimulus-defined significant edges that remain significant during spontaneous activity. In L4, ~51% of stimulus-defined edges persist at rest, whereas lower fractions are observed for L2/3(~31%) and L4-L2/3 (~34%) connections. **C.** Overlap of significant edges computed based on the residual signal, i.e. after removing the first two principal components from the spontaneous activity epochs. The subtracted components capture a substantial portion of global brain state variance (e.g., population activity and pupil size). Relative to the full resting-state signal, a larger fraction of residual-signal edges remains significant under stimulus presentation, consistent with the residual activity being enriched for connections associated with stimulus-related processing. Note that for intra-L4 connections, the difference with the values presented in A is statistically significant ($p<0.01$). **D–F.** Analyses corresponding to (A–C), restricted to connections between *orientation-tuned* (OT; color bars) neurons versus *non–orientation-tuned* (nOT; black bars) neurons. In all layers, nOT–nOT connections identified under stimulus conditions are more likely to persist under spontaneous conditions, compared to OT-OT connections (panel E). In contrast, nOT–nOT connections identified under spontaneous conditions have similar probability as OT-OT connections to persist under stimulus conditions. This suggests that nOT-nOT versus OT-OT neuronal pairs are driven differently by internal state modulations occurring during stimulus-driven vs spontaneous activity states. In all layers, residual-signal edges (panel F; edges computed in the spontaneous activity condition, after removing the first 2 principal components) show greater cross-condition persistence than edges derived from the full resting-state signal, consistent with panel (C). Error bars denote SEM across mice (n=5).

#### Composition of L4-1FC of orientation-tuned neurons

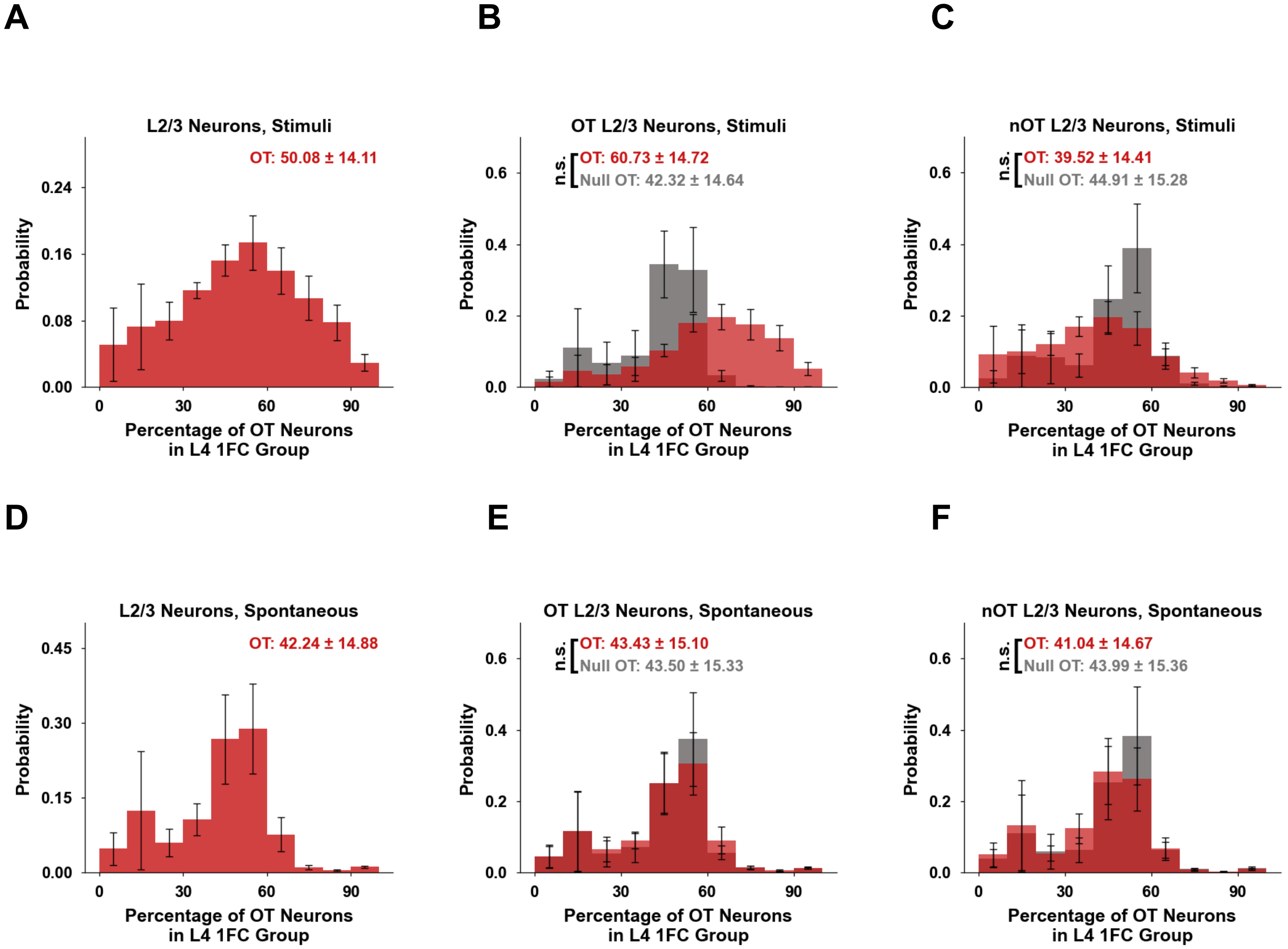


**Supplementary Figure 52 | Orientation selectivity of L4-1FC group membership under stimulation versus spontaneous conditions.** Distributions show the fraction of orientation-tuned (OT) L4 neurons within each L2/3 neuron's L4-1FC group. **(A)** Histogram of the percentage of OT neurons across all stimulus-defined sL4-1FC groups. **(B)** Histogram (red) of the percentage of OT neurons across all stimulus-defined sL4-1FC groups *that correspond to orientation tuned L2/3 neurons*. A null distribution (gray) corresponding to randomly selected size matched L4 groups of neurons outside each actual sL4-1FC group, is plotted for comparison. **(C)** As in (B) for sL4-1FC groups corresponding to non-orientation tuned (nOT) L2/3 neurons. Note that under stimulation, sL4-1FC groups corresponding to tuned (not-tuned) L2/3 neurons are more likely to contain a larger percentage of tuned (not-tuned) L4 neurons respectively. **(D–F)** Similar plots obtained under spontaneous activity conditions (rL4-1FC groups). Under resting state, rL4-1FC groups are broadly mixed with a higher percentage of nOT neurons (~58% on average) irrespective of whether they correspond to a tuned or not-tuned L2/3 neuron. This is consistent with a reduced tuning specificity of functional coupling under spontaneous conditions. Insets: mean ± SD across mouse means; error bars, SEM across mice (n = 5). Significance: permutation of means, Welch's *t* test, and ANOVA *F* test (largest *P* reported); n.s., not significant.

### 12. L2/3 Firing Prediction using the L4-1FC Cofirings of Groups Identified under Stimulus Presentation

This section comparatively analyzes the response of L2/3 neurons as a function of the number of cofirings of L4-1FC groups identified under stimulus presentation vs. resting state. We show that the rules of communication observed in resting state (e.g., ReLU function, normalization process) remain true under visual stimulation conditions, even though signal correlation architecture differs markedly from that at resting-state. This suggests that the rules of communication we observed may reflect a general property of cortical organization. Finally, we identified L4-to-L2/3 ensemble-to-ensemble transmission pathways, demonstrating superior sensitivity, precision, accuracy, and specificity compared to L4-ensemble to L2/3-neuron transmission, demonstrating that information transmission is best understood as ensemble-to-ensemble.

#### Under stimulus presentation

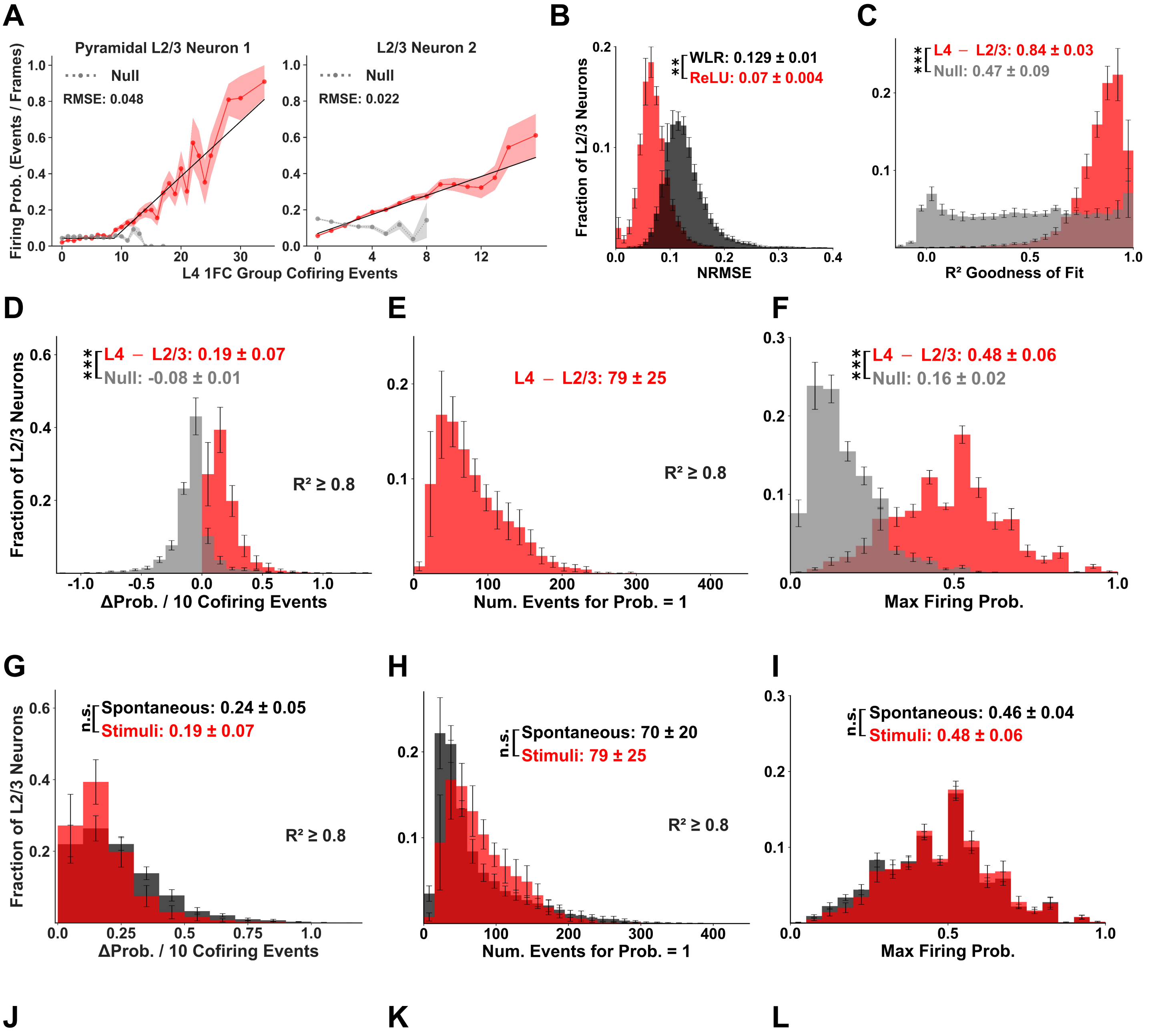

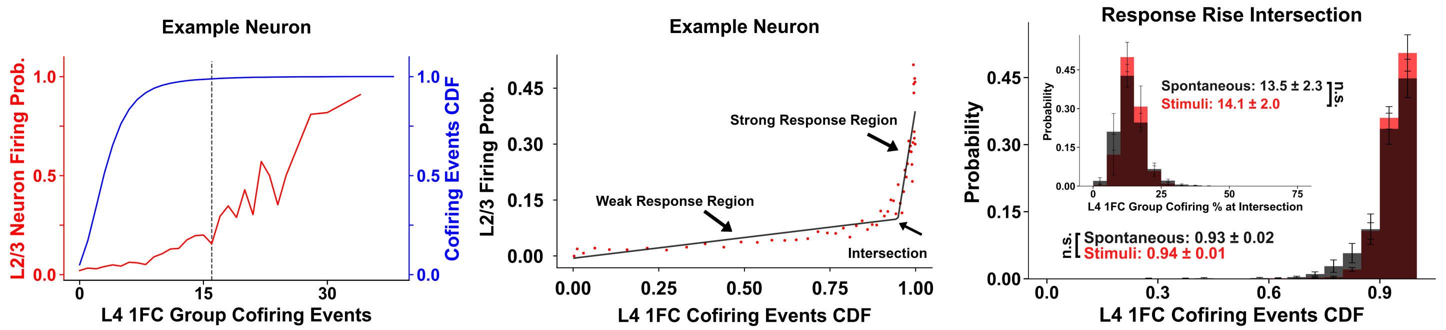


**M**

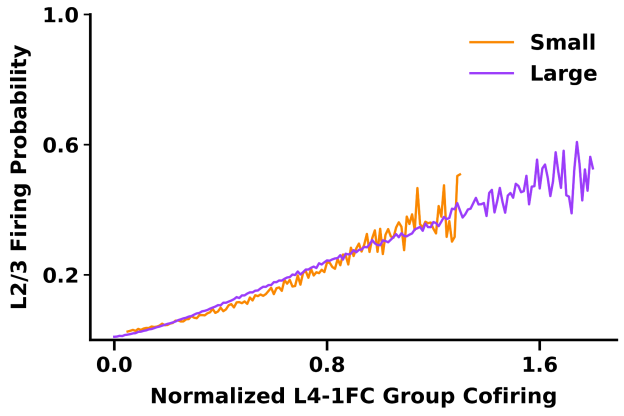


**N**

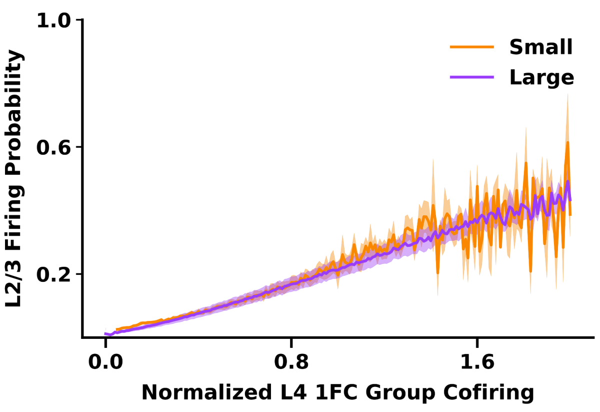


**Supplementary Fig. 53 | Stimulus-defined sL4-1FC ensemble cofiring predicts L2/3 responses with the same nonlinear structure as during spontaneous activity (at rest). A.** Example L2/3 pyramidal neurons showing calcium-event probability as a function of sL4-1FC cofiring (red; SEM shaded). Black lines represent ReLU fits. Null responses (gray dashed lines) remain approximately flat, and end earlier as randomly selected groups (controls) tend to have much fewer high-order cofiring events. **B.** ReLU fits yield lower normalized root mean square error (RMSE) compared to linear fits, confirming nonlinear ReLU-like input–output relationships. **C.** Histogram of goodness-of-fit ($R^2$) values of ReLU fits applied to the response functions (black line fits in A). Note that the majority of ReLU fits are excellent ($R^2 > 0.8$). **D–F.** ReLU-derived parameters: slope (Δ firing probability per 10 cofiring events), extrapolated cofiring required for firing probability = 1, and maximal firing probability. Gray distributions (Null) represent controls. **G–I.** Comparison of parameters obtained under spontaneous (black histogram) versus stimulus (red histogram) conditions shows trends are preserved under stimulation. **J-K.** For an example neuron, L2/3 responses emerge only for large, rare sL4-1FC events: _as we observed under spontaneous stimulation conditions,_ firing rises sharply near the right tail of the cofiring distribution, revealing two regimes (weak/flat and steep). **L.** The distributions of the cofiring cumulative distribution function values corresponding to the response "knee" (intersection of regimes) for all L2/3 neuron responses, under spontaneous (black) versus stimulus (red) conditions are not significantly different. On average, this transition occurs at CDF ≈ 0.94±0.01 during stimulation (corresponding to ~14%±2.0% of the sL4-1FC group cofiring) versus CDF ≈ 0.93±0.02 during spontaneous activity (~13.5%±2.3% of the rL4-1FC group cofiring). To reliably extract response parameters, only neurons with ReLU $R^2 \geq 0.8$ and peak firing probability ≥ 0.6 were included in that analysis. Neurons with response slope <1 were also excluded (2% of cells). **M-N. Normalization of L4 ensemble cofiring collapses L2/3 response functions under stimulus presentation. M.** Average firing probability of L2/3 pyramidal neurons during stimulus presentation plotted as a function of normalized sL4-1FC cofiring for an example mouse. Neurons are grouped by sL4-1FC size quartiles: smallest (Q0–Q25, orange) and largest (Q75–Q100, purple). Cofiring was normalized by dividing by the sL4-1FC group size raised to a constant exponent, yielding overlapping response slopes across quartiles. The exponent used matches that estimated for the same mouse under spontaneous conditions (see manuscript, Fig. 4B). **N.** Same analysis pooled across mice, demonstrating consistent collapse of response functions across sL4-1FC group sizes. Across animals, the normalization exponent ranged from ~0.6–0.7 (see Methods). Insets show mean ± SD across mice; error bars denote SEM (n=5). P-values: "**"< 0.01, "***" < 0.001, "n.s.": non statistically significant. The highest p-value obtained from the permutation of means, the Welch's t-test, and the one-way ANOVA F-test is considered for the level-of-significance.

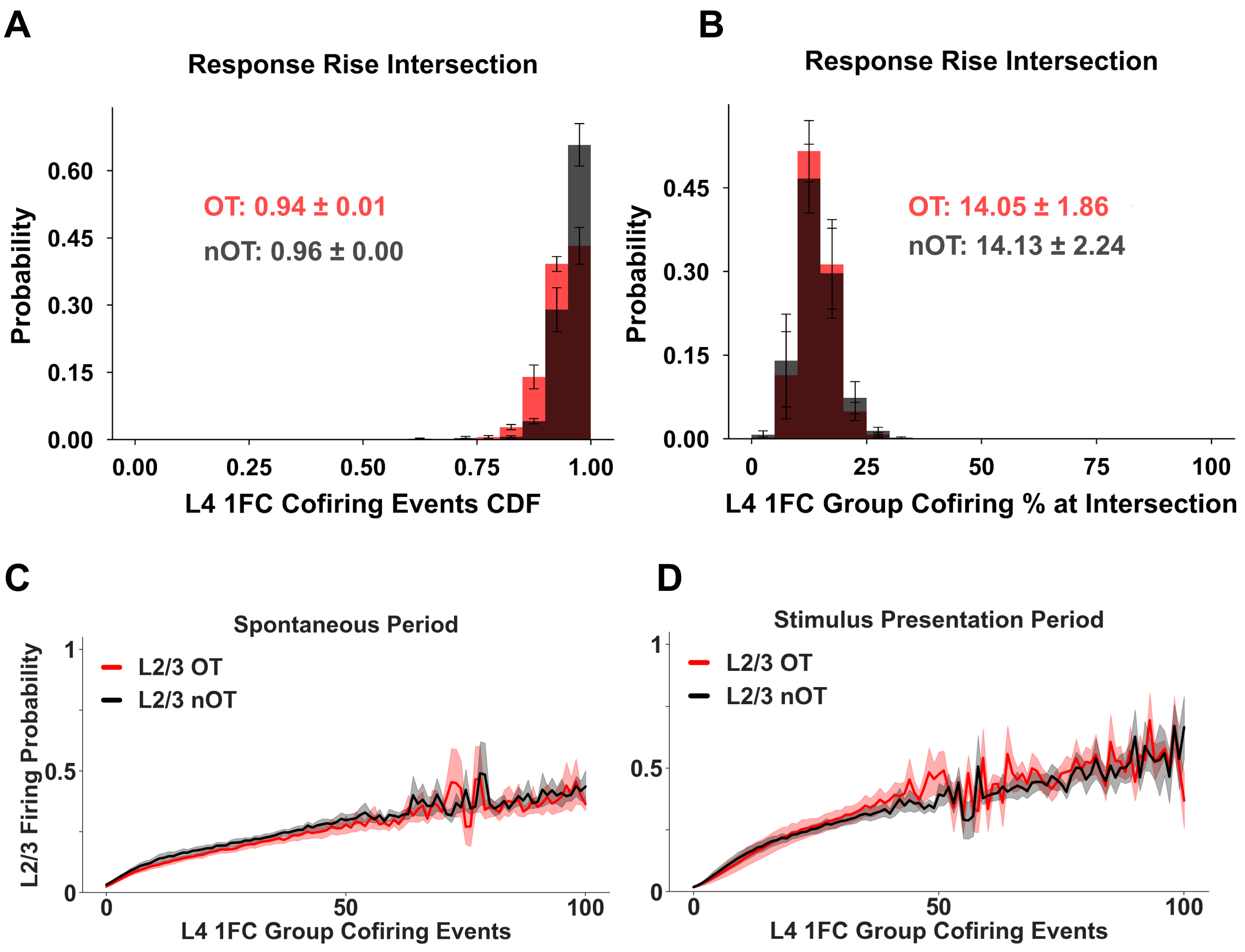


**Supplementary Fig. 54 | A-B) Linear-regime transition point is similar for OT and nOT L2/3 neurons during stimulation. (A)** Histogram of the L4 1FC co-firing CDF value at the transition point between two-piece linear fits of the stimulus-epoch input–output function for orientation-tuned (OT; red) versus non–orientation-tuned (nOT; black) L2/3 neurons. **(B)** Group means of the transition point expressed as percentile of cofiring L4-1FC group neurons for L2/3 OT versus nOT neurons; both populations show comparable transition points. Insets report mean ± SD across mice (n=5); error bars denote SEM. **C-D) The impact of the orientation tuning on L2/3 neuron response functions. (C)** The average response of L2/3 orientation-tuned (OT) neurons (red) plotted as a function of the number of cofiring events in their of **r**L4-1FC groups during *spontaneous conditions*. Black line plots the average response of L2/3 neurons that are not orientation-tuned. **(D)** As in (C) for the stimulus presentation period. Considering L4-1FC groups corresponding to orientation-tuned L2/3 neurons versus L4-1FC groups corresponding to non-orientation-tuned neurons makes no difference in the functional form of the average response function. Shaded regions: SEM across mice (n=5).

A B C D

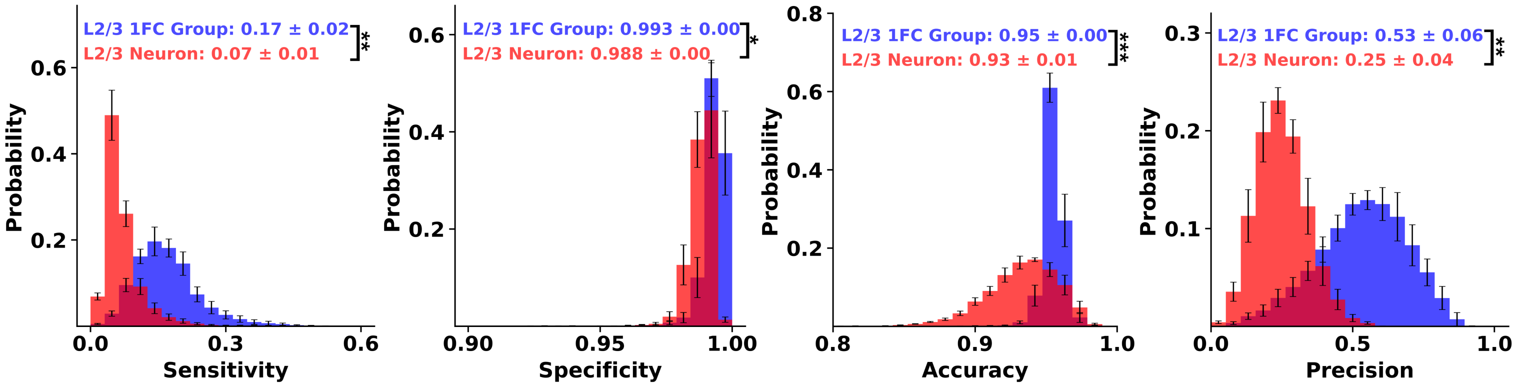


**Supplementary Fig. 55| Ensemble-to-ensemble outperforms ensemble-to-neuron activity transfer during stimulus presentation. A.** Histogram of the sensitivity of activation transfer from the stimulus-defined *sL4-1FC groups* of L2/3 neurons to the corresponding *sL2/3-1FC groups* (blue) versus to the *individual L2/3* "index" neuron (red). Sensitivity is defined as the fraction of frames in which the sL4-1FC group is active given that the L2/3 target (group or neuron) is active. *Active frames* are defined as imaging frames in which the group cofiring z-score is above 3 for the L4-1FC group and 2 for the L2/3-1FC group. The latter ensured the number of frames in which rL2/3-1FC groups are active is similar on average to the number of events occurring in individual L2/3-neurons, allowing for a fair comparison. Ensemble-to-ensemble transmission shows markedly higher sensitivity. **B.** Specificity, defined as the fraction of frames in which the sL4-1FC group is inactive given that the L2/3 target is inactive, is likewise higher for ensemble-to-ensemble transmission. **C.** Accuracy, defined as the fraction of frames in which L4 and L2/3 targets are concordant (both active or both inactive). **D.** Precision, defined as the fraction of frames in which the L2/3 target is active given that the sL4-1FC group is active. *Note that across all metrics, ensemble-to-ensemble (sL4-1FC → sL2/3-1FC) communication outperforms ensemble-to-neuron coupling, indicating that interlaminar information transfer under stimulation is more reliably captured at the ensemble level.* This is consistent with what we observed under spontaneous activity (Suppl. Fig. 41). Insets report mean ± SD across mice (n=5); error bars denote SEM. P-values: "*"< 0.05, "**"< 0.01,"***"< 0.001; "n.s.": non statistically significant. The highest p-value obtained from the permutation of means, the Welch's t-test, and the one-way ANOVA F-test is reported as the level-of-significance. Note that the results we report here are robust to an extensive sensitivity analysis performed over a *range of different thresholds*.

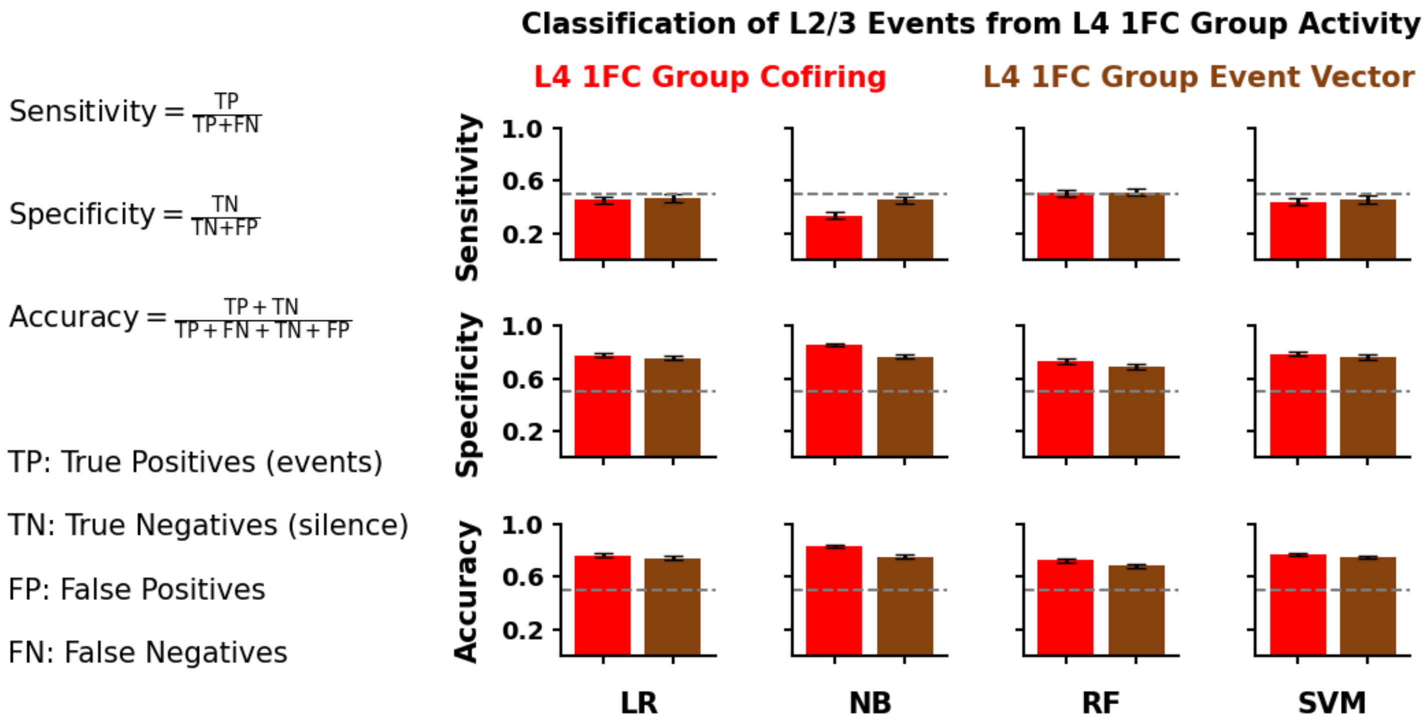


**Supplementary Fig. 56 | Aggregate sL4-1FC ensemble cofiring predicts L2/3 firing under stimulus presentation conditions.** We trained four classifiers—logistic regression (LR), Gaussian naïve Bayes (NB), random forest (RF), and support vector machine (SVM)—to predict L2/3 neuron firing during stimulus presentation per imaging frame, using two input representations: **(i)** the number of cofiring events within the neuron's sL4-1FC group (red), and **(ii)** the identity-resolved sL4-1FC activity vector, which corresponds to the binary firing pattern across all L4-1FC group members (brownblack). Performance, reported as sensitivity, specificity and accuracy (see Methods), was evaluated with a 70/30 train/test split of each one-hour long recording: 1FC groups were computed from the first 40 min (training epoch) and performance was tested on the last 20 minutes (test epoch). Analysis included neurons with L4-1FC size > 15, the training set was class-balanced (firing versus no firing frames) by subsampling non-firing frames. Error bars show SEM across mice (n=5).

#### 13. Stimulus Prediction as a Function of the 1FC Group Cofirings

This section treats **stimulus orientation** as a candidate interlaminar signal and tests how **L4–L2/3 ensembles** support its transmission. We compare ensembles defined by L4-1FC and L2/3-1FC groups identified during rest or stimulus presentation, including their orientation-tuned subsets, against size-matched random controls. For each orientation-tuned ("index") L2/3 neuron, we quantify the probability of *orientation mismatch* between the presented stimulus and the neuron's preferred orientation as a function of the stimulus-evoked co-firing count within the associated L4-1FC ensemble. We then compute *normalized mutual information (nMI)* between stimulus orientation and (i) single-neuron eventograms, (ii) sL4-1FC and sL2/3-1FC ensemble co-firing, and (iii) joint neuron–ensemble activity. Finally, we estimate the fraction of L2/3 neurons exhibiting significant within-frame ensemble co-firing ("active" ensembles) and evaluate stimulus predictability from L4-1FC and L2/3-1FC activity using *accuracy, precision, sensitivity, and specificity.*

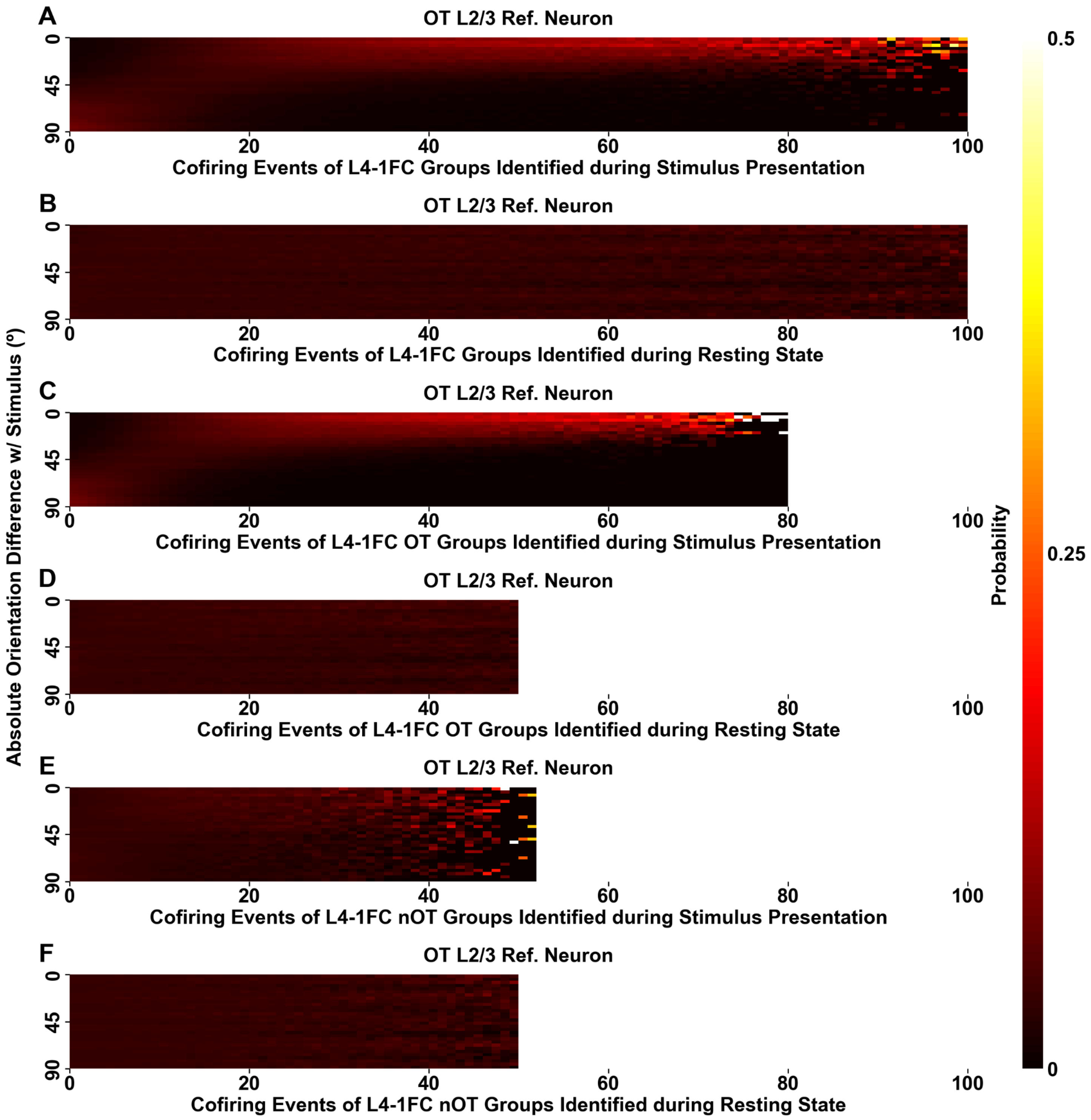


**Fig. 57 | Stimulus orientation is better predicted by stimulus-defined (sL4-1FC) group co-firing than by resting-state–defined (rL4-1FC) co-firing.** Heat maps show, across L2/3 index neurons, the conditional distribution of the absolute orientation mismatch |Δθ| between the presented stimulus and each neuron's preferred orientation as a function of the within-frame co-firing count in its L4-1FC group; color denotes probability per |Δθ| .

**(A)** Orientation-tuned (OT) L2/3 neurons only, using sL4-1FC groups. **(B)** Same as (A), but using rL4-1FC groups identified during spontaneous activity. **(C-D)** Same as (A-B), restricting co-firing to OT neurons within each L4-1FC group. **(E-F)** Same as (A-B), restricting co-firing to nOT neurons within each sL4-1FC group. Similar, slightly weaker trends are obtained when replacing L4-1FC with the corresponding L2/3-1FC group (not shown).

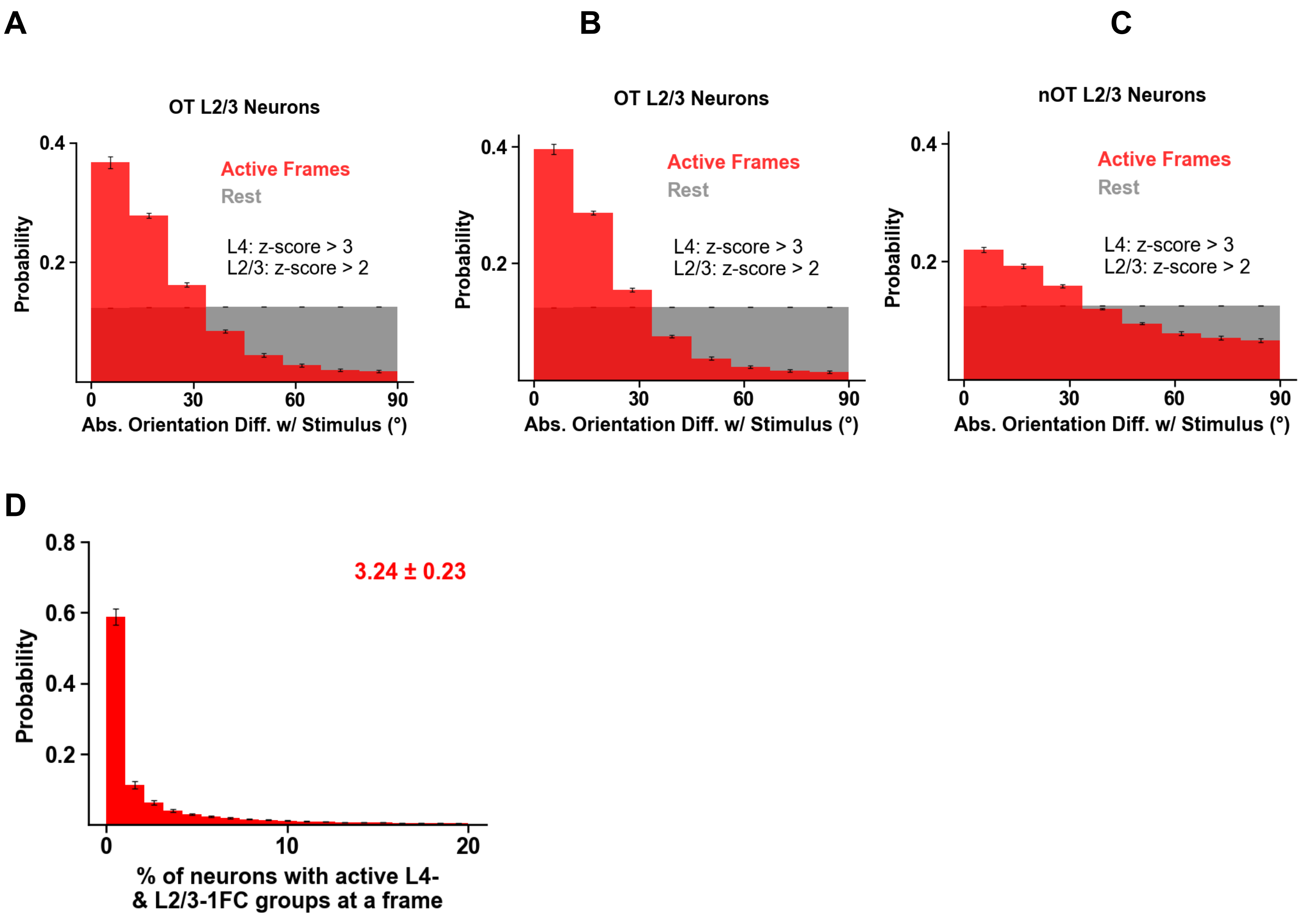


**Supplementary Figure 58 | Ensemble co-firing preferentially occurs at frames when the stimulus presented is "near" the index neuron's preferred orientation. A-C)** Histograms show the absolute orientation mismatch |Δθ| between the presented stimulus and each L2/3 index neuron's preferred orientation, evaluated at selected ("active") frames. *Red:* Active frames. A frame is defined as *active,* when both the sL4-1FC and sL2/3-1FC groups have *significant cofiring* (z-score > 3 and z-score > 2, respectively; note that threshold sensitivity analysis preserves the results). *Gray:* Null frames drawn from the same recording that did not meet the activity criterion. **(A)** Active frames for orientation-tuned (OT) L2/3 neurons are enriched for small |Δθ| relative to null, consistent with Suppl. Fig. 13.1. **(B)** Same as (A), further requiring that the *OT L2/3 index neuron fired* for a frame to be considered *active* (subset of A); as expected the enrichment persists and the resulting red histogram is very similar to (A). **(C)** Same as (A) for non–orientation-tuned (nOT) L2/3 neurons; here the enrichment is lost. **D) Sparse ensemble activation.** Histogram of the fraction of L2/3 neurons per imaging frame whose associated sL4-1FC and sL2/3-1FC groups simultaneously exceeded the co-firing criterion (≥94rd percentile) that indicates the approximate mean onset of their response regime (see Fig 6C). Frames in which there were no neurons that had both sL4-1FC and sL2/3-1FC ensembles active were excluded (≈12% of all frames). Insets show mean ± SD of mouse means; error bars, SEM across mice (n = 5).

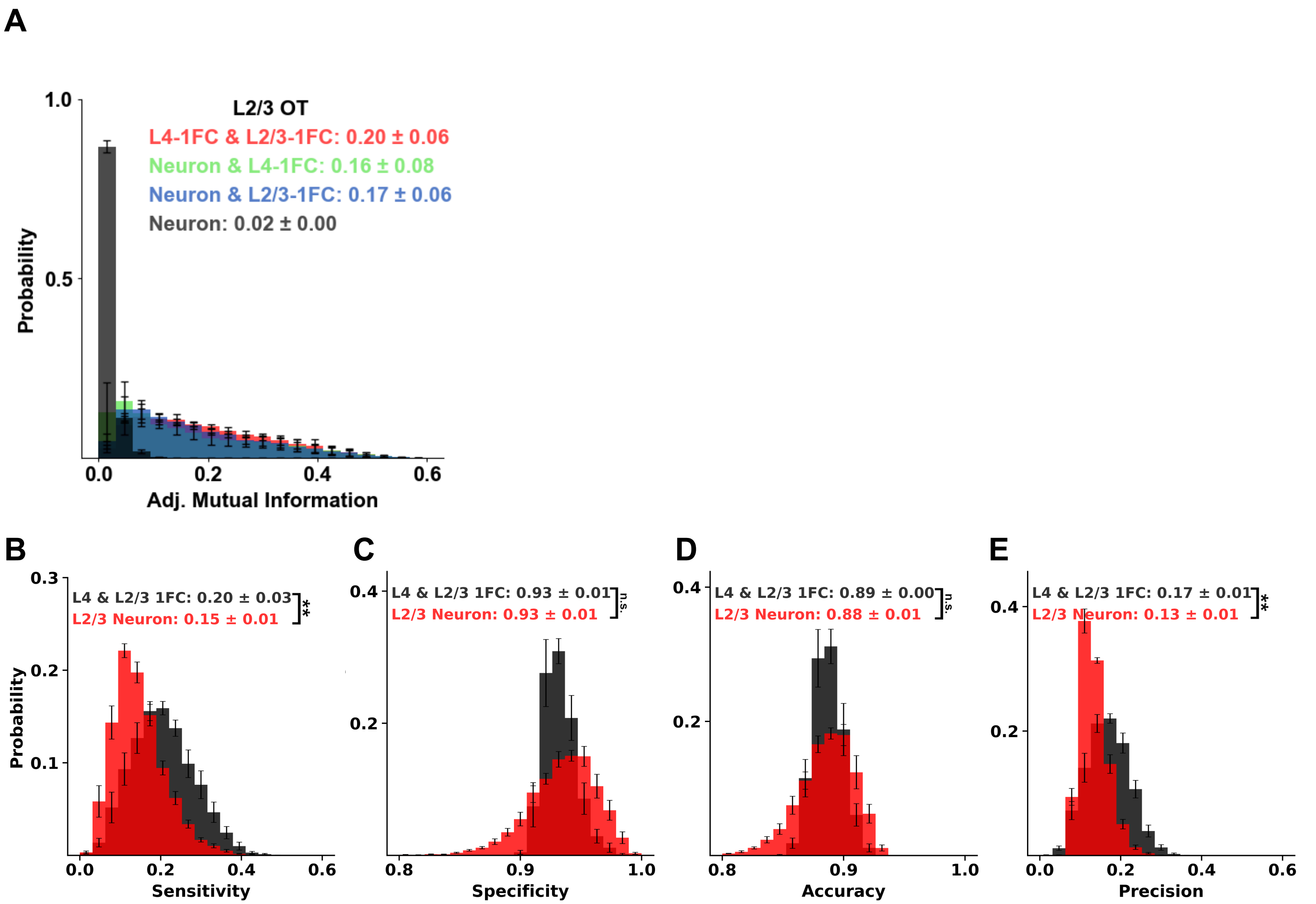


**Supplementary Figure 59 | Ensemble co-firing captures more information about the stimulus than single-neuron activity alone. A.** Distributions of adjusted mutual information between stimulus direction and activity of L2/3 orientation-tuned (OT) neurons. _Black:_ nMI between stimulus direction and each L2/3 orientation tuned (OT) neuron's calcium eventogram. _Green/Blue:_ normalized *joint* MI (jMI) between stimulus direction and the joint activity of the L2/3 OT neuron plus the co-firing in its sL4-1FC group (green) or sL2/3-1FC group (blue). _Red:_ jMI between stimulus direction and the joint activity of the L2/3 OT neuron together with both its sL4-1FC and sL2/3-1FC group co-firing. Joint representations convey more stimulus information than the neuron alone; sL4- versus sL2/3-1FC co-firing yields similar njMI, whereas combining both groups provides the highest njMI. **B-E.** Ensemble co-activation outperforms single-neuron firing for identifying preferred stimulus orientation. A *binary* target labeled frames in which the presented stimulus matched the preferred direction/orientation of the L2/3 index neuron (Methods). We compared two predictors: (i) significant synchronous co-firing of the index neuron's sL4-1FC and sL2/3-1FC groups (black) and (ii) firing of the index L2/3 neuron alone (red). Distributions across mice (n = 5) are shown for sensitivity (B), specificity (C), accuracy (D), and precision (E). Ensemble co-activation increases sensitivity and precision relative to single-neuron firing, while specificity and accuracy have comparable means but substantially reduced variance. Insets show mean ± SD of mouse means; error bars, SEM across mice (n = 5). Significance: permutation of means, Welch's t test, and ANOVA F test (largest P reported); **:$P < 0.01$, ***:$P < 0.001$, n.s.: not significant.
